# Catalytic Redundancies and Conformational Plasticity Drives Selectivity and Promiscuity in Quorum Quenching Lactonases

**DOI:** 10.1101/2024.05.01.592096

**Authors:** Marina Corbella, Joe Bravo, Andrey O. Demkiv, Ana Rita Calixto, Kitty Sompiyachoke, Celine Bergonzi, Mikael H. Elias, Shina Caroline Lynn Kamerlin

**Author notes:** Univ. Grenoble Alpes, CNRS, CEA, IBS, F-38000 Grenoble. Correspondence: Shina Caroline Lynn Kamerlin, Mikael Elias.

## Abstract

Several enzymes from the metallo-β-lactamase-like family of lactonases (MLLs) degrade *N-* acyl-L-homoserine lactones (AHLs). In doing so, they play a role in a microbial communication system, quorum sensing, which contributes to pathogenicity and biofilm formation. There is currently great interest in designing quorum quenching (*QQ*) enzymes that can interfere with this communication and be used in a range of industrial and biomedical applications. However, tailoring these enzymes for specific targets requires a thorough understanding of their mechanisms and the physicochemical properties that determine their substrate specificities. We present here a detailed biochemical, computational, and structural study of the MLL GcL, which is highly proficient, thermostable, and has broad substrate specificity. Strikingly, we show that GcL does not only accept a broad range of substrates but is also capable of utilizing different reaction mechanisms that are differentially used in function of the substrate structure or the remodeling of the active site *via* mutations. Comparison of GcL to other lactonases such as AiiA and AaL demonstrates similar mechanistic promiscuity, suggesting this is a shared feature across lactonases in this enzyme family. Mechanistic promiscuity has previously been observed in the lactonase/paraoxonase PON1, as well as with protein tyrosine phosphatases that operate *via* a dual general-acid mechanism. The apparent prevalence of this phenomenon is significant from both a biochemical and an engineering perspective: in addition to optimizing for specific substrates, it is possible to optimize for specific mechanisms, opening new doors not just for the design of novel quorum quenching enzymes, but also of other mechanistically promiscuous enzymes.

## Introduction

The molecular determinants for the high proficiency and specificity of enzymes are often discussed. However, while the chemical role of key active-site and catalytic residues is typically depicted as uniquely specific, recent examples of enzymatic promiscuity demonstrate that the same active-site residues may perform different roles in the same enzyme, allowing for both substrate and catalytic promiscuity.^1–4^ On the one hand, different subsets of active-site residues in a large binding pocket can be used to facilitate reactivity with different substrates.^4^ Conversely, those same active-site residues may also be capable of performing multiple tasks within the same active site to catalyze the same reaction, when the pre-organization of reactive residues allows for several, energetically close, reaction trajectories. These two scenarios are not mutually exclusive. For example, both scenarios have been observed with the enzyme serum paraoxonase 1 (PON1), a catalytically promiscuous organophosphatase/lactonase.^4, 5^ In this work, we focus on providing a detailed structural and mechanistic description of the multi-functional role of active site residues in different members of a catalytically and mechanistically promiscuous family of enzymes.

Our work discusses lactonases (EC 3.1.1.81) from the metallo-β-lactamase-like family of lactonases (MLLs). MLLs degrade *N*-acyl-L-homoserine lactones (AHLs), molecules that are used in a microbial communication system called quorum sensing (QS) to coordinate a variety of behaviors, including virulence and biofilm formation.^6, 7^ By degrading AHLs, these enzymes can interfere with microbial signaling and are therefore called quorum quenchers (QQ). They have previously been reported to inhibit behaviors that are regulated by bacterial QS, such as virulence factor production and biofilm formation, and can also alter microbiome population structure.^8–13^ As a result, the mechanisms of QQ enzymes, as well as their targeted biotechnological applications, are currently topics of intensive research. Furthermore, designer quorum quenching enzymes are being developed for a host of biotechnologically important processes, such as preventing biofouling and biocorrosion, and for use in biomedical applications.^10, 14–18^

Lactonases have been identified from a wide range of organisms, including archaea, bacteria, fungi, and mammals.^18–25^ Three main families of lactonases have been identified,^20^ all of which are metalloenzymes.^26–28^ Paraoxonases (PONs), primarily isolated from mammals, exhibit a six-bladed β-propeller fold, and a mono-metallic (calcium) active site center.^20^ The phosphotriesterase-like lactonases (PLLs) exhibit an (α/β)_8_-fold and a bi-metallic active site center.^29^ A third family of lactonases, which are the focus of the present work, are the metallo-β-lactamase-like lactonases (MLLs).^20^ These enzymes possess an αββα-fold and a conserved dinuclear metal-binding motif, HxHxDH, which is involved in the coordination of the bi-metallic active site center. Numerous representative enzymes from this family have been kinetically and/or structurally characterized, including AiiA,^30^ AiiB,^31^ AidC,^32^ MomL,^33^ and AaL.^34^

Efficient engineering of lactonases for biotechnological applications, through approaches such as the construction of focused libraries for directed evolution, would benefit from a detailed understanding of the mechanism and selectivity of these enzymes towards specific lactones. This is particularly important as the selectivity of these enzymes towards different substrates is complex and can depend on, for example, the acyl chain lengths of the different AHLs.^30, 33, 35, 36^ In addition, the chemical structure of the lactone modulates the specificity of signaling.^37–39^ Despite the importance of this question, the catalytic mechanism of lactonases (**Figure 1**) is not yet fully understood. In MLLs and PLLs, the bi-metallic center is hypothesized to activate the substrate and a catalytic water molecule.^20^ The nature of this nucleophilic water molecule is yet unclear: while the nucleophile is often hypothesized to be the metal-bridging water molecule in the form of a hydroxide ion (**Figure 1A**),^20^ compelling evidence for this mechanism has been elusive due to the difficulty of isolating transition states in crystal structures.

**Figure 1.**
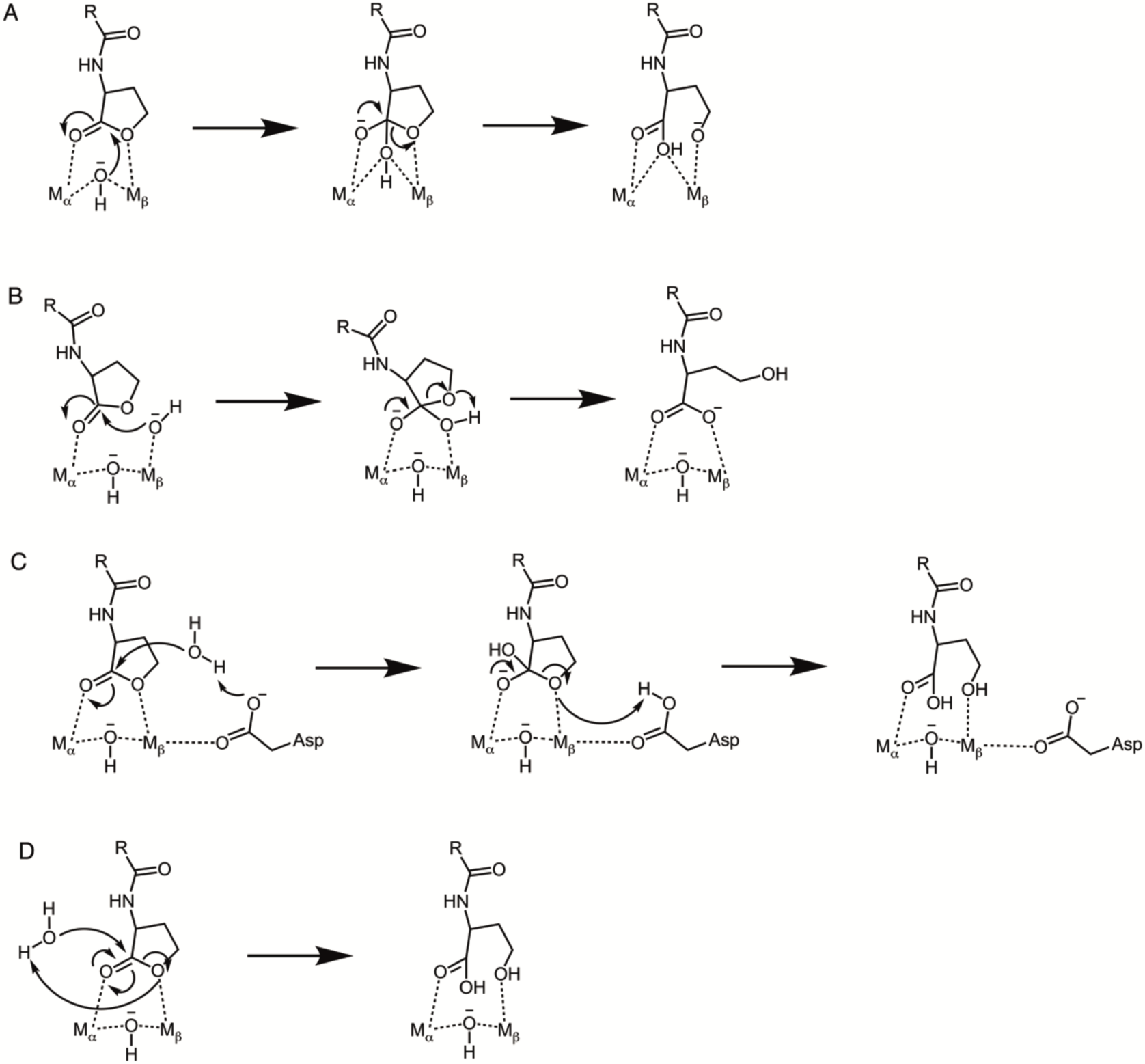
Plausible mechanisms for the hydrolysis of *N*-acyl-L-homoserine lactones by lactonases. (**A**) Stepwise nucleophilic attack on the carbonyl carbon of the lactone ring by the bridging hydroxide ion, followed by breakdown of the resulting tetrahedral intermediate (“bridging hydroxide mechanism”). (**B**) Stepwise nucleophilic attack on the carbonyl carbon of the lactone ring by a terminal hydroxide ion, followed by breakdown of the resulting tetrahedral intermediate (“terminal hydroxide mechanism”). (**C**) Stepwise general-base catalyzed mechanism, in which the side chain of a metal bound aspartic acid (Asp122 using GcL numbering) acts as a general base to activate the nucleophilic water molecule followed by the breakdown of the resulting tetrahedral intermediate (“Asp mechanism”). (**D**) Concerted nucleophilic attack on the carbonyl carbon of the lactone ring by an active site water molecule activated by proton transfer to the lactone ring oxygen, and opening of the lactone ring (“concerted mechanism”). Note that, for simplicity, we have shown the ring oxygen in the product state of mechanisms **A** and **B** to be deprotonated; however, the ring-opening reaction would benefit from protonation by an acid catalyst, the precise identity of which can vary depending on the system. The shorthand designations for each mechanism, shown in parenthesis, will be used throughout the text.

A near-identical mechanism to that shown for lactone hydrolysis in **Figure 1A** (“bridging hydroxide mechanism”) has been proposed for organophosphate hydrolysis by phosphotriesterases (PTEs),^40–42^ a closely related enzyme family to Phosphotriesterase-like Lactonases (PLLs). PTEs possess a similar bimetallic active site; however, more recent experimental evidence has suggested that rather than the bridging hydroxide ion, the nucleophile is more likely to be a terminal hydroxide ion (**Figure 1B**, “terminal hydroxide mechanism”).^43^ This is in agreement with data from studies of designed binuclear catalysts of phosphate hydrolysis reactions,^44^ computational studies of a related enzyme, methyl parathion hydrolase,^45^ as well as other metallophosphatases.^46, 47^ We note again here that a terminal hydroxide ion would be expected to have a more favorable p*K*_a_ for nucleophilic attack than the bridging hydroxide ion, the p*K*_a_ of which would be substantially depressed by coordination to two metal ions (in the range of 9-10 for the terminal hydroxide ion,^48–50^ depending on metal ion, compared to 7.3 for the bridging hydroxide ion in the case of the analogous enzyme phosphotriesterase^51^). The terminal hydroxide ion would also have more structural flexibility than the bridging hydroxide ion, which is held tightly in place by the two metal ions it coordinates. Moreover, it is possible that the nucleophile is not metal coordinated at all, but rather is an active site water molecule activated, for example, by general-base catalysis through a metal-bound aspartic acid in the active site*, e.g.* Asp122 (**Figure 1C**, “Asp-mechanism”), or by concerted proton transfer to the lactone ring oxygen (**Figure 1D**, “concerted mechanism”). Additionally, the hydrolysis of the lactone ring involves a leaving alcoholate group. This poor leaving group may benefit from protonation by an acid catalyst. It has been proposed that this protonation is carried out by a metal-coordinating aspartic acid residue in the case of the lactonase AiiA.^30^ In a mechanism such as that shown in **Figure 1C**, the protonated aspartic acid generated upon nucleophilic attack to open the lactone ring could then act as a general acid to assist in leaving group departure. When taking into account the potentially short lifetime of the tetrahedral intermediate formed upon lactone ring opening (*i.e.* the reaction can potentially proceed *via* a borderline mechanism that is almost concerted in nature^52^), then the number of potential viable mechanisms becomes a large combinational problem, and distinguishing between these different mechanisms does not seem experimentally possible. The full span of potential mechanisms has not been previously considered for either lactonases or organophosphate hydrolases with analogous active sites. Simulation approaches are ideal to sample these different mechanisms and directly compare different mechanisms with a range of lactone substrates with different acyl tail lengths. Obtaining deeper insight into possible mechanisms across a range of substrates will also provide insight into plausible catalytic mechanisms for other metallohydrolases that have similar active site architectures.

Here we aim to resolve the catalytic mechanism(s) of lactonases from the metallo-β-lactamase superfamily. We provide the structural and biochemical analysis of the lactonase GcL (WP_017434252.1), isolated from the thermophilic bacteria *Parageobacillus caldoxylosilyticus*.^1,2^ GcL is a thermostable, highly proficient lactonase with broad substrate specificity (*k*_cat_/*K*_M_ values range between 10^4^ to 10^6^ M^−1^ s^−1^) for substrates such as *N-*butyryl (C4) L-homoserine lactone (HSL) and *N*-decanoyl (C10)-HSL. We combine unique structural data, including the structure of GcL in complex with an intact substrate (*N*-hexanoyl-HSL (C6-HSL)) and a hydrolytic product (hydrolysis of *N*-octanoyl-HSL (C8-HSL)), with empirical valence bond (EVB) simulations^53^ to probe possible catalytic mechanisms for lactone hydrolysis. The results of computational modeling were tested against mutational data in GcL and extended to other lactonases from the metallo-β-lactamase superfamily, including AaL and the more distantly related AiiA. Results reveal that the enzymatic reaction can proceed *via* (at least) two chemically distinct but energetically similar mechanisms, with the precise pathway taken being dependent on the specific substrate and/or enzyme variant. This catalytic versatility, making use of a distinct subset of active site residues, is consistent with the reported enzymatic promiscuity, and broad selectivities of lactonases.

Mechanistic promiscuity has previously been suggested for an analogous lactonase, PON1, which was suggested to both possess catalytic backups in the active site,^4^ as well as have more than one mechanism being viable at the same time.^5^ Similarly, several protein tyrosine phosphatases appear to operate *via* a dual general acid mechanism.^54–56^ Such mechanistic promiscuity has not been observed in the literature as a widespread phenomenon, however, the data presented here suggests that it is, at minimum, common to multiple distinct quorum quenching lactonases. This is significant not just from a biochemical standpoint but also from an engineering perspective: protein engineering studies often focus on optimizing activity for specific substrates and while doing so it can optimize for a specific *reaction mechanism* out of a pool of viable mechanisms.^5^ Here, we extend the potential generalizability of that concept across several enzymes from a different family of lactonases. This opens the door to new strategies in the design and engineering of not just biotechnologically important quorum quenching enzymes, but also other enzymes that may be mechanistically promiscuous.

## Results and Discussion

### The Structure of GcL in Complex with L-Homoserine Lactone

The structure of GcL was previously determined and described in refs. ^1,2^. GcL possesses a bi-metallic active site center, which is comprised of iron and cobalt cations. The binuclear center, common to all metallo-β-lactamase-like lactonases, is coordinated by five histidine residues and two aspartic acid residues.^20^ A water molecule bridges both metal cations, and has previously been hypothesized to be the reaction nucleophile in this family.^20^ The structure of GcL bound to C6-HSL is overall similar to the previously obtained structures of GcL bound to the substrates C4- and 3-oxo-dodecanoyl (3-oxo-C12)-HSL^36^ (**Figure S1**). The lactone ring of the substrate sits on the bi-metallic active site, with the carbonyl oxygen atom of the lactone ring interacting with the cobalt cation (2.6 Å) and the ester oxygen atom interacting with the iron cation (2.2 Å). In combination, these interactions likely increase the electrophilic character of the carbonyl carbon atom. The carbonyl oxygen atom is also hydrogen bonded to the hydroxyl group of Tyr223 (3.0 Å). The *N-*alkyl chain of the C6-HSL is kinked and interacts with residue Ile237 (**Figure 2**).

**Figure 2.**
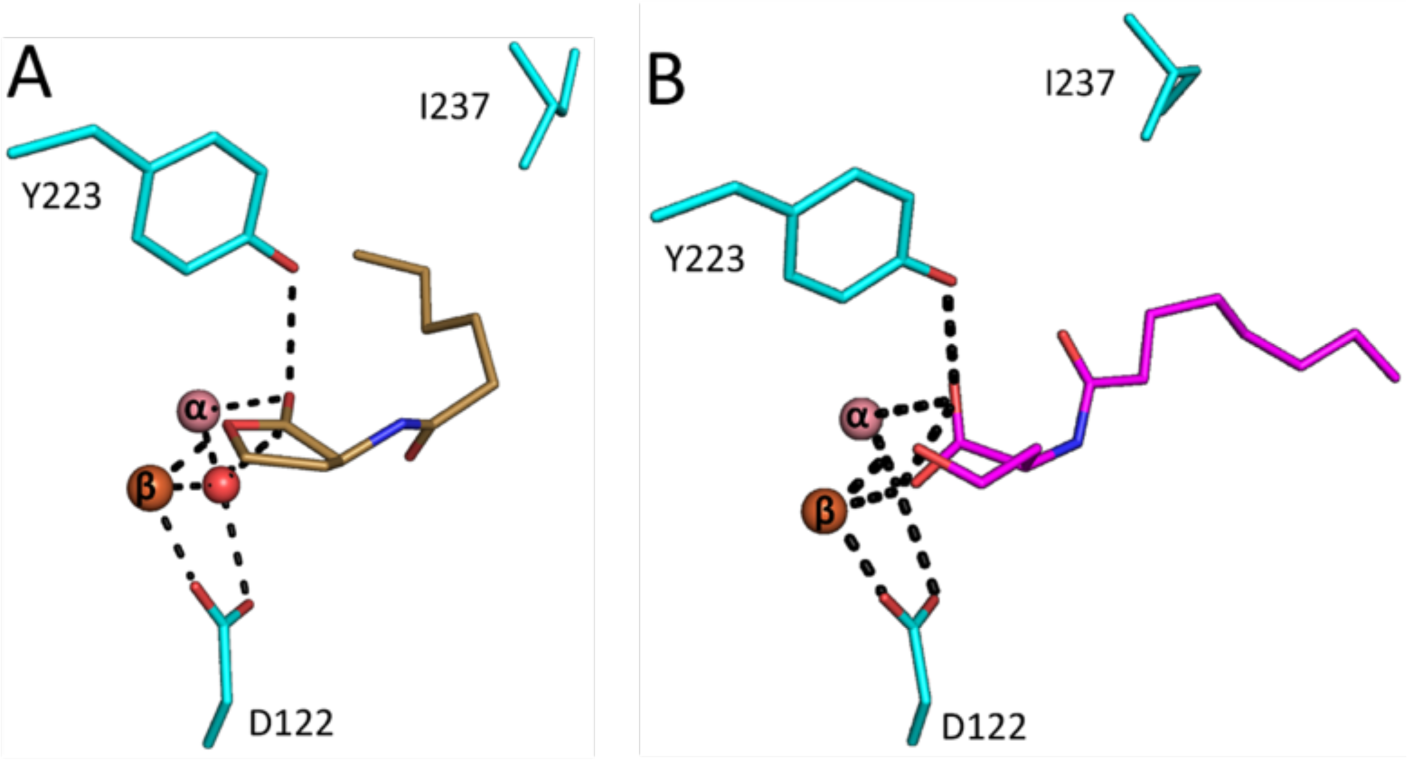
Structures of GcL structures in complex with lactone substrate and hydrolysis products. (**A**) Active site of GcL (cyan sticks) bound to its substrate C6-HSL (9AYT; yellow sticks, modeled at 0.8 occupancy). The lactone ring sits on the bi-metallic active site (pink and orange spheres). (**B**) The structure of GcL in complex with a product of the hydrolysis of C8-HSL (pink sticks; PDB ID: 9B2O; modeled at 0.7 occupancy). Metal cations are shown as pink (cobalt, α) and orange (iron, β) spheres, and are shown at reduced size for clarity.

### Structure of GcL in Complex with the L-Homoserine Lactone Hydrolysis Products

Co-crystallization trials in the presence of the C8-HSL substrate allowed for the resolution of the structure of GcL in complex with AHL-hydrolytic products bound to the active site (**Table S1**). Attempts to co-crystallize GcL with C6-HSL resulted in crystals with low ligand occupancy (∼0.5, not shown). The structure in complex with C8-HSL was modeled with a ligand occupancy of ∼0.7, and while the moderate occupancy limits the accuracy of the model, it provides important insights into the potential catalytic mechanism of GcL (**Figures 2** and **S2**). Specifically, it reveals that the negative charge of the hydrolytic product carboxylate group is stabilized by the bi-metallic active site center, and that the alcohol group interacts with the α-metal (iron; 2.7 Å; monomer L).

In addition, the carboxylate group of the product is positioned between the two metal cations, 2 Å from the cobalt-cation and 2.8 Å from the iron-cation (**Figure 2B**; monomer L). The other oxygen atom of the carboxylate is hydrogen bonded to the hydroxyl group of Tyr223 (2.7 Å). During lactone hydrolysis, the protonation of the leaving alcoholate group may be an important, catalytically limiting step. The position of the alcohol group is 3.3 Å and 3.5 Å from Asp122 and Tyr223, respectively. This suggests that the side chains of Tyr223 and Asp122 are possible acid catalysts for the reaction, although a protonated Asp122 side chain (protonated in the first reaction step, **Figure 1**) would be expected to have a more favorable p*K*_a_ to fulfill the role of an acid catalyst. An alternative hypothesis, also considered here (**Figure 1**), would be an intramolecular protonation mechanism. These mechanisms are very similar (they all involve proton transfer either from an amino acid side chain or a water molecule), making it very hard to distinguish between them experimentally.

### Substituting Acid Catalyst Candidates Tyr223 and Asp122

Tyr223 and Asp122 are the only polar residues in the vicinity of the substrate in the active site of GcL. Remarkably, the presence of a tyrosine residue side chain is a conserved feature of lactonases, including those from different folds.^57^ Asp122 is also conserved, and is involved in the metal cation coordination. The corresponding residue of Asp122 in the lactonase AiiA was previously hypothesized to protonate the leaving group in a structural and mutagenesis study.^30, 58^ We therefore undertook mutagenesis of both Tyr223 and Asp122, and characterized the resulting Tyr223Phe and Asp122Asn GcL variants (**Table 1** and **Figures S3** to **S8**).

**Table 1.**
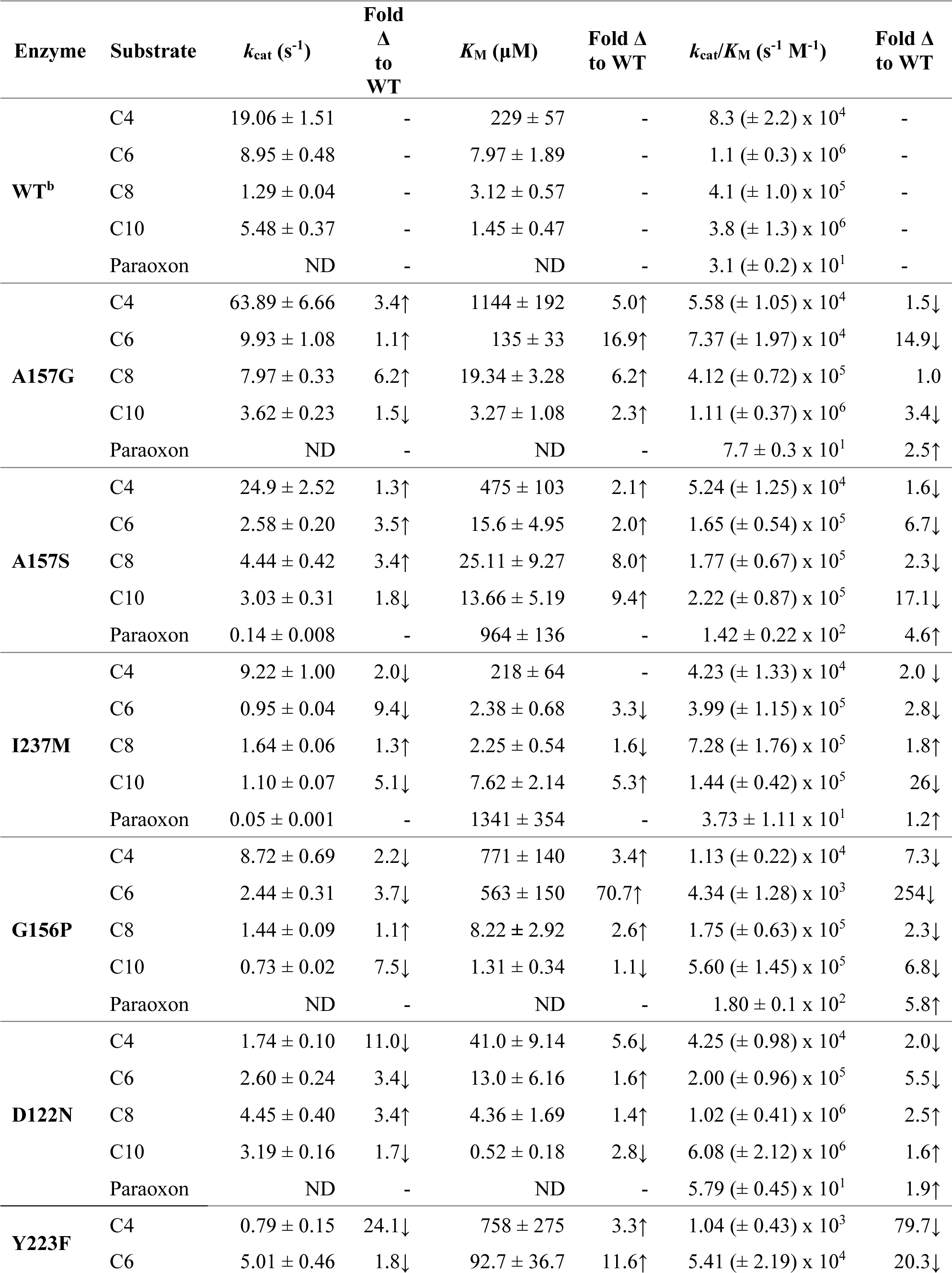
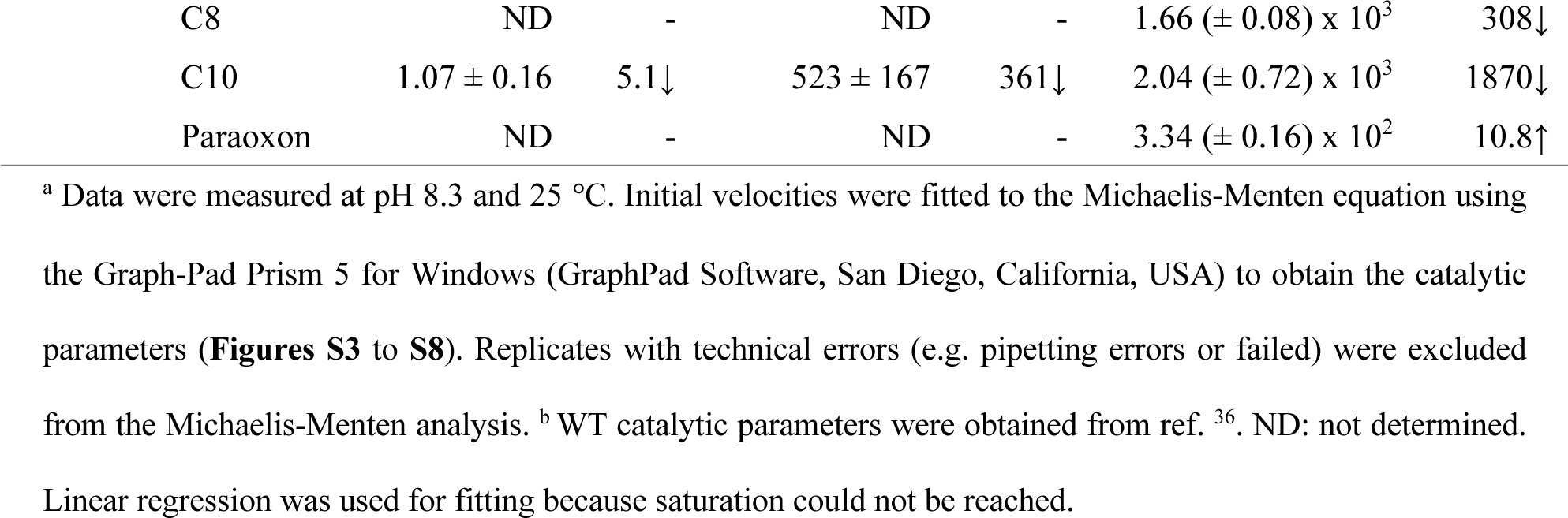
Kinetic parameters for the hydrolysis of AHL substrates and the phosphotriester paraoxon by wild-type GcL and variants.^a^.

The determination of the kinetic parameters for these two mutants reveal that the Asp122Asn substitution causes a drop in catalytic efficiency (∼6-fold; with C6-HSL) and more modest changes with other substrates (C4, C8 and C10-HSL) (**Table 1**). This effect is smaller than those observed for the corresponding mutation made to AiiA (D108N; ∼36-fold reduction of catalytic efficiency with C6-HSL and Co^2+^, see ref. ^30^), yet for both enzymes the mutation does not abolish the lactonase activity. The alterations in the activity of GcL might still suggest that this residue is important but the interpretation of this effect is complicated by the clear role of this residue in metal coordination. This was further evidenced through the capture of multiple conformations through crystallization: one conformation where both metals are present at high occupancy, exhibiting an active site configuration very close to that of the wild-type enzyme (**Figure S9**), and a second form where the active site shows no bound (or with very low occupancy) β-metal (**Figure S9**). This provides evidence that this substitution may destabilize the bimetallic center, possibly by decreasing the affinity of the β-site for metals.

Conversely, Tyr223 is not involved in metal cation coordination, yet, the Tyr223Phe substitution consistently reduces the lactonase activity of the enzyme for all tested AHLs (>2 orders of magnitude reduction in catalytic efficiency for C8- and C10-HSL; **Table 1**). This effect is in the range of the changes described for the equivalent mutation performed in AiiA (Y194F; ∼169-fold reduction of catalytic efficiency with C6-HSL and Co^2+^, see ref. ^30^). In addition, the evaluation of the paraoxonase activity, *i.e.* the promiscuous ability of GcL to hydrolyze this phosphotriester, shows that the Tyr223Phe substitution only impairs lactonase activity: the variant exhibits a ∼11-fold higher paraoxonase activity than the wild-type enzyme. This observation where the same substitution has opposing effects on alternative lactonase and phosphotriesterase activities is not necessarily surprising, as previously observed in our prior work on an analogous enzyme, serum paraoxonase 1 (PON1)^59, 60^. While the impact of this substitution on *k*_cat_/*K*_M_ for the lactonase activity is significant, the impact on the turnover number, *k*_cat_, is much smaller (**Table 1**), suggesting that this side chain is more likely to play an important role in substrate binding or positioning during the chemical reaction as illustrated by its impact on the *K*_M_ values for AHL substrates. Overall, these mutagenesis data suggest that Tyr223 plays an important role in catalysis.

The previously elucidated GcL structure allowed the identification of other key residues lining the active site binding cleft and that may be involved in substrate binding.^36^ Substitutions at these positions that are probing the acyl chain binding crevice produced some variants with changes in kinetic properties. We here report the kinetic characterization for several variants, including Ala157Gly, Ala157Ser, Gly156Pro and Ile237Met (**Table 1** and **Figure S10**). While some of these variants show only modest changes in lactonase activity as compared to the wild-type for several AHL substrates (*e.g.* Ile237Met), they show altered kinetics for specific substrates, resulting in changes in substrate preference. For example, the variant Ala157Ser exhibits a 6.7-fold and 17.1-fold drop in catalytic efficiency against C6- and C10-HSL, respectively. The variant Gly156Pro exhibits larger changes in preference, with a 254-fold drop in catalytic efficiency against C6-HSL. Intriguingly, the drop in substrate preference is not linearly changing with the increase in chain length: it is 2.3 and 6.8-fold for C8- and C10-HSL, respectively. This suggests that subtle substrate conformational sampling may occur differentially as a function of chain length and consequently the acyl chain hydrophobic character and entropy.

To gain some molecular insights into the effects of these substitutions on the active site configuration, structures were solved for both Gly156Pro and Ile237Met variants. The structure of the Gly156Pro variant reveals the significance of the Asn152-Ala157 loop, involved in the accommodation of long chain AHL substrates (**Figure S11**). The conformational change of this loop may be responsible for the altered substrate preference of the enzyme. On the other hand, the Ile237Met variant shows a slightly altered conformation of the Pro234-Asp240 loop, in the vicinity of the amide group of the AHL substrate (**Figure S12**). This conformational change is more distant from the atoms of a long acyl chain AHL substrate, and this is therefore consistent with the minimal changes in catalytic properties recorded for this variant.

### Empirical Valence Bond Simulations of the Hydrolysis of C6-HSL by Wild-Type GcL

As shown in **Figure 1**, lactone hydrolysis by GcL (and other lactonases) can proceed through multiple pathways that are difficult-to-impossible to distinguish between experimentally. As shown in **Figure 1**, these mechanisms can be either stepwise or concerted in nature. They can also involve either a metal-bound bridging or terminal hydroxide ion or a free water molecule as the nucleophile, and, in the case of the Asp mechanism (**Figures 1** and **S2**), can recruit an active site side-chain (Asp122) to act as a general base.

As our starting point, we constructed EVB models for the hydrolysis of C6-HSL by wild-type GcL through any of four possible reaction mechanisms: a bridging hydroxide mechanism (**Figure 1A**), a terminal hydroxide mechanism (**Figure 1B**), an Asp mechanism (**Figure 1C**), and a concerted mechanism (**Figure 1D**). Simulations were initiated from the crystal structure of wild-type GcL in complex with C6-HSL (PDB ID: 9AYT, this study), as described in the **Materials and Methods**. The results of these simulations are summarized in **Figures 3** and **S13**, and **Table 2**.

**Figure 3.**
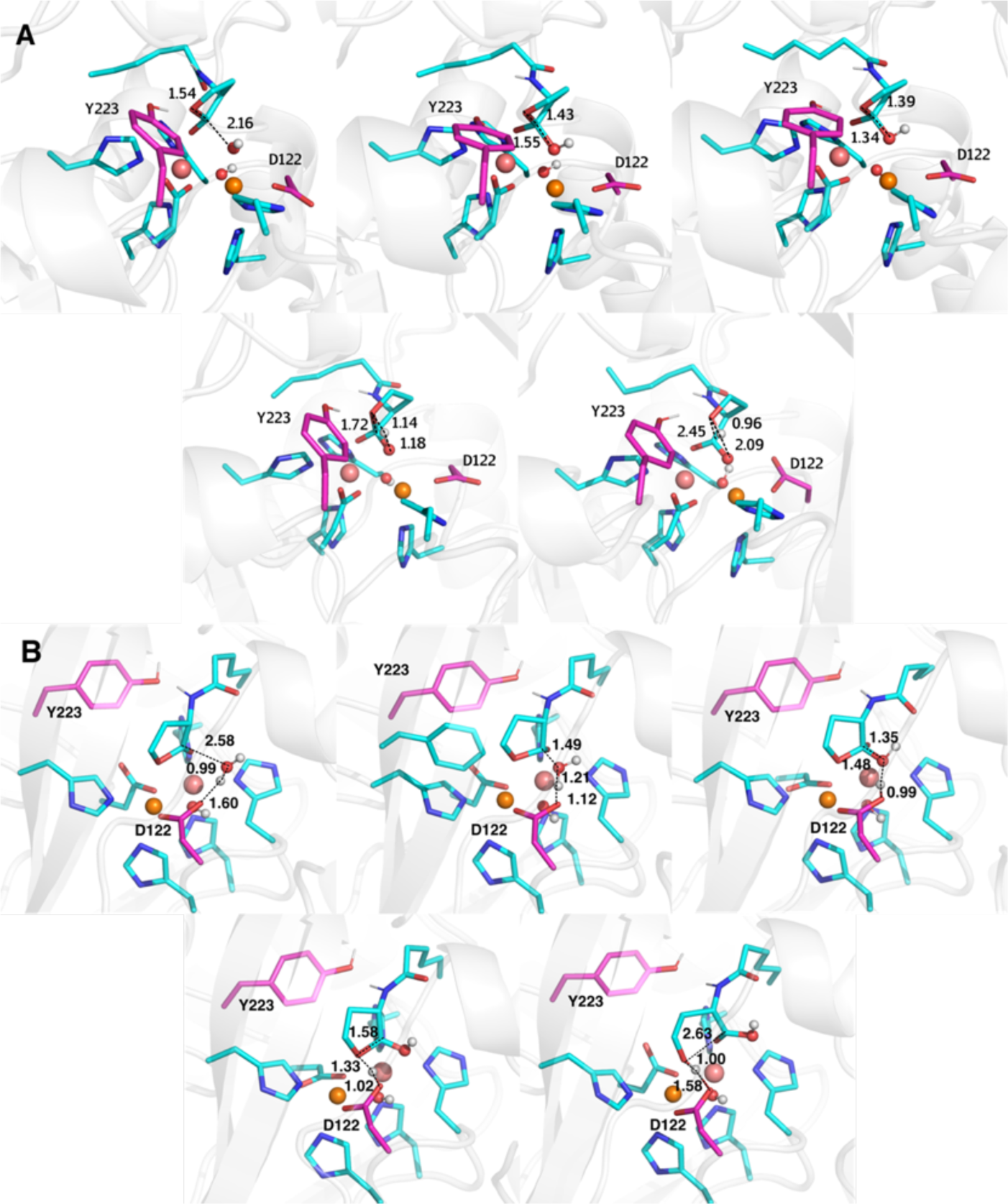
Representative structures of the Michaelis complexes, transition states, and intermediate states, for the hydrolysis of C6-HSL catalyzed by wild-type GcL *via* energetically favorable (**A**) terminal hydroxide and (**B**) Asp mechanisms (**Figures 1B and C** and **Table 2**), as obtained from empirical valence bond simulations of these reactions. The structures shown here are the centroids of the top ranked cluster obtained from clustering on RMSD, performed as described in the **Materials and Methods**. The distances labeled on this figure (Å) are averages at each stationary point over all the EVB trajectories (see **Table S2**, with the corresponding data for the non-enzymatic reaction shown in **Table S3**, and metal-metal distances shown in **Table S4**). Shown here are the substrate, nucleophilic water, bridging hydroxide, Fe^2+^ (brown), Co^2+^ (salmon), and key catalytic residues. The reminder of the protein has been omitted for clarity. The corresponding data for the bridging hydroxide and concerted mechanisms are shown in **Figure S13**.

**Table 2.** A comparison of calculated activation free energies (kcal mol^−1^) for the hydrolysis of C6-HSL catalyzed by wild-type GcL through the different mechanisms considered in this work, compared to an experimental value of 16.2 kcal mol^−1^.^a^

| Mechanism | $\Delta G_1^\ddagger$ | $\Delta G_{\text{int}}$ | $\Delta G_2^\ddagger$ | $\Delta G^0$ |
| --- | --- | --- | --- | --- |
| **Bridging Hydroxide** | 35.7 ± 0.5 | 30.2 ± 0.8 | **36.7 ± 1.1** | 32.6 ± 1.2 |
| **Terminal Hydroxide** | **16.3 ± 0.5** | 11.7 ± 0.6 | 13.8 ± 0.8 | 0.9 ± 1.1 |
| **Asp** | 15.8 ± 0.2 | 11.7 ± 0.5 | **16.2 ± 0.8** | -4.5 ± 0.7 |
| **Concerted** | **21.1 ± 0.4** |  |  | -3.8 ± 0.5 |

Based on our simulations, we obtain very high activation free energies for the bridging hydroxide mechanism (**Figure 1A, Table 2**). This reaction pathway involves the loss of the electrostatically favorable metal-hydroxide interaction as the charge migrates away from the metal ion, resulting in the high activation free energy presented in **Table 2**. We note that nucleophilic attack by the bridging hydroxide on paraoxon has been suggested to be energetically viable based on DFT-based QM cluster or QM/MM calculations in other systems with similar active sites to GcL.^41, 42, 61^ However, interpretation of this data is complicated by the fact that DFT calculations involving hydroxide as a nucleophile tend to significantly underestimate the activation free energies involved,^62–67^ a problem that is likely to be further exacerbated by the presence of the binuclear metal center in the active site, and that no alternate mechanisms were considered in these studies.

In contrast, both the terminal hydroxide and Asp mechanisms (**Figure 1**) appear to be energetically favorable and within a reasonable range of the upper limit of 16.2 kcal mol^−1^ for the experimental value (derived from the turnover number, *k*_cat_, **Table 1**), with indistinguishable activation free energies based on our calculations. In the case of the terminal hydroxide mechanism, this reaction follows a stepwise pathway, involving nucleophilic attack of a terminal hydroxide bound to the β-metal ion on the lactone ring with monodentate coordination to the α-metal ion through the C=O bond, with intramolecular protonation of the intermediate concomitant to ring opening (**Figure 1**). The initial conformation of the lactone necessary to facilitate nucleophilic attack of a terminal hydroxide ion is slightly distorted compared to the putative structure from the crystal structure (**Figure S14**). However, the large active site of GcL could potentially accommodate multiple substrate binding modes. Furthermore, a similar pathway involving a terminal hydroxide ion has been suggested based on both experimental and computational data for a range of analogous systems.^44–47^

In the Asp mechanism (**Figure 1C**), the Asp122 side chain participates in acid-base catalysis during the reaction, first deprotonating the attacking nucleophile, then subsequently protonating the leaving group. A similar mechanism involving a metal-bound aspartic acid side chain has been suggested as a catalytic backup in an analogous lactonase, PON1,^4, 5^ and, by extension, the existence of backup mechanisms is likely to be evolutionarily beneficial in scavenger enzymes. This is also in agreement with the observation that the Asp122Asn substitution shows almost no effect on the turnover number (*k*_cat_) compared to wild-type (**Table 1**), making it likely that a backup mechanism is present. As shown in **Figure S14**, this mechanism is in good agreement with both the crystal structure and the experimental activation free energy of 16.2 kcal mol^−1^ (based on kinetic data presented in **Table 1**).

The final potential mechanism is a concerted mechanism (**Figure 1D**) involving intramolecular proton transfer from the attacking nucleophile, which is an active site water molecule. Although this pathway is less energetically unfavorable than the bridging hydroxide mechanism, it is significantly higher in energy than either the terminal hydroxide or Asp mechanisms, although mutations, in particular D122N (which eliminates the aspartic acid necessary for the Asp mechanism, as well as the corresponding electrostatic repulsion between this side chain and the hydroxide nucleophile) could render this a viable pathway. However, despite the high energies of the bridging hydroxide and concerted pathways, both the terminal hydroxide and Asp mechanisms are energetically plausible, suggesting the presence of a catalytic backup, as observed in PON1^4^ and archaeal protein tyrosine phosphatases.^54–56^ Representative structures of key stationary points for each pathway are shown in **Figures 3**, and **S13** and **S15**.

Overall, our calculations of the hydrolysis of the C6-HSL rule out the bridging hydroxide mechanism (**Figure 1A**) as an energetically viable mechanism, and similarly suggest a high barrier for the concerted mechanism (**Figure 1D**). In contrast, the terminal hydroxide (**Figure 1B**) and Asp (**Figure 1C**) mechanisms are shown to be similar in energy, and competing pathways for the hydrolysis of this lactone.

### Empirical Valence Bond Simulations of the Hydrolysis of a Range of N-Acyl Homoserine Lactones by GcL wild-type and variants

To further explore the viability of backup mechanisms across multiple substrates and enzyme variants, we performed additional EVB simulations of the hydrolysis of the C4-, C6-, and C10-HSLs (**Figure S1**) by wild-type GcL, as well as by the Asp122Asn, Gly156Pro, Ala157Gly, Ala157Ser, Tyr223Phe and Ile237Met GcL variants, following experimental data presented in **Table 1**. As the bridging hydroxide mechanism appears not to be energetically viable (**Table 2**), we focus here on modeling lactone hydrolysis proceeding through the terminal hydroxide, Asp, and concerted mechanisms (**Figures 1B** through **D**). The resulting data is shown in **Tables S5** – **S8**, and a comparison of experimental and calculated activation free energies is shown in **Figure 4**.

**Figure 4.**
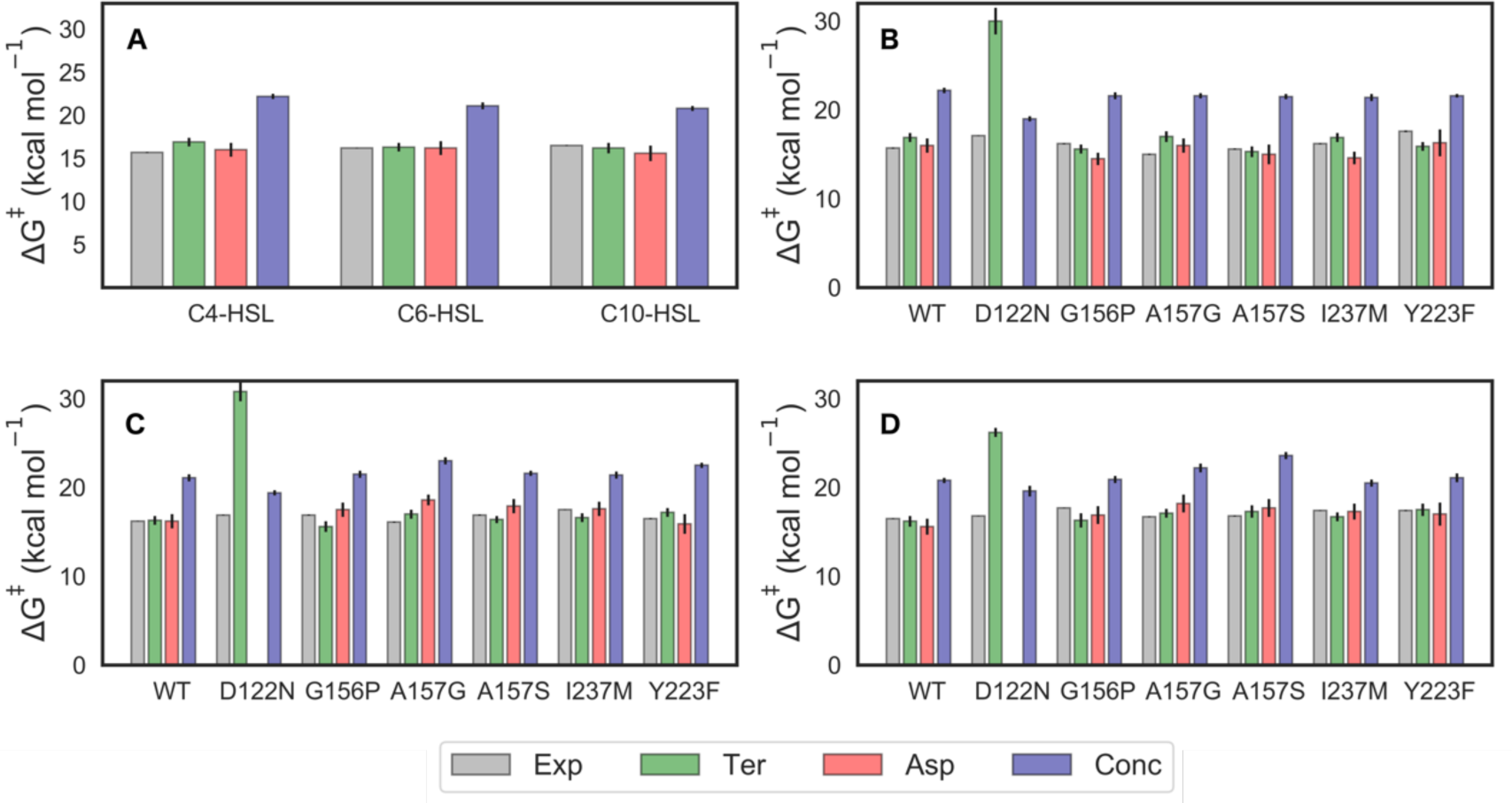
Comparison of the experimental (ΔG^‡^_exp_, gray) and calculated activation free-energies for the terminal hydroxide (Ter), Asp and concerted (Conc) mechanisms (ΔG^‡^_calc,_ green, salmon, and blue, respectively, see **Figure 1B** through **D**), for the hydrolysis of (**A**) a range of AHLs by wild-type GcL and (**B, C, D**) C4-, C6- and C10-HSL, respectively, by wild-type GcL and the Asp122Asn, Gly156Pro, Ala157Gly, Ala157Ser, Tyr223Phe and Ile237Met GcL variants. Error bars on the calculated values represent standard error of the mean calculated over 30 discrete EVB trajectories for each system. The corresponding calculated data is shown in **Tables 2,** and **S5** – **S8**. ΔG^‡^_exp_ values and their associated error bars were obtained from the kinetic data shown in **Table 1**.

From this data, it can be seen that as in the case of the C6-HSL (**Table 2**), for all additional substrates and variants studied in this work with the exception of the D122N variant, both the terminal hydroxide and Asp mechanisms appear to be energetically accessible, with calculated activation free energies within ∼2 kcal mol^−1^ of both the experimental data and of each other (**Figure 4** and **Table S8**, note that for the terminal hydroxide mechanism the first nucleophilic attack step is rate-limiting while for the Asp mechanism, the breakdown of the tetrahedral intermediate is rate-limiting, as shown in **Tables S5** and **S6**).

In the case of the D122N variant, the Asp mechanism is no longer accessible due to mutation of D122, leaving either the terminal hydroxide or concerted mechanisms as potential options. Curiously, in this variant, the activation free energy for the terminal hydroxide mechanism increases substantially, leaving the concerted mechanism (**Figure 1D**) as the only energetically plausible pathway with calculated activation free energies within 3 kcal mol^−1^ of the experimental data. Note that while this is still higher than the experimental values, the energy difference between calculated and experimental values is also smaller than for other substrates/GcL variants, where the concerted pathway can be substantively higher in energy than experiment (**Figure 4** and **Table S8**). Visual examination of our EVB trajectories indicates that during our simulations, the substituted N122 side chain interacts with (and thus stabilizes) the terminal hydroxide ion at the Michaelis complex, contributing to higher calculated activation free energies through reactant state stabilization.

In summary, an EVB comparison of hydrolysis of different HSL substrates by wild-type GcL and variants indicates that the energetically preferred mechanism shifts depending on both substrate (tail length) and variant, suggesting that multiple mechanisms are plausible within the same enzyme active site, and that the selected mechanism will depend on precise environmental conditions, similar to prior work on PON1.^4, 5^

### Molecular Dynamics Simulations of Effect of Tail Length on N-Acyl Homoserine Lactone Binding to GcL

In contrast to other lactonases,^30, 31^ GcL is a generalist enzyme and is highly proficient towards AHLs with both short and long acyl chains as well as γ-, δ-, ε- and whiskey lactones, with catalytic efficiencies (*k*_cat_/*K*_M_) in the range of 10^4^ to 10^7^ M^−1^ s^−1^.^36^ However, even in this generalist enzyme, both lactone tail length and substituents impact both *k*_cat_ and *k*_cat_/*K*_M_, with longer lactone tail lengths showing improvements in catalytic efficiency, but diminished turnover numbers (note that the associated energy differences are small, on the range of 1.5 kcal mol^−1^ or less, based on kinetic data shown in **Table 1**). Furthermore, the AHL with the shortest acyl chain, C4-HSL, displays *K*_M_ values that are substantially higher than their longer chain counterparts C6-, C8-, or C10-HSL, and this feature is conserved not only for the wild-type enzyme but also for most of the variants in studied this work (**Table 1**). Our EVB calculations (**Figure 4** and **Table S8**) give good agreement with experiment for individual substrates but are unable to reproduce these rankings as the experimental differences in activation free energy are extremely small (1 kcal/mol or less) and invisible to the resolution of current computational approaches. Therefore, to better understand the drivers of selectivity between different HSL substrates, we performed molecular dynamics simulations to explore the differences in structural stability of these substrates in the active site pocket as well as the binding modes of the lactone tail.

Structurally, the GcL active site comprises of three sub-sites:^36^ a hydrophobic sub-site (comprised of the side chains of Met20, Met22, Phe48 and Tyr223) involved in the accommodation of the lactone ring, a second hydrophobic patch (comprised of the side chains of Trp26, Met86, Phe87, Leu121 and Ile237) that accommodates the amide group and the beginning of the *N-*acyl chain of the substrate, and a hydrophilic region (comprised of the side chains of Ser82, Thr83, Glu155, Gly156 and Ala157), that is open to the protein surface and exposed to bulk water (**Figure 5**).

**Figure 5.**
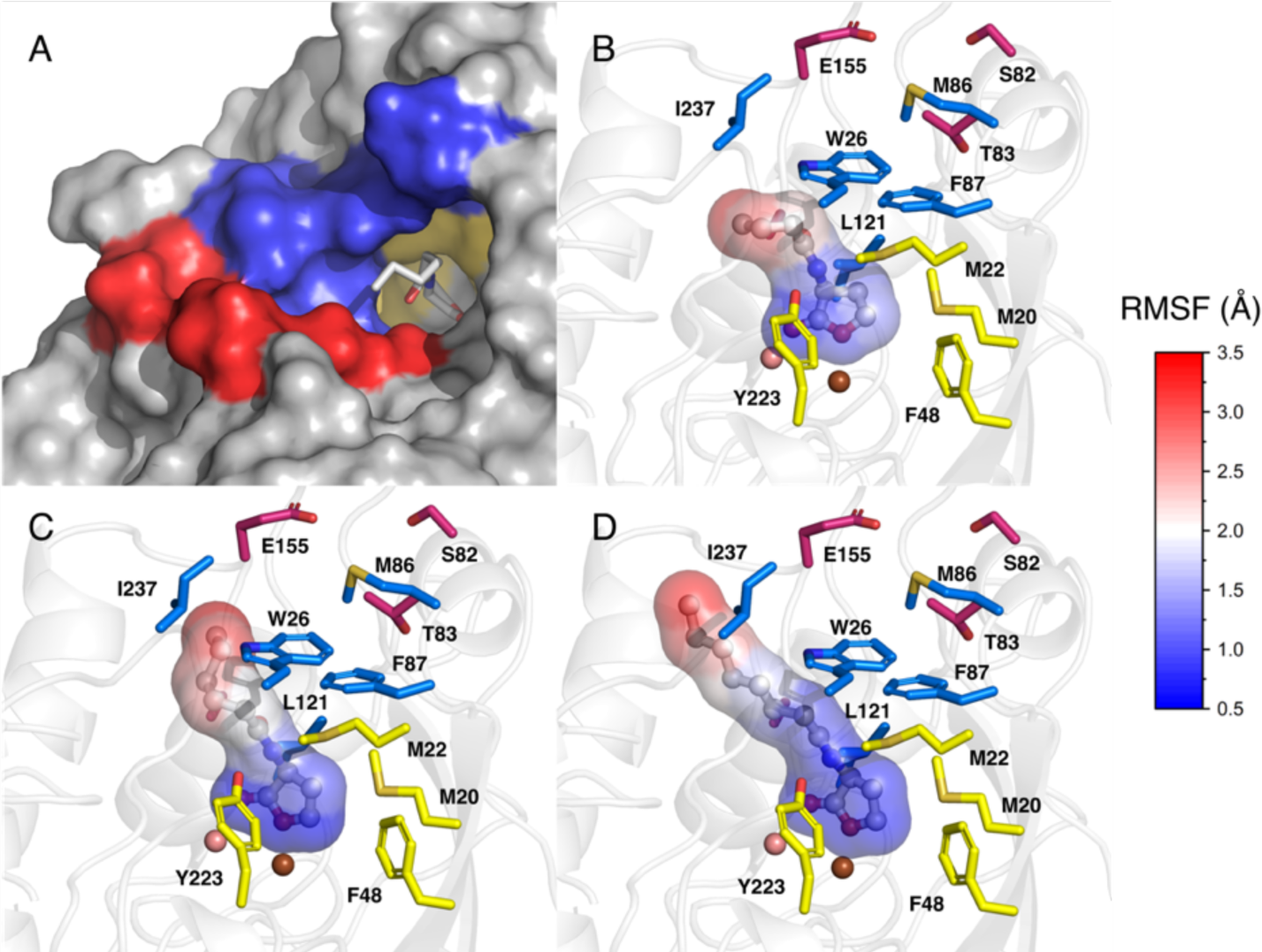
Root-mean squared fluctuations (Å) of the heavy atoms of the C4-, C6- and C8-HSL substrates during molecular dynamics simulations of wild-type GcL. Shown here are (**A**) a view of the active site pocket entrance of GcL in complex with the C6-HSL, with the first and second hydrophobic patches shown in yellow and blue respectively, and the hydrophilic region is shown in red. (**B, C, D**) Close-ups of the positions of the (**B**) C4-, (**C**) C6- and (**D**) C8-HSL substrates in the GcL active site, colored by the root mean square fluctuations (RMSF) of the heavy atoms, to indicate substrate flexibility in the pocket. The side chains of the residues comprising the first and second hydrophobic patches are colored in yellow and blue, respectively, and those comprising the hydrophilic patch are shown in mauve.

To shed light on how different tail lengths may affect the way different AHL substrates interact with the GcL active site, we performed molecular dynamics simulations of wild-type GcL in complex with C4-, C6- and C8-HSL, as described in the **Materials and Methods**. The root mean square fluctuations (RMSF) of the heavy atoms of each substrate during these simulations show that the *N-*acyl chain of the substrate is highly mobile, positioning itself onto different pockets on the protein surface, as illustrated in **Figures 5** and **S16**. In contrast, the metal-coordinating lactone ring is relatively rigid overall, although the shorter the alkyl tail is, the greater (subtly) the flexibility of the ring (**Figure 5**). This flexibility of the tail may in turn render substrate stabilization through interactions with the second hydrophobic sub-site.^36^ This is offset to some extent however by the fact that, based on our simulations, the shorter-chain C4-AHL substrate can bend its acyl tail to fit inside the sub-site binding the lactone ring itself, resulting in the slightly higher *K*_M_ value observed for this substrate (**Table 1**).

Following this, when considering the locations of the amino acid substitutions performed in the variants studied here, two of them (Tyr223Phe and Asp122Asn) are part of the first hydrophobic sub-site where the lactone ring is accommodated, Ile237Met is located at the second hydrophobic sub-site, and the rest (Gly156Pro, Ala157Gly and Ala157Ser) are located in the third hydrophilic sub-site. Especially noteworthy in **Table 1** is the huge increase in the *K*_M_ values towards all the studied AHLs when the polar -OH group of Tyr223 is removed. The hydroxyl group of Tyr223 is hydrogen bonded to the carbonyl oxygen atom of the lactone ring in the crystal structure (PDB ID: 6N9Q^36^ and 9AYT). Simulations of wild-type GcL in complex with C4-, C6- and C8-HSL (**Table S9**) indicate the presence of an interaction between the OH group of the Tyr223 side chain and either the amine nitrogen or the carbonyl oxygen of the alkyl tail for at least 22% of the simulation time (this is most pronounced in simulations with C6-HSL). Our simulations indicate that this is either a direct interaction between the tyrosine side chain and the lactone tail, or a water-mediated interaction with a bridging water molecule (present for an additional ∼10% of simulation time). As this interaction contributes to the stability of the lactone in the active site pocket, clearly its elimination in the Tyr223Phe variant results in the corresponding *K*_M_ values of its substrates as well as its catalytic activity.

Furthermore, while most of the amino acid substitutions summarized in **Table 1** do not lead to important structural changes (based on structural data), the crystal structure of the Gly156Pro variant (PDB ID: 9B2I) reveals structural rearrangement of the Asn152-Ala157 loop, such that the polar residue Glu115 is relocated from pointing out of the binding pocket, to pointing into the binding cleft (**Figure S16**). When examining the effect of this mutation on the reaction kinetics of the different substrates, it is surprising to observe a large increase in the *K*_M_ value of the C6-HSL substrate, while that of C8-HSL remains similar to the wild-type variant. We used MDpocket^68^ to locate the hydrophobic/hydrophilic regions of the cavity along the MD simulations of C6-, and C8-HSL in complex with wild-type GcL and Gly156Pro variant. **Figure 6** shows representative structures from the main clusters (describing the most sampled populations along the simulations) of each substrate in the main representative binding pocket of the relevant GcL variant (obtained from RMSD clustering across our simulations, as described in the **Materials and Methods**), with the corresponding hydrophilic regions colored in red. Interestingly, the end of the C6-HSL acyl chain, which is the most mobile part of the substrate, lays in the same position as the rearranged residue Glu155 side chain, creating strong repulsion and destabilizing the substrate. In contrast, the end of the C8-HSL acyl chain lies further from the active site and the repulsion between Glu155 and the substrate tail is diminished, allowing tighter binding of C8-compared to C6-HSL.

**Figure 6.**
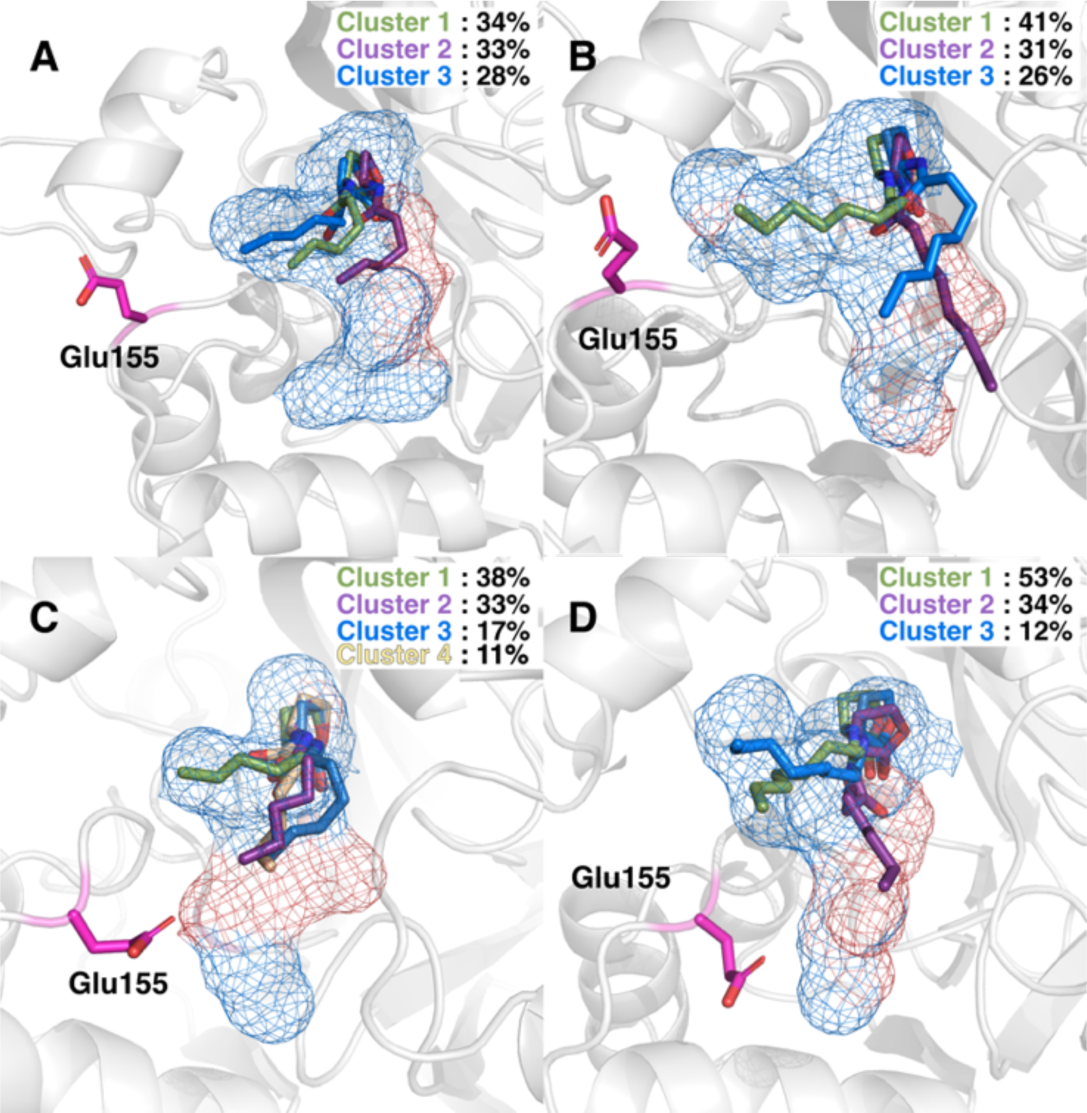
Main conformation of the binding pocket from RMSD clustering of our MD simulations of (**A**, **C**) C6- and (**B**, **D**) C8-HSL, in complex with (**A**, **B**) wild-type GcL and **(C**, **D)** the Gly156Pro variant, The binding pocket is shown as a blue grid, with the hydrophilic regions colored in red. Substrate structures from the three/four principal clusters of each system (obtained from RMSD clustering) in complex with C6-, or C8-HSL, as well as the dominant position of the Glu155 side chain are highlighted. Only substrates structures from clusters accounting for more than 10% of the simulation time are shown here.

Finally, **Figure 6** also illustrates significant conformational flexibility of the alkyl tail of both C6- and C8-substrates in both wild-type GcL and the Gly156Pro variant, illustrating the conformational heterogeneity of the substrate in the active sites and the fact that it can occupy multiple binding poses of the alkyl tail. This effect is significantly more pronounced in the complex of Gly156Pro with C8-HSL than that with C6-HSL, suggesting that part of the reason for the better *K*_M_ value of this variant towards C8-HSL is simply due to its ability to be accommodated through multiple binding modes in the active site pocket, avoiding Glu155. This flexibility in binding mode will also affect how different AHL substrates interact with different GcL variants, as amino acid substitutions reshape the active site pocket. When this is coupled with the ability of short-chain HSLs such as the C4-HSL to explore and occupy new binding pockets (**Figure 5**), this conformational plasticity will impact both activity and substrate specificity, as shown in **Table 1**.

### Empirical Valence Bond Simulations of Lactone Hydrolysis by other metallo-β-lactamase-like lactonases

Our EVB simulations support the existence of catalytic redundancies in GcL and provides a molecular rationale for this redundancy, as well as the associated substrate specificity towards different AHL substrates. Such catalytic backups have been previously suggested in the case of an analogous lactone, PON1.^4^ Furthermore, several archaeal protein tyrosine phosphatases have been suggested to operate *via* dual general acid mechanisms, with built-in redundancies in the active site.^54–56^ As GcL and PON1 (let alone protein tyrosine phosphatases) have rather different active site architectures, in particular in terms of the identity and coordination of the metal centers involved, this then raises the question of whether catalytic redundancies and backups are a common feature of promiscuous lactonases (and enzymes more broadly).

To address this question in the context of lactonases, we extended our EVB simulations to two additional MLLs: AiiA and AaL. These systems were selected because of the structural similarity of the corresponding binding domains, the availability of high resolution crystal structures,^34, 58^ and the availability of kinetic data for these enzymes against C6-HSL,^27, 69^ allowing for direct comparison to our GcL simulations (**Table 2**). The overall structure of AaL is very similar to GcL with an RMSD of 0.42 Å (sequence identity 81.1%). There are larger differences between AiiA and GcL, with an RMSD of 1.22 Å between the two structures (both proteins share sequence identity 24.4%), as well as the absence of a protruding loop in AiiA involved in dimerization in GcL and AaL,^34, 36^ where AiiA is instead organized as a monomer.^70^ Noteworthy is that Tyr223 is conserved in all MLLs, with the exception of AidC.^32^ Furthermore, the different regions of the AHLs binding pocket are highly conserved within the three lactonases (**Figure S17)**.

We extended our EVB simulations of all studied mechanisms to the AiiA and AaL lactonases in complex with C6-HSL using the same set of parameters as in GcL (**Table 3**). As in wild-type GcL (**Tables 2** and **S8**), the bridging hydroxide and concerted mechanisms **(Figures 1A** and **D**) yield too high activation free energies and are therefore unlikely, while both the bridging hydroxide and Asp mechanisms (**Figures 1B** and **C**) are energetically feasible and within range of the experimental data. We note the slightly lower calculated activation free energies for the bridging hydroxide mechanism for all enzymes studied; however, this could be the same underestimation of the activation free energy for this mechanism for the hydrolysis of the C6-HSL substrate, as in the case of GcL. Based on this data, we demonstrate that the mechanistic plasticity of GcL is conserved across these three diverse lactonases and is not a feature unique to GcL.

**Table 3.** A comparison of experimental and calculated activation free energies (kcal mol^−1^) for the hydrolysis of C6-HSL catalyzed by three wild-type MLLs, GcL, AiiA and AaL, through the different mechanisms considered in this work.^a^

|  | **GcL** | **AiiA** | **AaL** |
| --- | --- | --- | --- |
| **Experiment** | 16.2 | 14.8 | 15.9 |
| **Bridging Hydroxide Mechanism** | 36.7 ± 1.1 | 25.6 ± 0.4 | 29.7 ± 0.4 |
| **Terminal Hydroxide Mechanism** | 16.3 ± 0.5 | 15.5 ± 0.7 | 14.9 ± 0.8 |
| **Asp Mechanism** | 16.2 ± 0.8 | 13.8 ± 0.3 | 14.4 ± 0.7 |
| **Concerted Mechanism** | 21.1 ± 0.4 | 18.4 ± 0.3 | 25.4 ± 0.9 |

<sup>a</sup> The different mechanisms considered here are illustrated in **Figure 1**. All values shown here are averages and standard error of the mean over 30 independent EVB trajectories, shown in kcal mol<sup>-1</sup>. Experimental (exp) activation free energies for the hydrolysis of C6-HSL by each of GcL, AiiA and AaL are obtained based on kinetic data provided in refs. <sup>27, 69</sup> (**Table S10**).

## Overview and Conclusions

Lactones have a wide range of biological activities,^71^ including as antimicrobial agents,^72^ anti-inflammatory compounds,^73^ antitumor agents^74, 75^ and mycotoxins^76^, as well as being abundant in cellular metabolism.^77^ The importance of lactones to bacterial communication (quorum sensing)^78^ and biofilm formation^13^ makes the enzymes that degrade them biotechnologically important as quorum-quenching agents for a whole host of industrial and biomedical applications. Because of this diversity of roles of lactones in biology, the actual primary purpose of many lactonases remains unclear: for example the first identified lactonase AiiA,^70, 79^ exhibits millimolar *K*_M_ values towards typical AHL substrates and is a broad generalist, with very little discrimination between substrates with varying chain lengths and/or substitutions on the chain.^27, 30, 58, 80^ It is therefore unclear whether AiiA evolved specifically for the purpose of quenching microbial signaling: it is clearly capable of doing so, but likely has much broader biological purpose than just quorum quenching. Serum paraoxonase 1 (PON1) is a lactonase/organophosphate hydrolase^20^ that acts as a broad scavenger enzyme. In contrast, GcL, the primary focus of this study, remains a generalist enzyme but with micromolar *K*_M_ values, making it perhaps more likely specialized towards quorum quenching as its native function.

We examined the mechanism and substrate selectivity of wild-type GcL as well as two related lactonases, AiiA and AaL (sequence similarity to GcL 41.5 and 89.2%, respectively), through a combination of structural, biochemical, and computational approaches. Our structural and mutagenesis analysis point to the likely important roles for both Asp122 and Tyr223 in catalysis and substrate positioning, yet no residue seems completely required for catalysis. Remarkably, our mechanistic analyses indicate that there are not one but three viable (and energetically similar) mechanisms for AHL hydrolysis by GcL with the preferred mechanism between the three mechanisms shifting, depending on both different AHL substrates and different enzyme variants. This is complemented by substrate plasticity in the active site, with the alkyl tail of the AHL substrates taking on multiple conformations depending on tail length and enzyme variant. This mechanistic redundancy is observed again in our simulations of both AiiA, and AaL as well as in both computational and experimental studies of other enzymes such as PON1^4, 5^ and archaeal protein tyrosine phosphatases.^54–56^ The importance of conformational dynamics to enzyme selectivity and evolvability is by now well established.^81–88^ Our data indicate that, similarly to catalytic promiscuity and broad substrate specificity, mechanistic promiscuity also plays an important role in modulating enzyme activity and selectivity. This expands the question from “how does a promiscuous enzyme chooses a specific substrate from a pool of different substrates?” to also “how does the enzyme utilize a specific mechanism from a pool of different mechanisms?” and “what makes an active site catalytically versatile?” This is a broader issue requiring examination as it is observed across an increasing number of systems, and is a significant consideration when engineering generalist enzymes for specific functions.

## Materials and Methods

### Mutagenesis

Site-directed mutagenesis was performed for the mutations D122N and Y223F using Pfu polymerase (Invitrogen) on 100 ng of plasmid using primers (**Table S11**), with an annealing temperature of 63°C for 34 cycles. After DpnI digestion, plasmids were concentrated by alcoholic precipitation and then transformed (Gene-Pulser, Bio-Rad) into *Escherichia coli* cells DH5α (Invitrogen) by 30 seconds of heat shock at 42 °C.

Additional GcL mutations of key positions identified from the previous structural analysis of GcL and inferred from structural analysis^36^ were ordered from Genscript Biotech Corporation (catalog SC2029) as part of saturation mutagenesis libraries designed to optimize GcL properties (results from this engineering efforts will be reported in details in a separate work). 100 ng of the pooled plasmid library was introduced into *E. coli* DH5α by heat shock transformation at 42 °C, and the cells were spread onto LB agar supplemented with ampicillin. Individual colonies were resuspended in phosphate-buffered saline and sent to ACGT Inc. for direct colony sequencing. Identified single mutant plasmids were expressed in *E. coli* DH5α and purified using a Qiagen miniprep kit. Purified mutant plasmids were confirmed by sequencing (University of Minnesota Genomics Center) before finally being transformed into an *E. coli* protein production strain by heat shock.

### Protein Production

The various proteins were produced in *Escherichia coli* strain BL21(DE3)-pGro7/GroEL strain (TaKaRa). The sequences were enhanced by an N-terminal strep tag (WSHPQFEK) with a TEV cleavage site sequence (ENLYFQS). Protein productions were performed at 37 °C in the autoinducer media ZYP (with 100 mg.ml^−1^ ampicillin and 34 mg.ml^−1^ chloramphenicol). When cells reached the exponential growth phase chaperone GroEL was induced by adding 0.2% L-arabinose, 2 mM CoCl_2_ was added, and the cultures were transitioned to 18 °C for 16 hours. Cells were harvested by centrifugation (4400 xg, 4 min, 4 °C), resuspended in lysis buffer (150 mM NaCl, 50 mM HEPES pH 8.0, 0.2 mM CoCl_2_, 0.1 mM PMSF and 25 mg mL^−1^ lysozyme), and left on ice for 30 minutes. Cells were then sonicated (amplitude 45% in three steps of 30 seconds; 1 pulse-on; 2 pulse-off) (Q700 Sonicator, Qsonica, USA). Cell debris was removed by centrifugation (5000 xg, 45 min, 4°C).

### Protein Purification

The lysis supernatant was loaded on a Strep Trap HP chromatography column (GE Healthcare) in *PTE buffer* consisting of 50 mM HEPES pH 8.0, 150 mM NaCl, and 0.2 mM CoCl_2_ at room temperature. TEV cleavage was performed by adding the Tobacco Etch Virus protease (TEV, reaction 1/20, w/w) overnight at 4 °C. Then, the sample was loaded on a size exclusion column (Superdex 75 16/60, GE Healthcare) to obtain pure protein. SDS-PAGE was performed to confirm the identity and purity of the proteins. Proteins were quantified by measuring their absorbance at 280 nm and using the Beer-Lambert law. The protein molecular extinction coefficient was generated using the protein primary sequence and the ProtParam tool implemented into ExPASy.^89^

### Determination of Kinetic Parameters

All kinetic experiments were performed in at least triplicates in 200 µL reaction volumes using 96-well plates (6.2mm path length cell) and a microplate reader (Synergy HT), using the Gen5.1 software at 25 °C. The time course of the hydrolysis of AHL substrates was analyzed by monitoring the decrease in absorbance at 577 nm. Lactone hydrolysis assays were performed in lactonase buffer (2.5 mM bicine pH 8.3, 150 mM NaCl, 0.2 mM CoCl_2_, 0.25 mM m-cresol purple, and 0.5% DMSO), using m-cresol purple (p*K*_a_ 8.3 at 25 °C) as a pH indicator to follow the acidification caused by lactone ring hydrolysis. The molar extinction coefficient was measured by recording the absorbance of the buffer over a range of acetic acid concentrations between 0-0.35 mM. The initial rates of reactions were fitted using Graph-Pad Prism 5 for Windows (GraphPad Software, San Diego, California, USA) and fitted to a Michaelis-Menten curve to obtain the catalytic parameters. Due to the limits in the sensitivity of this pH-indicator based assay at low substrate concentration, measurements for substrate with very low *K*_M_ values are challenging and can result in poorer fit to the Michaelis-Menten equation. Replicates with technical errors (e.g. pipetting errors or failed) were excluded from the Michaelis-Menten analysis.

The catalytic activity of the enzymes against ethyl-paraoxon was monitored using a previously described assay.^36^ The reaction was monitored by following the production of paranitrophenolate anions at 405 nm. Reactions were performed in 50 mM HEPES pH 8.0, 150 mM NaCl, and 200 mM CoCl_2_ using *ε*_405nm_ =17,000 M^−1^.cm^−1^.

### Crystallization, Data Collection, and Refinement

Crystallization of GcL wild-type and variants was performed using protein samples concentrated to 10.0 −11.5 mg mL^−1^ using the hanging-drop vapor-diffusion method and previously reported conditions.^36^ The best crystals were produced with 1.0 −1.25 M ammonium sulfate and 0.1 M sodium acetate, pH 4.0 −5.5. The structure in complex with the substrate C6-HSL was obtained by soaking the crystals for 10 minutes in a solution containing the cryoprotectant and 20 mM of C6-HSL. The GcL complex with the reaction product of C8-HSL was obtained by co-crystallizing the enzyme with 20 mM (final). The crystals were cryoprotected in a solution composed of 30% PEG 400 and frozen in liquid nitrogen. X-ray diffraction datasets were collected at 100 K using synchrotron radiation on the 23-ID-B and 23-ID-D beamlines at the Advanced Photon Source (APS, Argonne, Illinois, USA, **Table S1**). The structures were resolved in the H3 space group for the Asp122Asn-2metals and the structure bound to C6-HSL, and in C2 for the other structures. The integration and the scaling of the X-ray diffraction data were performed using the XDS package.^90^ The molecular replacement was performed using the wild-type GcL structure as a model (PDB ID: 6N9I^36^) and using MOLREP.^91^ Manual model construction was performed with Coot.^92^ Cycles of refinement were performed using REFMAC.^93^ Statistics are shown in **Table S1**. We note that the ligand occupancy in the structures is variable in the different monomers that are present in the asymmetric units, and the highest occupancy models (presented) are 0.8 for the C6-HSL and 0.7 for the C8-HSL product. These occupancy levels limit the accuracy of the models.

### Empirical Valence Bond Simulations

The empirical valence bond (EVB) approach^53^ is a force-field based approach that describes chemical reactivity within a valence-bond based quantum mechanical framework. This approach has been used extensively to describe enzyme reactivity in general,^94, 95^ and lactone hydrolysis in particular.^5, 59, 60^ In this work, we have modelled the hydrolysis of the C4-, C6- and C10-HSL (**Figure S18**) by wild-type AiiA, AaL and GcL, as well as the Asp122Asn, Gly156Pro, Ala157Gly, Ala157Ser, Tyr223Phe, and Ile237Met GcL variants, through four different mechanisms shown in **Figure 1**. All four mechanisms were tested for the hydrolysis of C6-HSL by wild-type GcL, and the energetically accessible pathways, *i.e.* the terminal hydroxide, Asp and concerted mechanisms, were tested for the hydrolysis of all other compounds. The corresponding valence bond states are shown in **Figure S19**. Note that the bridging and terminal hydroxide mechanisms shown in **Figure 1** use identical valence bond states for the first step of the reaction, as the only difference between them is whether the nucleophile is in the metal coordination of the hydroxide ion. In addition, the first three mechanisms considered in **Figure 1** are all 3-state stepwise processes, whereas the final mechanism is a 2-state concerted process. In the case of the stepwise processes, the intermediate state structures generated at the endpoint of EVB simulations of the first step for each replica were used as starting points for EVB simulations of the second step of the reaction. All relevant input and parameter files necessary to reproduce our calculations, as well as snapshots from our simulation trajectories, have been uploaded as a data package to Zenodo, DOI: 10.5281/zenodo.11072674. Full details of system setup and EVB simulations, are provided in the **Supporting Information** and summarized here in brief. All EVB simulations were performed using the *Q5* simulation package^96^ and the OPLS-AA force field,^97^ as implemented into *Q5*. Metal and ligand parameterization, and calibration of the EVB off-diagonal term and gas-phase shift, were performed as described in the **Supporting Information**. The same parameter set was then transferred unchanged in simulations of each substrate with all enzyme variants, as the EVB off-diagonal term has been shown to be phase-independent and thus transferable.^98, 99^

Simulations of wild-type AaL were performed using the structure of wild-type AaL in complex with C6-HSL, obtained from the Protein Data Bank^100^ (PDB ID: 6CGZ^34^), while for wild-type AiiA simulations, the structure of AiiA in complex with C6-HSL hydrolytic product was used (PDB ID: 3DHB^58^), where the product was replaced with C6-HSL by aligning AiiA and AaL wild-type structures. Due to the high negative charge of the system that fits inside the water droplet in the case of AiiA, 10 Na^+^ counterions were added to neutralize the system. These counterions were placed so that they interact with negatively charged residues near the surface of the enzyme, that still fall within the water droplet. All simulations of wild-type GcL were performed using the structure of wild-type GcL in complex with C4-HSL and C6-HSL (PDB IDs: 6N9Q^36^ and 9AYT). The structure of the Tyr223Phe variant was generated by manual deletion of the Tyr223-OH group from the wild-type structure, while the Asp122Asn, Gly156Pro, Ala157Gly, Ala157Ser and Ile237Met substitutions were introduced by use of the Dunbrack 2010 Rotamer Library,^101^ as implemented in USCF Chimera, v. 1.14.,^102^ trying to reproduce as much as possible the rotamer found in the crystal structures where unliganded structures including those substitutions were available (PDB IDs: 9B2L, 9B2I and 9B2J for the Asp122Asn, Gly156Pro, and Ile237Met variants, respectively). The reason the unliganded structures were not used directly for these simulations is that the loop comprising residues 236-238 is found in a closed conformation in the unliganded structures, creating a steric clash with the substrate tail when aligned with the liganded complex, whereas it opens up in the liganded complex (based on its conformation in the wild-type structure).

All the variants were simulated in complex with the different substrates of interest to this work (**Figure S18**). In each case, starting structures for all Michaelis complexes with the substrates C4-, C6- and C10-HSL were either taken directly from the crystal structure (where a liganded complex was available), or generated manually based on an overlay with the coordinates for C4- or C6-HSL in the wild-type structure (PDB IDs: 6N9Q^36^ and 9AYT, respectively).^36, 100^ The system was then solvated in a 30 Å droplet of TIP3P water molecules,^103^ described using the surface constrained all-atom solvent (SCAAS) approach.^104^ The protonation states of all relevant ionizable residues, as well as histidine protonation patterns, are provided in **Table S12**. The starting structures for the terminal hydroxide mechanism were generated as described above but with rotation of the lactone ring to place an extra hydroxide ion on the Fe^2+^ metal center (The starting structures used in our simulations can be found in DOI: 10.5281/zenodo.11072674).

All systems were gradually heated from 1 to 300 K, as described in the **Supporting Information**, followed by 50 ns of molecular dynamics equilibration at the target temperature, the convergence of which is shown in **Figure S20-S23**. 30 individual replicas were generated per system using different random seeds to assign initial velocities. The endpoint of each equilibration was used as the starting point for 30 subsequent EVB simulations (1 EVB simulation per replica), which were performed using the valence bond states shown in **Figure S19**, using 51 EVB mapping windows of 200 ps/length each (*i.e.* 10.2 ns simulation time per EVB trajectory). This led to a cumulative total of 1.5 μs equilibration and 306 ns EVB sampling per system and mechanism (612 ns EVB sampling for the mechanisms comprising a 3-state process), and a total of 154.6 μs simulation time (equilibration + EVB) over all systems.

All equilibration and EVB simulations were performed using the leapfrog integrator with a 1 fs time step, using the Berendsen thermostat^105^ to keep the temperature constant with a 100 fs bath coupling time, and with the solute and solvent coupled to individual heat baths. Long-range interactions were treated using the local reaction field (LRF)^106^ approach, while cut-offs of 10 and 99 Å were used for the calculation of non-bonded interactions involving the protein and water molecules and the EVB region respectively (effectively no cut-off for the latter). In all but the very initial minimization step to remove bad hydrogen contacts, the SHAKE^107^ algorithm was applied to constrain all bonds involving hydrogen atoms. Further simulation details, as well as details of simulation analysis are provided in the **Supporting Information**.

### Molecular Dynamics Simulations

All molecular dynamics simulations were performed using the GROMACS v2021.3 simulation package,^108, 109^ in combination with the OPLS-AA force field^97^ for compatibility with our EVB simulations. In this work, we have performed simulations of the wild-type GcL, as well as G156P and Y223F variants, in complex with C4-, C6- and C8-HSL. The starting structure for our simulations was taken from crystallographic coordinates of wild-type GcL in complex with C4- and C6-HSL (PDB IDs: 6N9Q^36^ and 9AYT). The histidine protonation patterns in our MD simulations were identical to those used in our EVB simulations, as listed in **Table S12**, and all other ionized residues (Asp, Glu, Arg, and Lys) were modeled in their standard ionization states at physiological conditions, *i.e*. Asp and Glu side chains were negatively charged while Arg and Lys side chains were positively charged. The resulting complex was put in the center of an octahedral box filled with TIP3P water molecules,^103^ with at least 10 Å distance between the surface of the complex and the edge of the box. Na^+^ ions were added to neutralize each system. After the system setup was complete, three independent replicas were generated, where a 5000-step minimization was performed on each system using the steepest descent and conjugate gradient methods, followed by heating of the solvated system from 0 to 300 K over a 500 ps MD simulation in an NVT ensemble, using the velocity rescaling thermostat^105, 110^ with a time constant of 0.1 ps for the bath coupling. This was again followed by a further 500 ps of simulation in an NPT ensemble at 300 K and 1 bar, controlled by the same thermostat and a Parrinello-Rahman barostat^105^ with a time constant of 2.0 ps. Positional restraints of 2.4 kcal mol^−1^ Å^−2^ were applied on every heavy atom in each of the xyz directions for the first two steps of equilibration. Afterwards, the positional restraints were released and instead, distance restraints were applied between all the side chains coordinating the dummy particles and the metal centers, including the ligand, and the central atom of the dummy complex, during the first 25 ns of production to ensure crystallographic ligand coordination around the metal ions is maintained. These distance restraints were set to 40 kcal mol^−1^ Å^−2^ during the first 20 ns of simulation time, with the force constant halved to 20 kcal mol^−1^ Å^−2^ the last 5 ns of simulation time. Finally, 500 ns of unrestrained molecular dynamics simulations (x 3 replicas) were performed for each system, the convergence of which are shown in **Figure S24**. For all the simulations, 12 Å non-bonded interaction cut-off was used to evaluate long range electrostatic interactions, using the Particle Mesh Ewald (PME) algorithm,^111^ and the LINCS algorithm^112^ was applied to constrain all hydrogen bonds, using a 1 fs time step.

### Simulation Analysis

The physicochemical properties of the active site pocket of the wild-type, Gly156Pro and Tyr223Phe GcL enzymes when either C4-, C6- or C8-HSL is bound, were tracked along the corresponding conventional molecular dynamics simulations trajectories using the MDpocket^68^ tool, published within the fpocket^113^ suite of pocket detection programs. To account for the structural differences inferred by each ligand on the cavity, the liganded trajectories were used where the corresponding ligand was stripped from the cavity to assess the pocket. All cavities were identified using a frequency iso-value of 0.7 and points corresponding to the active site pocket were selected and tracked throughout the trajectories.

All other analyses were performed using the CPPTRAJ^114^ module of the AmberTools19^115^ suite of programs. The most-populated structures were obtained by clustering together 3 independent 500 ns MD simulations for each system using a hierarchical algorithm and selecting the centroid of the top-ranked clusters. The clustering was performed based on pairwise RMSD calculations over all the atoms of the ligand. All hydrogen bonds formed between the AHL and Tyr223 were identified using a donor-acceptor distance cut-off of 3.5 Å, and a donor-hydrogen-acceptor angle of 135±45°. Only hydrogen bonds with an occupancy of >1% of the cluster simulation time were considered. Root mean square fluctuations (RMSF, Å) of the heavy atoms of the ligand, were calculated over 3 independent 500 ns MD simulations for each system.

## Acknowledgments

This work was supported by the Knut and Alice Wallenberg Foundation (grant numbers 2018.0140 and 2019.0431), the Swedish Research Council (grant number 2019-03499), the National Institute of General Medical Sciences (award no. R35GM133487 to MHE), and the MnDrive initiative (to MHE). The simulations were enabled by resources provided by the Swedish National Infrastructure for Computing (SNIC) at multiple supercomputing centers (NSC, HPC2N, UPPMAX), partially funded by the Swedish Research Council through grant agreement no. 2016-07213. Further simulations were performed at the Barcelona Supercomputing Center (QSB-2019-2-0005). We are very grateful to the scientists at the Advanced Photon Source (APS, Argonne, IL, USA) and particularly the beamline scientists and coordinators at 23ID-D and 23ID-B for their assistance. The content is solely the responsibility of the authors and does not necessarily represent the official views of the National Institutes of Health.

## Supporting Information

Additional computational methodology, simulation analysis, and kinetic data is provided as Supporting Information.

