## Supplementary Information for "Catalytic Redundancies and Conformational Plasticity Drives Selectivity and Promiscuity in Quorum Quenching Lactonases"

† Current address: Univ. Grenoble Alpes, CNRS, CEA, IBS, F-38000 Grenoble

Correspondence:

**S1. Supplementary Methodology……………………………………………………….….S3**

Parameterization and Calibration of Empirical Valence Bond Simulations…………………..S3

System Preparation for Empirical Valence Bond Simulations………………………………..S4

Empirical Valence Bond Simulation Details…………………………………………………S5

**S2. Supplementary Figures………………………………………………………………S9**

**S3. Supplementary Tables……………………………………………………………...S33**

**S4. Supplementary References………………………………………………………S48**

### S1. Supplementary Methodology

***Parameterization and Calibration of Empirical Valence Bond Simulations***

All simulations were performed with the protein described using the OPLS-AA force field,[1] as implemented into *Q5*.[2] OPLS-AA-compatible structural and van der Waals parameters used to describe lactone hydrolysis were obtained using Schrödinger's Macromodel (version 10.5),[3] whereas partial charges were obtained at the HF/6-31G(d) level of theory, using the standard RESP protocol,[4] and calculated using Gaussian09 Rev. E.01[5] and Antechamber.[6] The active site iron and cobalt ions were modeled using a multisite cationic dummy model, using parameters provided in earlier work.[7] This greatly improves the stability of the metal center in the simulations, while correctly describing the key thermodynamic properties of the metal without the need for any artificial restraints or bonds between the metal and ligands.[7] However, in the case of the Asp122 and Asp220 side-chains, the charges on the oxygen atoms were modified on the basis of charge distributions from quantum mechanical calculations of propanoic acid coordinated to cobalt and iron centers, with the coordination shell completed with extra water molecules. The atomic charges were obtained using Gaussian09 Rev. D.01,[5] using the M11L density functional[8] and the 6-31G* basis set, with implicit solvation described using the polarized continuum model (PCM).[9] The resulting partial charges for each aspartic acid side-chain are shown in **Table S13**.

Central to the philosophy of the EVB approach is the existence of a well-defined reference state, which can be the non-enzymatic reaction in vacuum or solvent, or, for example, the wild-type enzyme for comparison against a set of enzyme variants. The EVB off-diagonal term and gas-phase shift (which are described in detail in *e.g.* refs. [10, 11]) are calibrated to reproduce the activation and reaction free energies for the reference state obtained from either (preferably) experimental or (where experimental data is not available) higher level quantum mechanical calculations, and, as the off-diagonal term is phase independent,[12, 13] the same parameter set is then transferred unchanged to all enzyme variants of interest.

For all mechanisms shown in **Figure 2** of the main text, our reference state was the non-enzymatic reaction in aqueous solution. In the case of the stepwise processes (**Mechanisms A** to **C**), the EVB parameters (**Table S14**) were calibrated to reproduce activation free energies of $\Delta G_1^{‡}$ = 21.8 kcal mol$^{-1}$, $\Delta G_{int}$ = 14.5 kcal mol$^{-1}$, and $\Delta G_2^{‡}$ = 17.5 kcal mol$^{-1}$ for lactone hydrolysis in basic conditions (where $\Delta G_1^{‡}$ and $\Delta G_2^{‡}$ are the free energies of activation for nucleophilic attack and ring opening, respectively, and $\Delta G_{int}$ is the free energy of the corresponding tetrahedral intermediate), based on quantum chemical calculations[14] and experimental rates for the hydrolysis of a range of AHLs.[15] In the case of the concerted pathway, or neutral hydrolysis (**Figure 1D**), the activation barrier was fitted to $\Delta G_1^{‡}$ = 25.0 kcal mol$^{-1}$, based on analogy to experimental data.[15] For both the stepwise and concerted pathways, we set $\Delta G_0$ = 0.0 kcal mol$^{-1}$ for simplicity, as the thermodynamic barrier to the final product state merely shifts the absolute position of the EVB parabola relative to each other, and does not affect the relative energies of any transition and intermediate states.

***System Preparation for Empirical Valence Bond Simulations***

Starting structures for each of our enzyme-substrate complexes were obtained as described in the main text **Materials and Methods section.** Michaelis complexes were manually generated based on overlay of the coordinates of either the C4-, C6- or C10-HSL present in the initial structure, as outlined in the main text. Specifically, in all cases, the lactone ring and the amide moiety were kept in the same position as in the initial structure, but the lipid tail was elongated based on tail position in the PDB structure of wild-type GcL in complex with the 3-oxo C12-HSL hydrolysis product (PDB ID: 6N9R). In the case of the metal ions, the central atom of the multisite model was aligned with the metal coordinates obtained from the crystal structure and the model was placed into the active site such as to optimize interactions between the dummy particles and the ligands coordinating each metal ion. In the case of the enzyme variants for which crystal structures were not available, starting complexes were generated as described in the main text.

Each system was then solvated in a 30 Å radius water droplet of TIP3P[16] water molecules, centered on the oxygen atom of the bridging hydroxide ion, subject to surface constrained all-atom solvent (SCAAS) boundary conditions.[17] In this model, all atoms within the inner 85% of the solvent sphere were allowed to move freely, while atoms in the external 15% and outside of the solvent sphere where subject to 10 and 200 kcal $mol^{-1}$ $Å^{-2}$ harmonic position restraints, respectively, to retain their crystallographic positions. All protein atoms outside this droplet were restrained to their crystallographic positions by the aforementioned 200 kcal $mol^{-1}$ $Å^{-2}$ harmonic position restraints and kept in their neutral forms in order to avoid system instabilities during the simulation due to the presence of charged residues outside the explicitly solvated region. The protonation states of all ionizable side-chains and the protonation patterns of all histidine side-chains within the inner 85% of the solvent sphere were assigned on the basis of standard $pK_a$ values at physiological pH using PROPKA 3.1[18] and verified by visual inspection (**Table S12**).

Non-enzymatic systems were prepared using the same starting structures as for the enzymatic complexes, for each mechanism, but removing the protein and only simulating the lactone and the catalytic water, hydroxide and/or Asp122 side-chain, with the $C_\alpha$-carbon atom saturated with hydrogens. The system was then solvated as in the enzymatic systems.

***Empirical Valence Bond Simulation Details***

Once system setup was completed, all systems were first minimized for 3 ps at 1 K using a 0.1 fs step size in order to ensure the removal of steric clashes and bad contacts within the system after solvation. During this minimization step, a 200 kcal $mol^{-1}$ $Å^{-2}$ harmonic restraint was placed on all heavy atoms of the system, to maintain crystallographic coordinates.

Subsequently, the step size was increased to 1 fs, while the temperature was gradually increased from 1 to 300 K while simultaneously decreasing the harmonic restraints on the mobile region of the system (the inner 85% of the water droplet) from 200 to 5 kcal mol$^{-1}$ Å$^{-2}$ over the course of 210 ps of simulation time. After reaching the target temperature of 300 K, each system was subjected to a further 50 ns of equilibration. In parallel, 100 kcal mol$^{-1}$ Å$^{-2}$ harmonic positional restraints were applied to the heavy atoms of the AHL substrate, the nucleophilic water molecule / terminal hydroxide ion (as relevant), the metal ions, the bridging hydroxide ion, and the side-chains of the two metal coordinating aspartic acid residues (D122 and D220), as well as the Tyr223 side-chain in the case of the concerted mechanism (**Figure 1D**). These restraints were gradually reduced during the first 8 ns of simulation time, until only weak 0.5 kcal mol$^{-1}$ Å$^{-2}$ harmonic positional restraints remained on the AHL substrate, the nucleophilic water molecule / hydroxide ion (depending on mechanism) and, in the case of the Asp mechanism (**Figure 1C**), the side-chain of Asp122 which acts as a general base in the reaction, for the remaining simulation time. The only other restraints applied to the enzyme during our EVB simulations were two 25 kcal mol$^{-1}$ Å$^{-2}$ flat-bottomed harmonic distance restraints to keep the oxygen atom of the Asp220 side-chain bridging the two metal ions (**Figure 1**) between 2.0 and 2.5 Å from the central atom of the multisite models used to describe each metal ion, in order to maintain the bridging position of this oxygen atom and preventing the side-chain from rotating to double coordinate the two metal ions.

All initial equilibrations were performed at the approximate transition state of the reaction ($\lambda$ = 0.5), with the subsequent EVB simulations propagated in each direction towards the Michaelis complex and relevant intermediate or product states, respectively. In the case of the second step of each of the three stepwise mechanisms (**Figures 1A** to **C**), the intermediate state generated from modeling the initial nucleophilic attack on the carbonyl atom of the lactone ring was used as a starting point to model the subsequent ring opening reaction. All EVB simulations for each step were performed using 51 EVB mapping windows of 200 ps length per EVB trajectory, and 30 replicates were performed per system, leading to a total of 1.5 μs equilibration and 306 ns EVB sampling per system and mechanism (612 ns EVB sampling for the mechanisms comprising a 3-state process), and a total of 154.6 μs simulation time (both equilibration and subsequent EVB simulations) over all systems.

Finally, the EVB simulations were calibrated to corresponding simulations of the non-enzymatic hydrolysis of each AHL shown in **Figure S18**, proceeding through each of the four mechanisms shown in **Figure 2** of the main text. Equilibration and EVB simulations were performed as outlined for the enzymatic reaction, using the same simulation timescales, EVB parameters and simulation settings, with the exception of the fact that a slightly larger 1 kcal $mol^{-1}$ $Å^{-2}$ harmonic restraint was applied to the reacting atoms during the simulations. In the case of the bridging/terminal hydroxide and concerted mechanisms (**Figures 1A, B** and **D**), the reacting system was comprised of the relevant AHL and either a hydroxide ion or water molecule; in the case of the Asp mechanism (**Figure 1C**), we also included a propionate ion in the reacting system as a model for the side-chain of Asp122. We note here that in the case of the bridging/terminal hydroxide mechanisms, as the nucleophile is a metal bound hydroxide ion, it is important to also take into account the energetic cost of generating this hydroxide ion, which can be obtained from the relationship $\Delta G_{PT} = 2.303RT(pK_a(OH) - pH)$,[19] where R is the ideal gas constant, and T is the temperature. In the case of the bridging water molecule, the $pK_a$ of this water molecule can be estimated to be ~7 based on analogous systems,[20] and thus deprotonation of the water molecule to generate hydroxide would be spontaneous at physiological pH (7.5). In the case of the terminal water molecule, the $pK_a$ of an iron bound water is in the range of 9.3-9.5,[21, 22] but this does not take into account the impact of the enzyme environment on this $pK_a$. However, it provides a ballpark estimate for the expected free energy cost of deprotonating the terminal hydroxide ion, based on the equation above, and thus we have added a correction of 2.6 kcal mol$^{-1}$ to the energies of all stationary points throughout for this mechanism, as an estimate of the cost of generating this hydroxide ion at physiological pH.

#### S2. Supplementary Figures

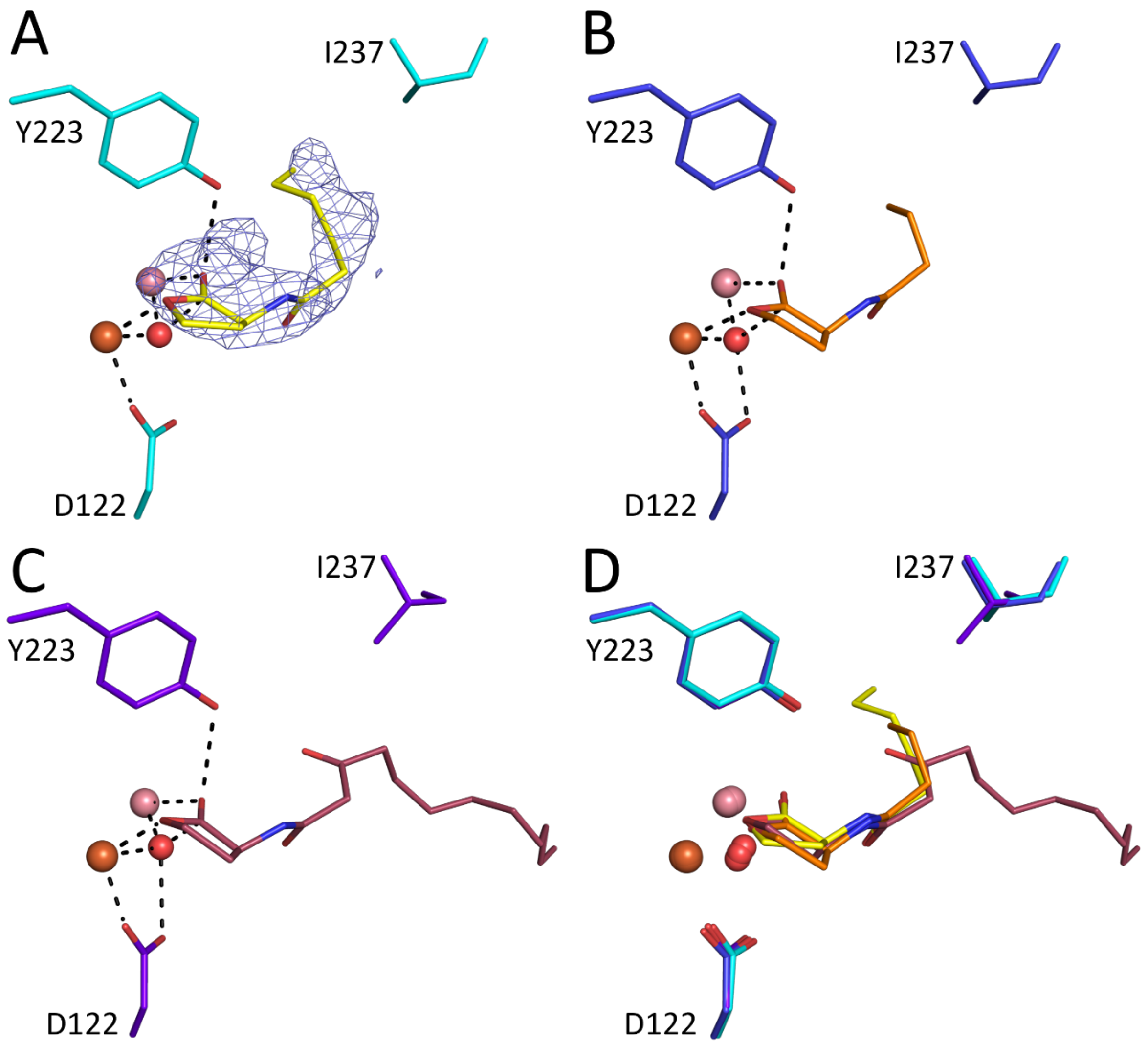


**Figure S1**. Comparison of the structure of GcL bound to C6-HSL (this work; 9AYT) and to C4- and 3-oxo-C12-HSL (prior work, PDB IDs: 6N9Q[23] and 6N9R[23]). (**A**) GcL (cyan) bound to C6-HSL (yellow; 9AYT; monomer D). Mesh shows the Fo-Fc omit map contoured at 2.2 σ, with a ligand occupancy of 0.80. (**B**) GcL (blue) bound to C4-HSL (orange). (**C**) GcL (purple) bound to 3-oxo-C12-HSL (red). (**D**) superimposition of the GcL active site with substrates bound: C4-HSL (orange), C6-HSL (yellow), and 3-oxo-C12-HSL (red).

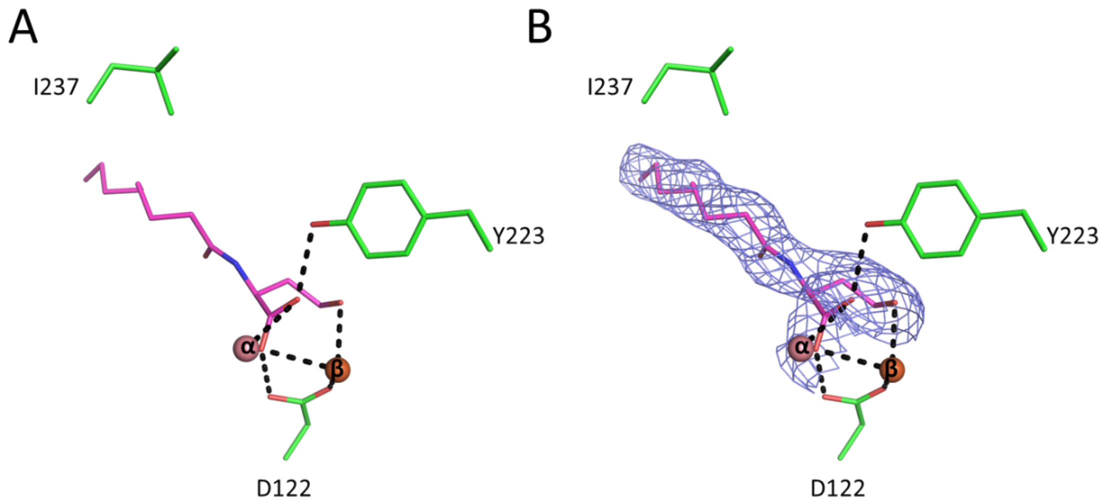


**Figure S2**. (**A**) Structure of GcL bound to the hydrolytic product for C8-HSL modeled at 0.7 occupancy (PDB ID: 9B2O). (**B**) Blue mesh shows the Fourier difference Fo-Fc omit map contoured at 2.2 σ.

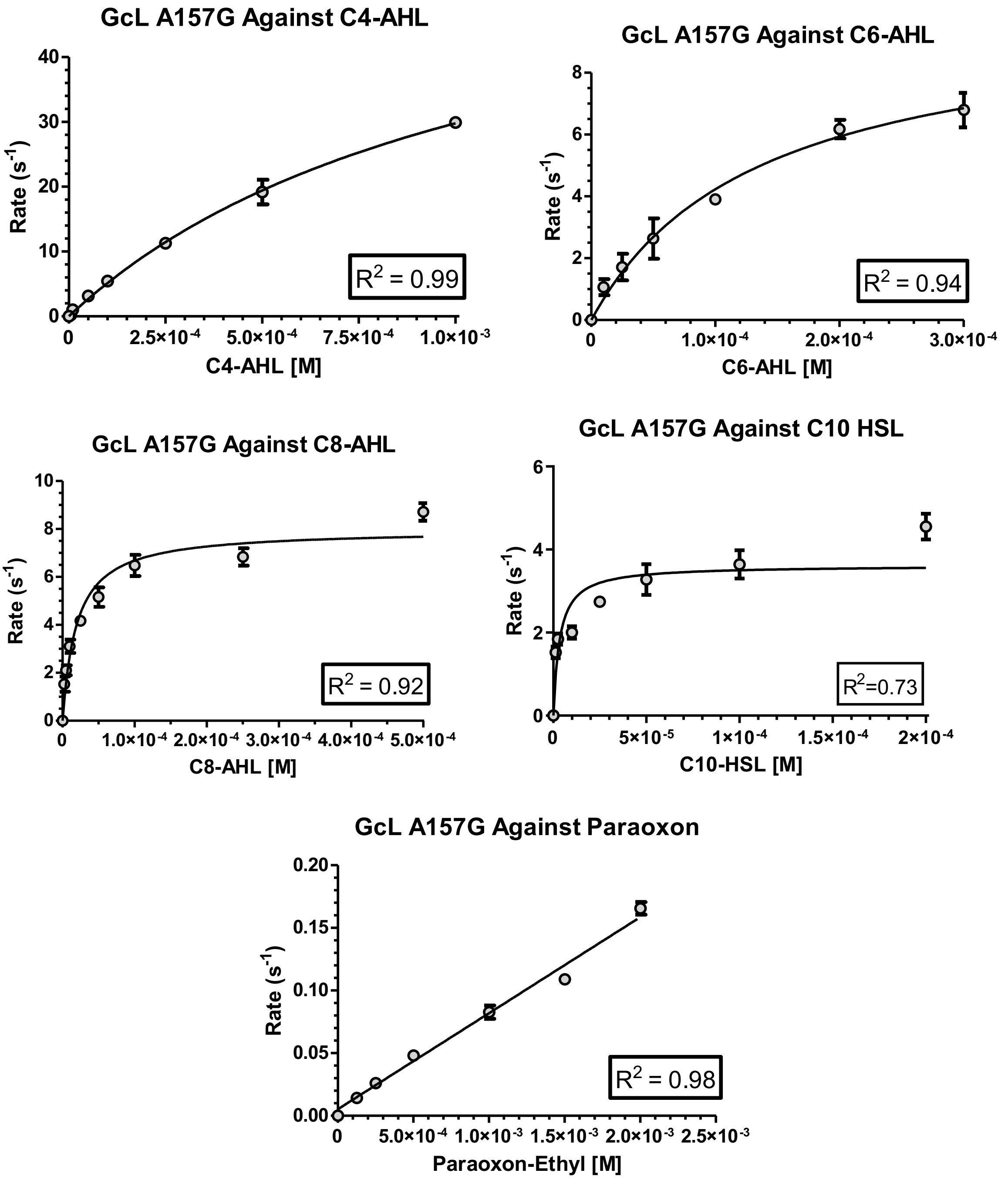


**Figure S3.** Fits for AHL substrates and paraoxon of GcL mutant A157G to the Michaelis Menten equation. For substrates which did not fit the Michaelis Menten equation, a linear regression was generated to determine catalytic efficiency.

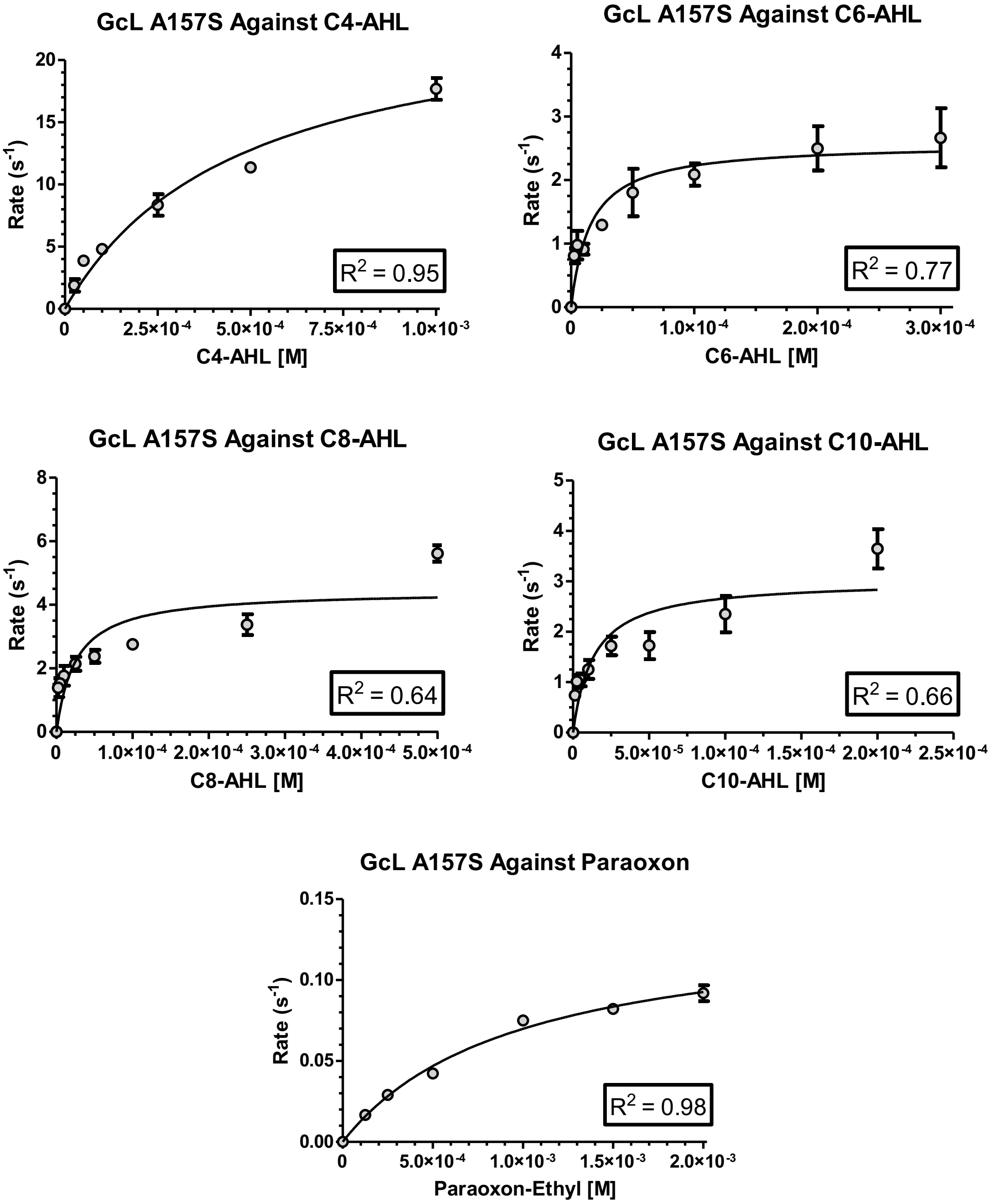


**Figure S4.** Fits for AHL substrates and paraoxon of GcL mutant A157S to the Michaelis Menten equation.

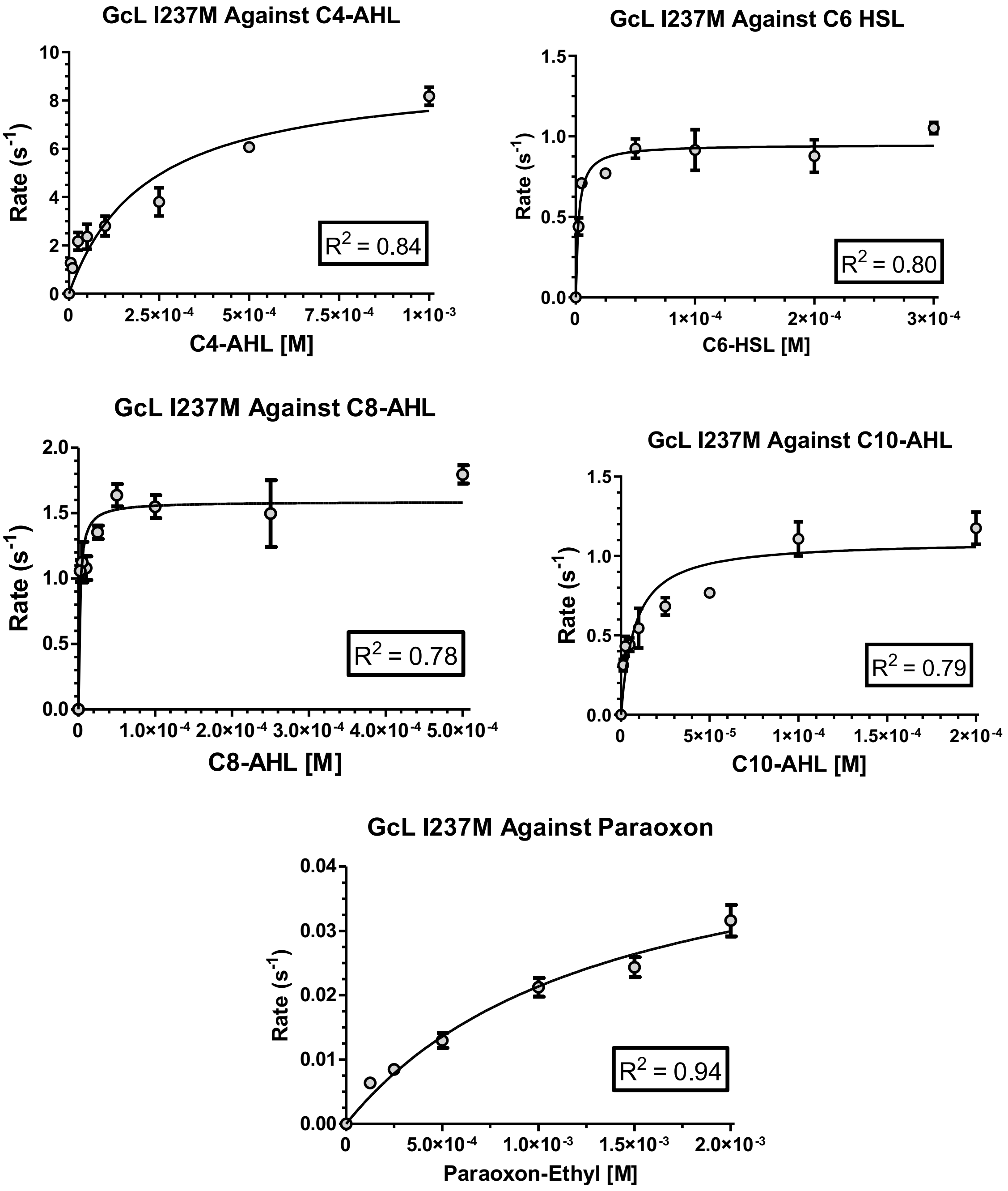


**Figure S5.** Fits for AHL substrates and paraoxon of GcL mutant I237M to the Michaelis Menten equation.

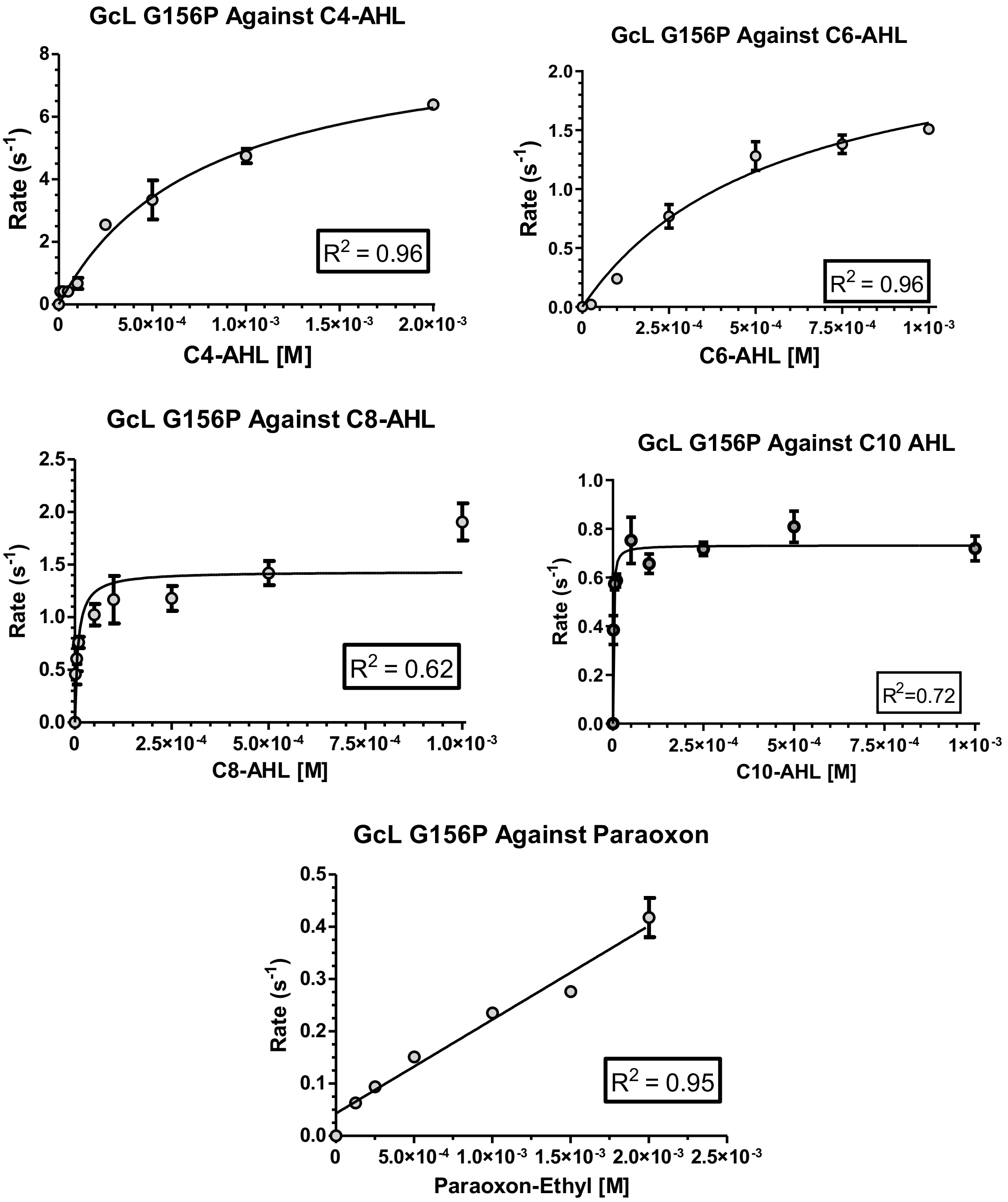


**Figure S6.** Fits for AHL substrates and paraoxon of GcL mutant A157G to the Michaelis Menten equation. For substrates which did not fit the Michaelis Menten equation, a linear regression was generated to determine catalytic efficiency.

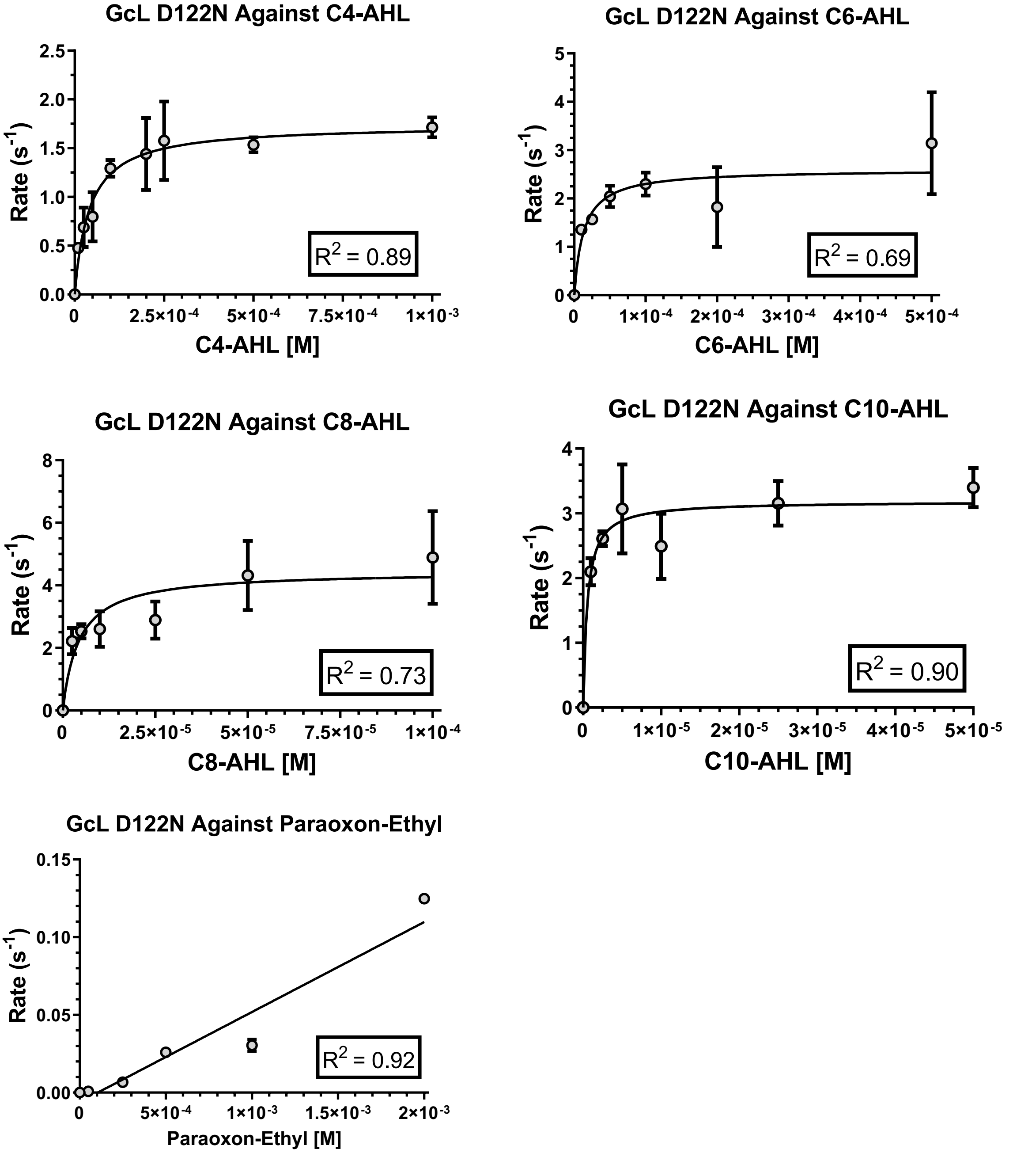


**Figure S7.** Fits for AHL substrates and paraoxon of GcL mutant D122N to the Michaelis Menten equation. For substrates which did not fit the Michaelis Menten equation, a linear regression was generated to determine catalytic efficiency.

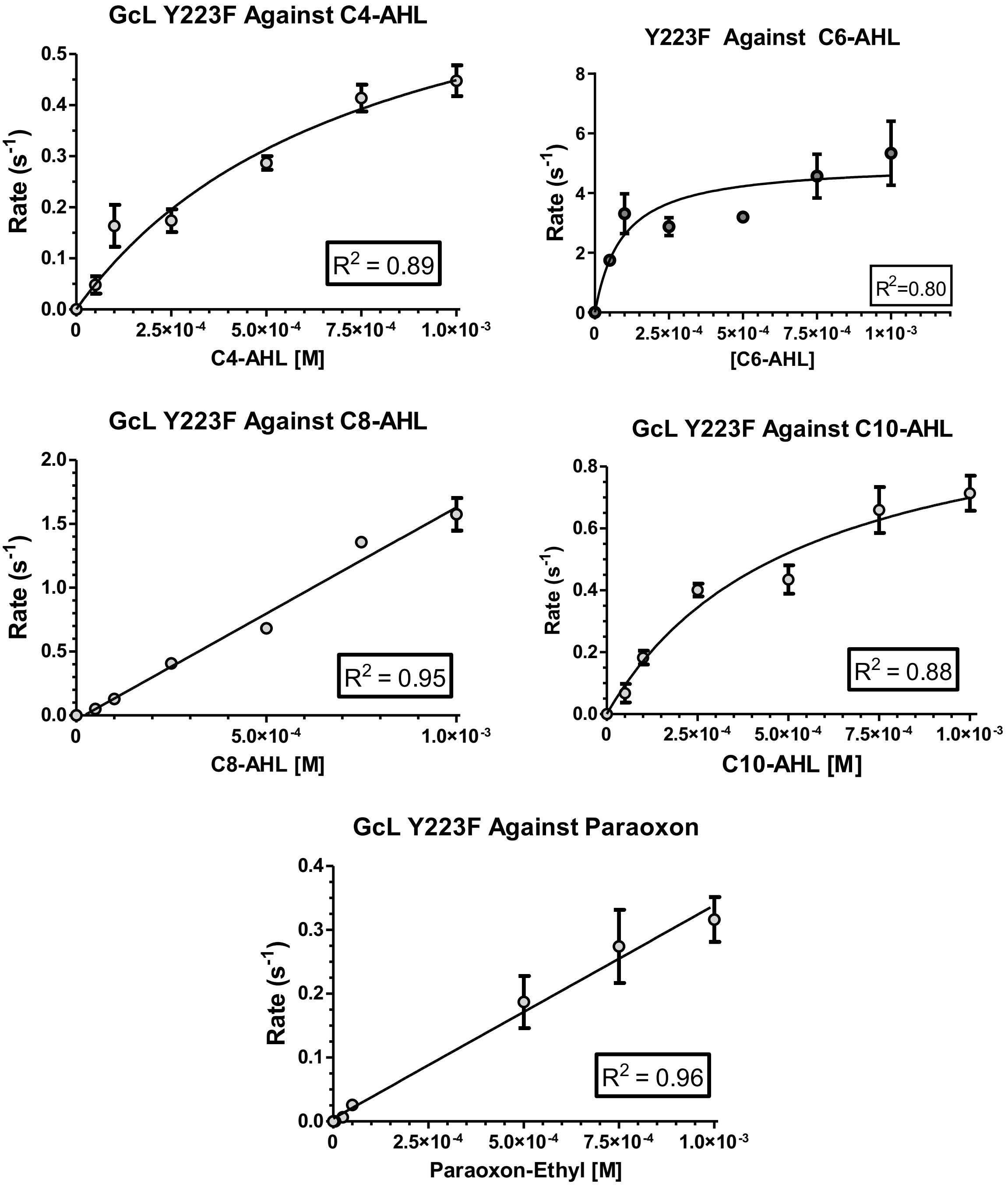


**Figure S8.** Fits for AHL substrates and paraoxon of GcL mutant Y223F to the Michaelis Menten equation. For substrates which did not fit the Michaelis Menten equation, a linear regression was generated to determine catalytic efficiency.

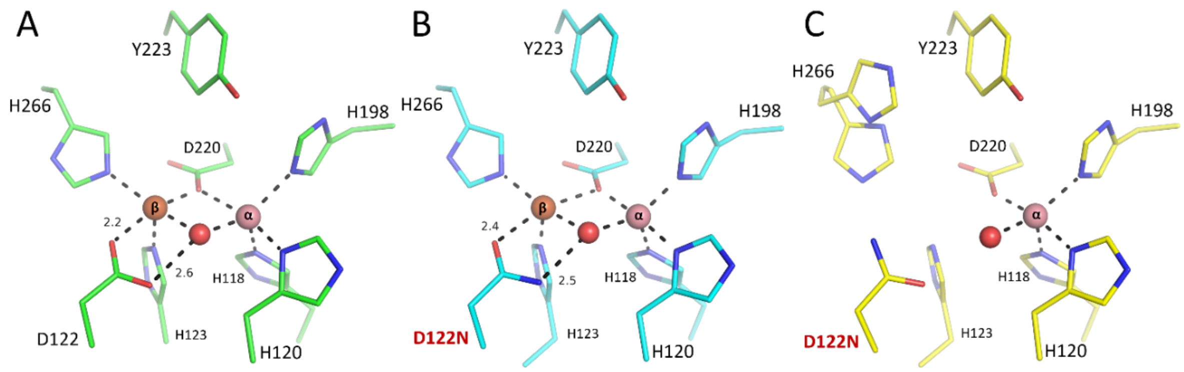


**Figure S9**. Overview of the obtained structures of the D122N GcL variant. Shown here for comparison is also (**A**) the structure of wild-type GcL (PDB ID: 6N9I,[23] green). The D122N variant was crystallized in two forms, (**B**) one with a complete bi-metallic center (PDB ID: 9B2L, this study, cyan, in H3 space group and in C2 space group (PDB ID: 9B2P; not shown)) and (**C**) another conformation where the β-metal has very low occupancy (not modelled, PDB ID: 9B2N, this study, yellow). Metal cations and active site water molecules are shown as spheres. Distances are indicated in Ångströms. The structure of the D122N variant with complete bi-metallic center crystallized in two different space groups, but the active site geometry is essentially the same.

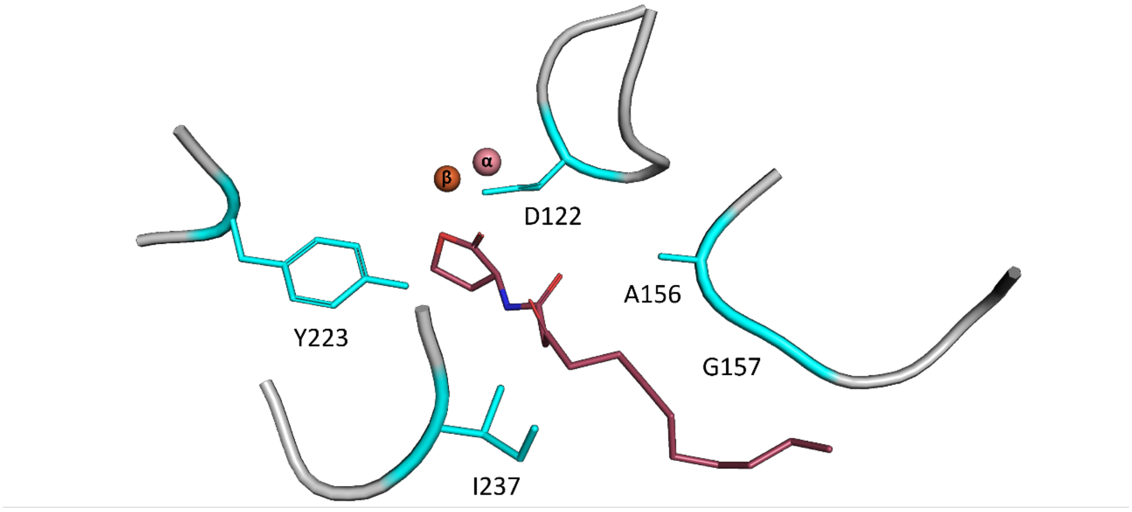


**Figure S10**. Active site of GcL bound to 3-oxo-dodecanoyl L-homoserine lactone (PDB ID: 6N9R), highlighting key binding residues. Residues mutated in this study are highlighted in cyan and are involved in ligand binding. Metals are shown in reduced scale, α: cobalt, β: iron.

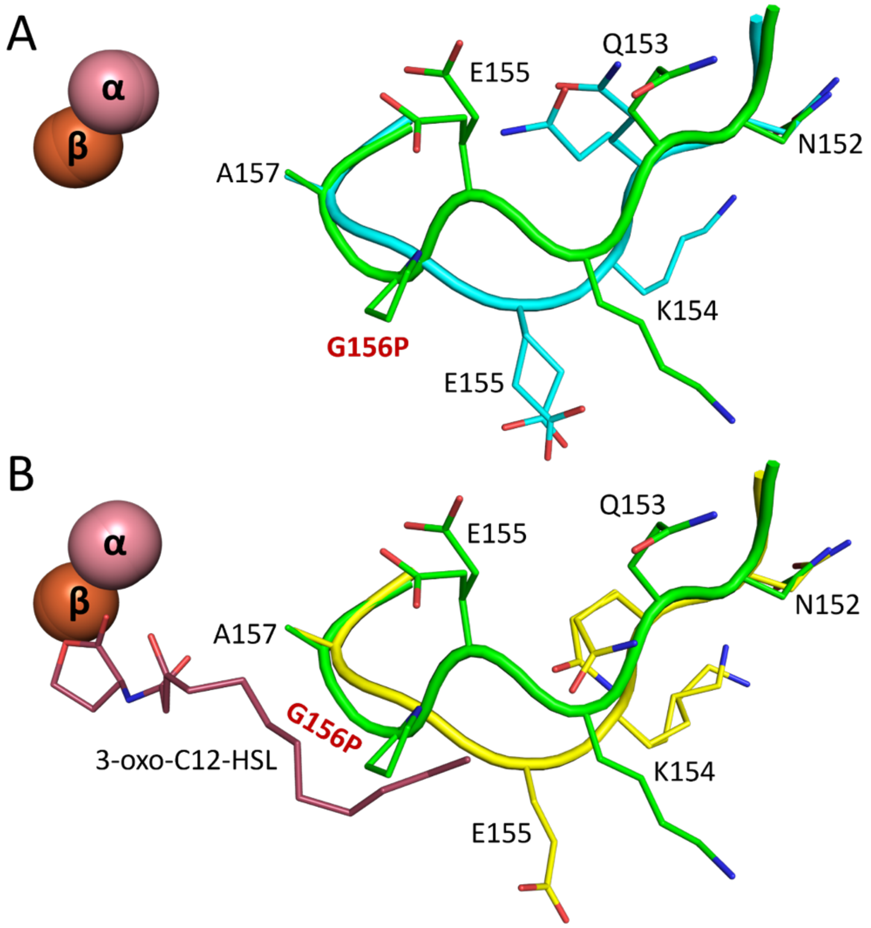


**Figure S11.** Alternate conformation of the Asn152-Ala157 loop in the Gly156Pro variant (PDB ID: 9B2I, this study). (**A**) GcL Gly156Pro (green) aligned with GcL WT (cyan – PDB: 6N9I[23]), showing the displacement of several residues and side-chains. (**B**) GcL Gly156Pro (green) aligned with GcL WT bound to 3-oxo-C12-HSL (yellow – PDB 6N9R[23]). The change in loop conformation has potential effects on substrate binding, as the proline side-chain is oriented into the AHL binding cleft, and restructured loop can alter interactions between other side-chain residues (such as Glu155) and AHL.

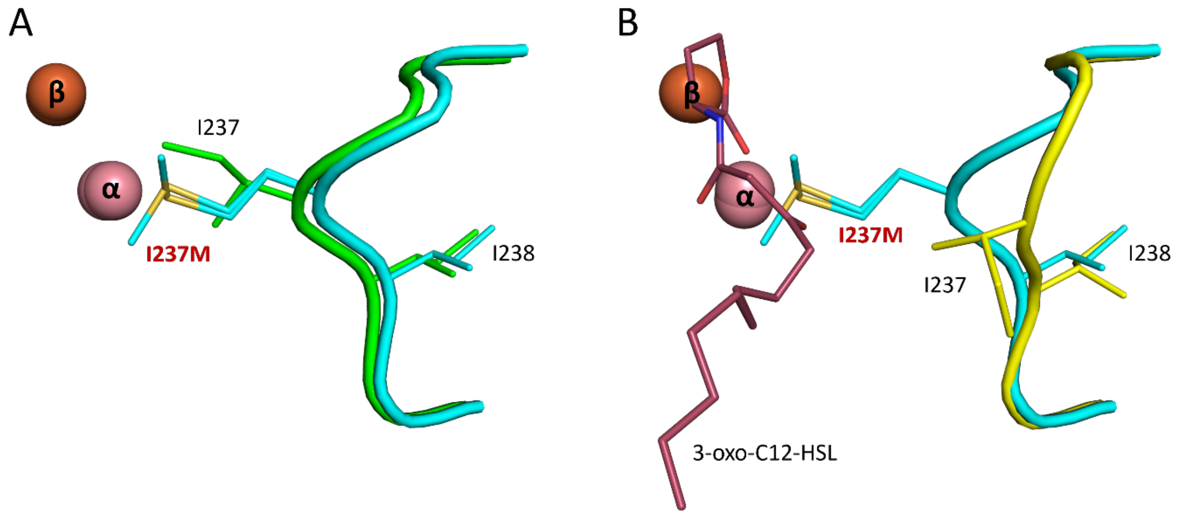


**Figure S12**. Comparison between GcL WT and GcL I237M variant, showing the alternative conformation of the loop consisting of Pro234-Asp240, between apo-GcL and substrate bound GcL. The Ile237Met residue must move to accommodate substrate binding. (**A**) GcL WT (PDB: 6N9I,[23] green) and the GcL Ile237Met variant (PDB: 9B2J, this study, cyan). (**B**) GcL WT (PDB: 6N9R,[23] yellow) bound to 3-oxo-C12-HSL (red). Active site metals are shown as spheres.

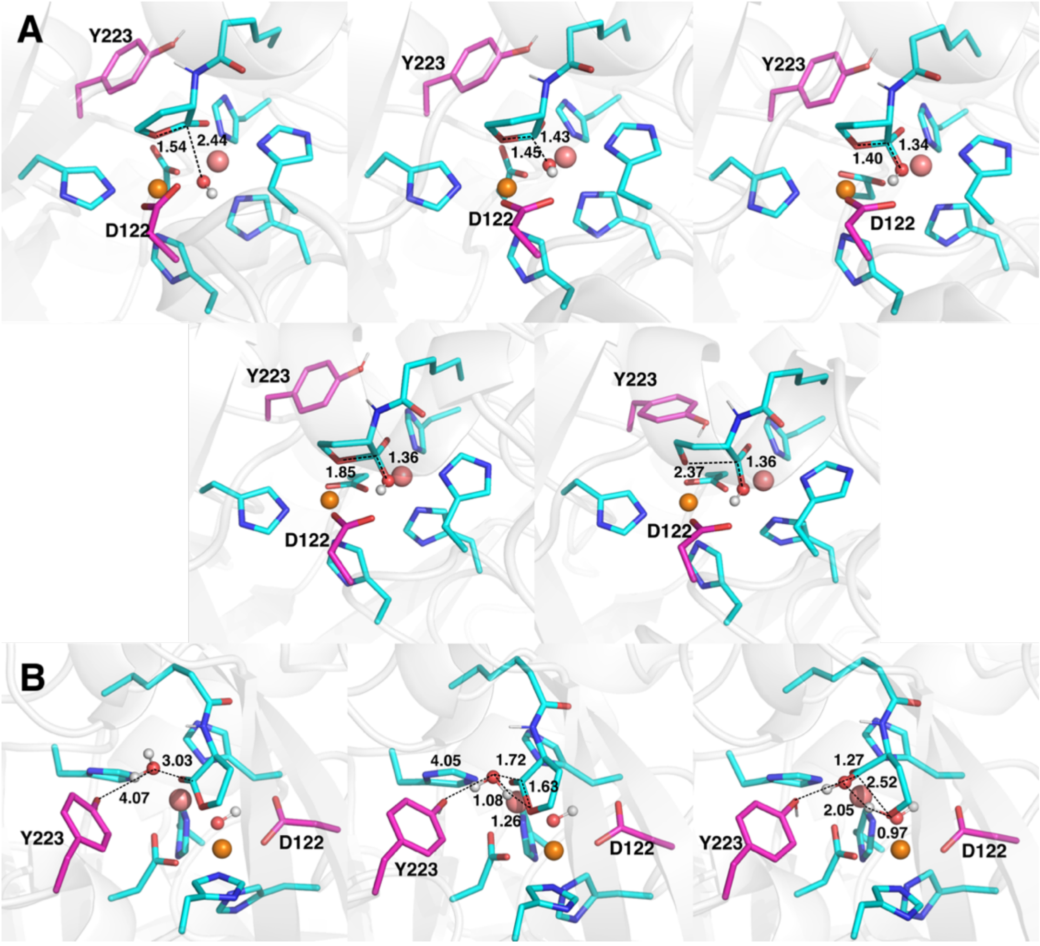


**Figure S13.** Representative structures of the Michaelis complexes, transition states, and intermediate states, for the hydrolysis of C6-HSL catalyzed by wild-type GcL *via* energetically unfavorable (**A**) bridging hydroxide and (**B**) concerted mechanisms (**Figures 1A** and **D**, and **Table 2** of the main text), as obtained from empirical valence bond simulations of these reactions. The structures shown here are the centroids of the top ranked cluster obtained from RMSD clustering, performed as described in the **Materials and Methods**. The distances labeled on this figure (Å) are averages at each stationary point over all the EVB trajectories (see **Table S2**, with the corresponding data for the non-enzymatic reaction shown in **Table S3**, and metal-metal distances shown in **Table S4**). Highlighted here are the substrate, nucleophilic water, bridging hydroxide, $Fe^{2+}$ (brown), $Co^{2+}$ (salmon), and key catalytic residues. The corresponding data for the terminal hydroxide and Asp mechanisms are shown in **Figure 3**.

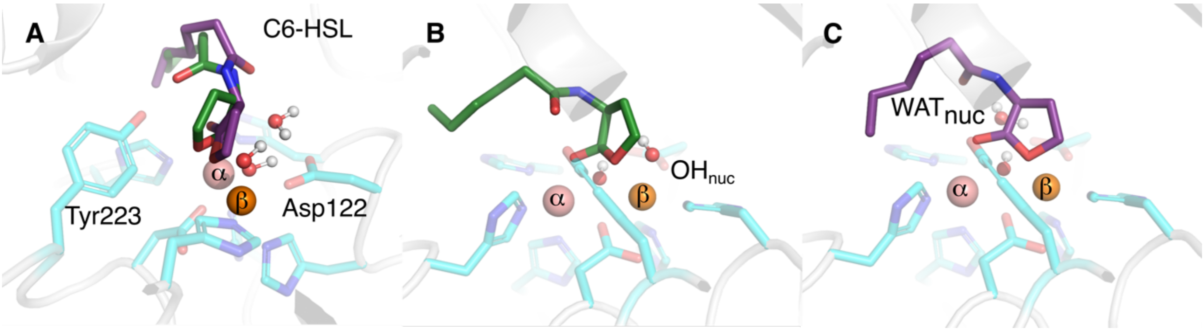


**Figure S14**. (**A**) Overlay of the starting structures for empirical valence bond (EVB) simulations of the terminal hydroxide **(green)** and Asp **(purple)** mechanisms **(Figure 1B** and **C)**, and (**B, C**) the individual starting structures for each mechanism, respectively. The nucleophilic hydroxide ion and water molecule are shown as $OH_{nuc}$ and $WAT_{nuc}$ respectively. For details, see the **Materials and Methods**.

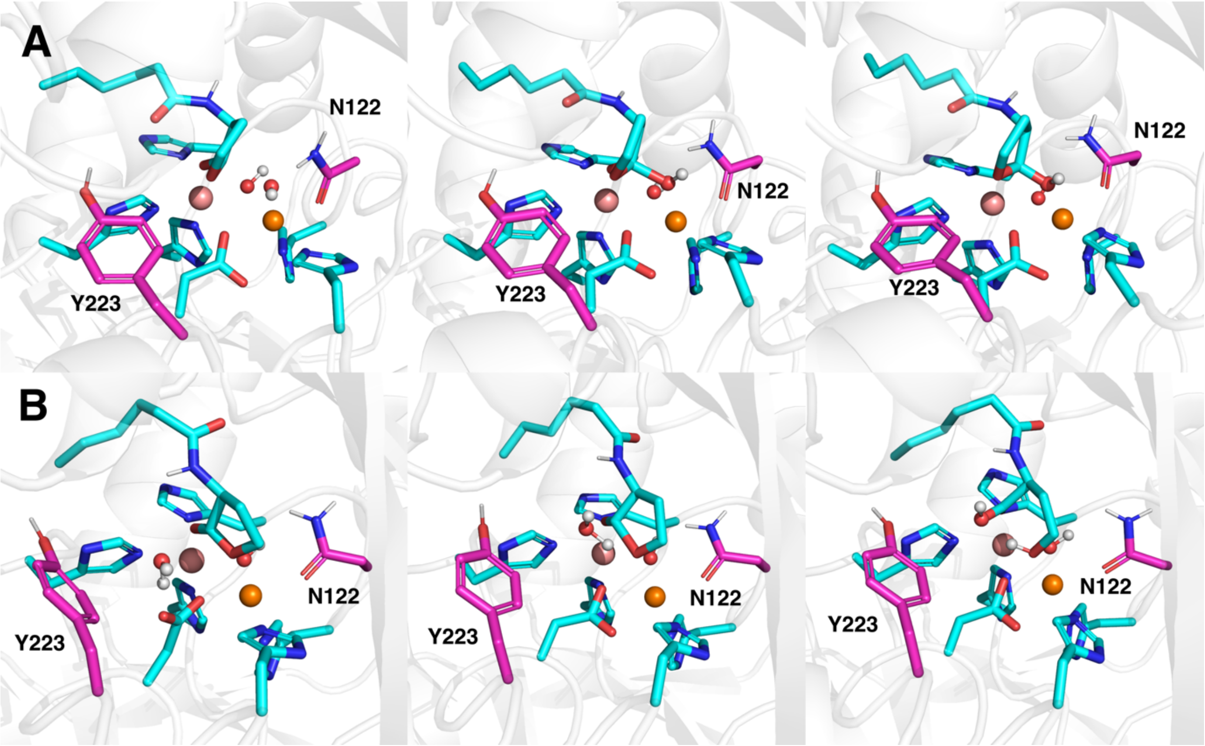


**Figure S15**. Representative structures of the Michaelis complexes, first transition state, and intermediate/product states, for the hydrolysis of C6-HSL catalyzed by D122N GcL *via* the (**A**) terminal hydroxide and (**B**) concerted mechanisms, as obtained from empirical valence bond (EVB) simulations of these reactions. The structures shown here are the centroids of the top ranked cluster obtained from RMSD clustering, performed as described in the **Materials and Methods**. Highlighted here are the substrate, nucleophilic water, bridging hydroxide, $Fe^{2+}$ (brown), $Co^{2+}$ (salmon), and the side-chains of key catalytic residues.

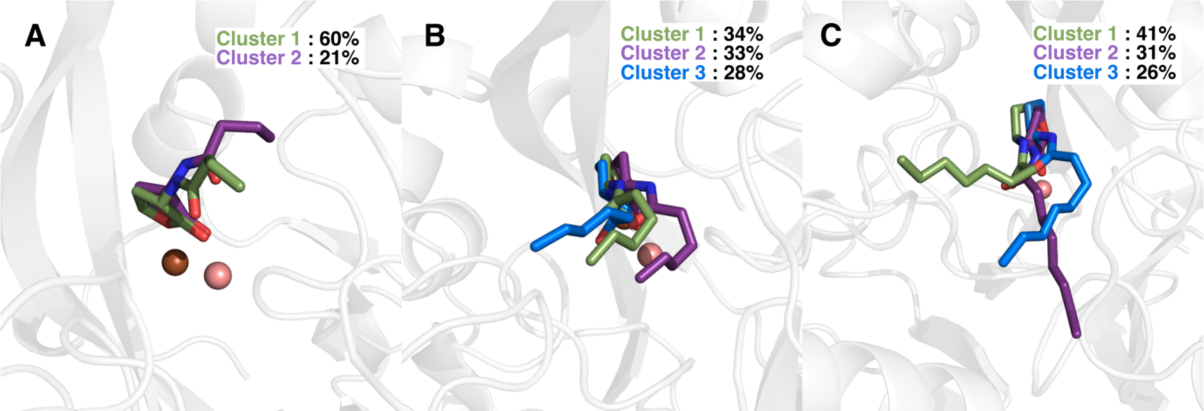


**Figure S16.** Overview of the main principal clusters (only shown clusters accounting for more than 10% of the simulation time) from simulations of (**A**) C4-HSL, (**B**) C6-HSL and (**C**) C8-HSL bound to wild-type GcL. The clustering was performed based on pairwise RMSD calculations over all atoms of the ligand using CPPTRAJ,[24] as described in the **Materials and Methods**.

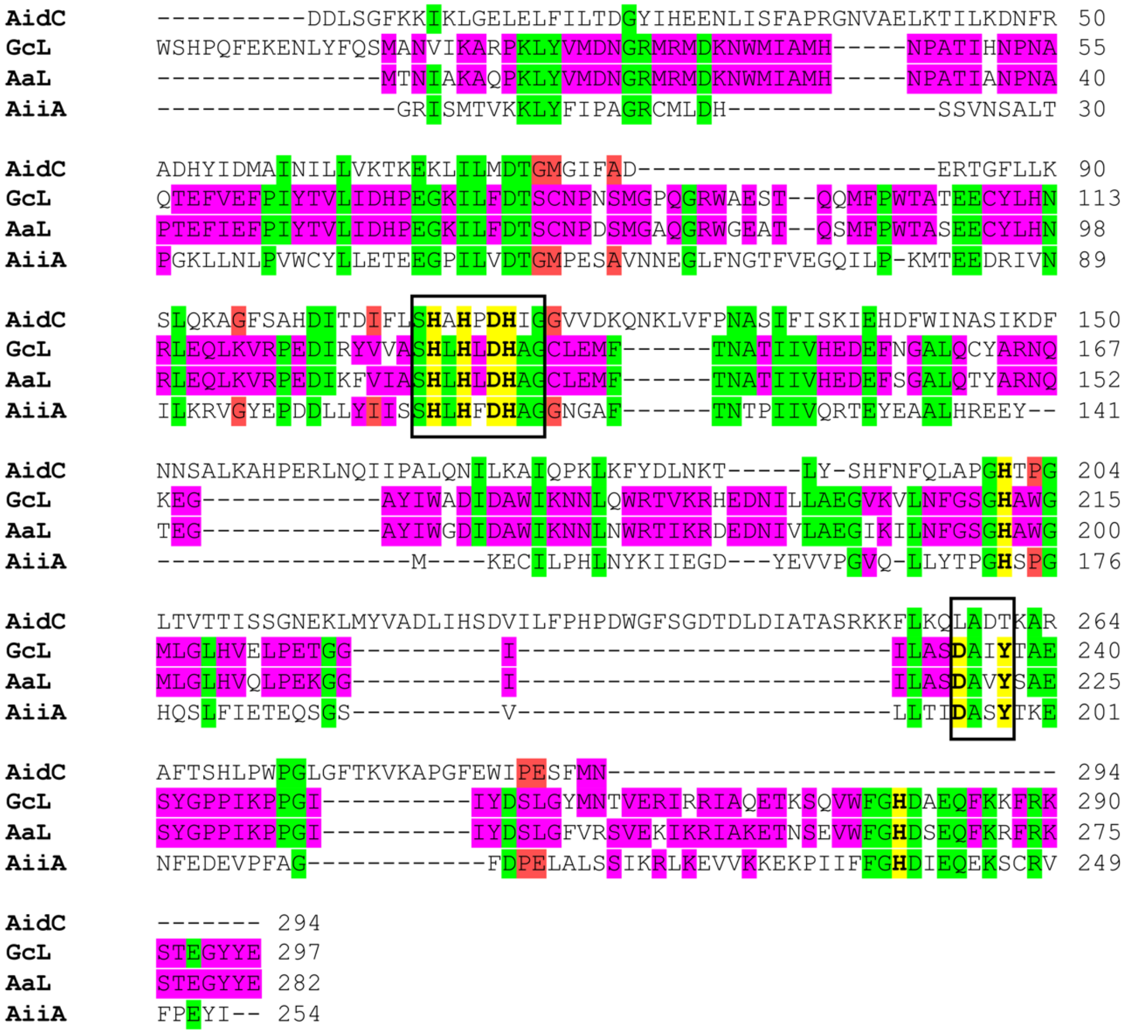


**Figure S17.** Sequence alignment of four members of the metallo-β-lactamase-like lactonase (MLLs) family, AidC (PDB ID: 4ZO3[25]), GcL (PDB ID: 6N9I[23]), AaL (PDB ID: 6CGZ[26]) and AiiA (PDB ID: 3DHB[27]). Residues coordinating the bimetallic center in the binding domain are highlighted in bold, as well as the binding domain Tyr223. Residues conserved in three or four sequences are highlighted in green, while residues conserved in two sequences are highlighted in pink, for clarity. Sequence alignment was performed using the web interface of Clustal Omega.[28]

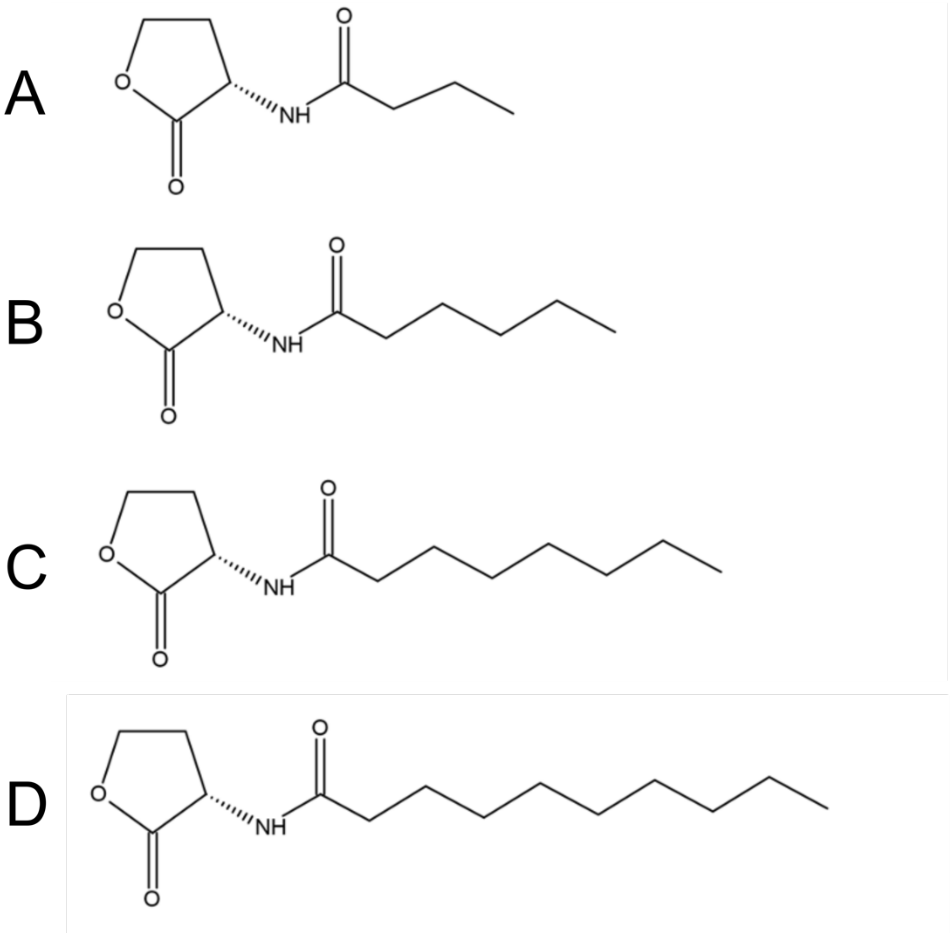


**Figure S18.** Schematic representation of the *N*-acyl-L-homoserine lactone (AHL) substrates studied in this work. (**A**) C4-HSL, (**B**) C6-HSL, (**C**) C8-HSL, and (**D**) C10-HSL.

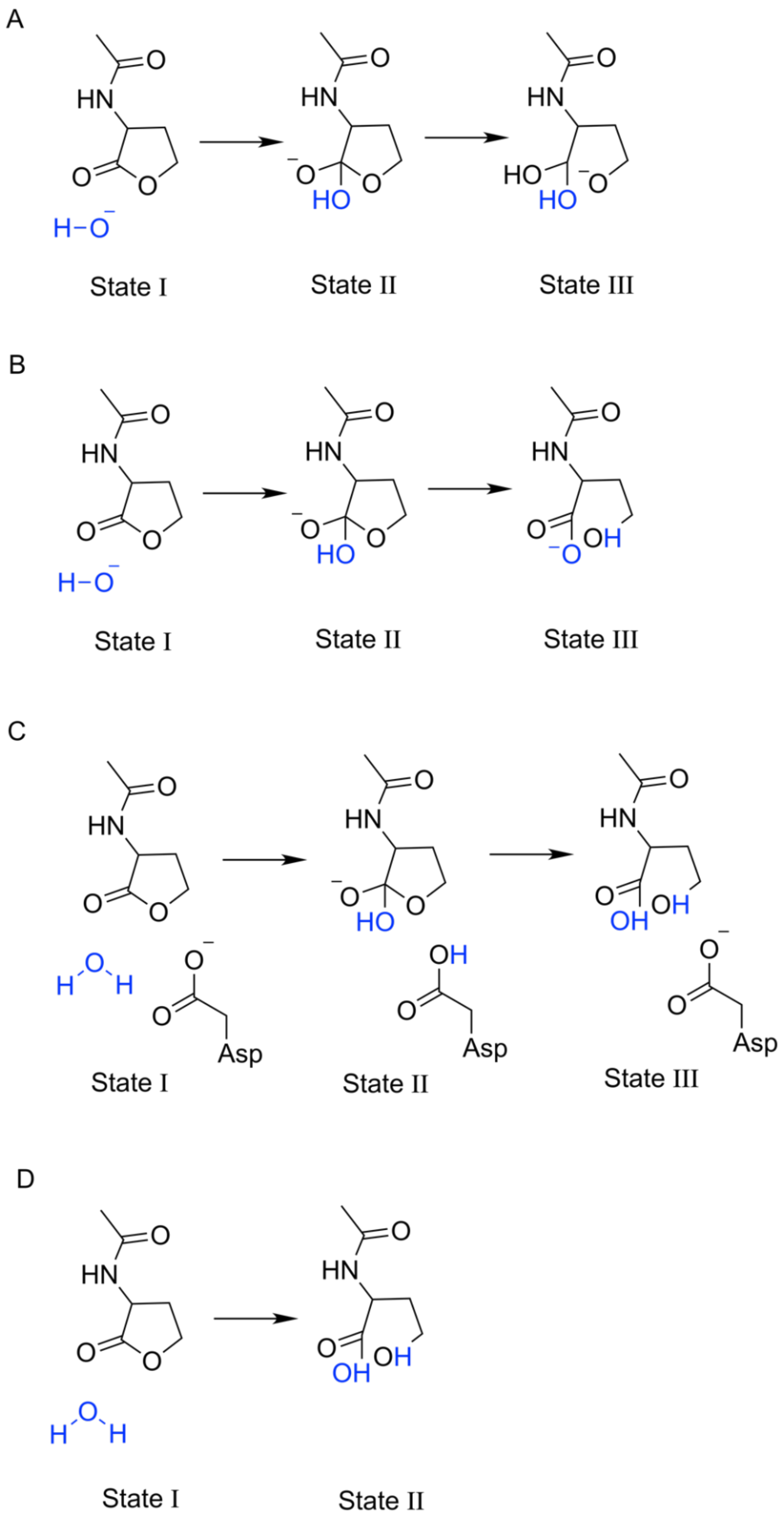


**Figure S19**. The valence bond (VB) states used to describe each of the mechanisms studied in this work (**Figure 1**). The corresponding EVB parameters are provided on Zenodo, DOI: 10.5281/zenodo.11072674.

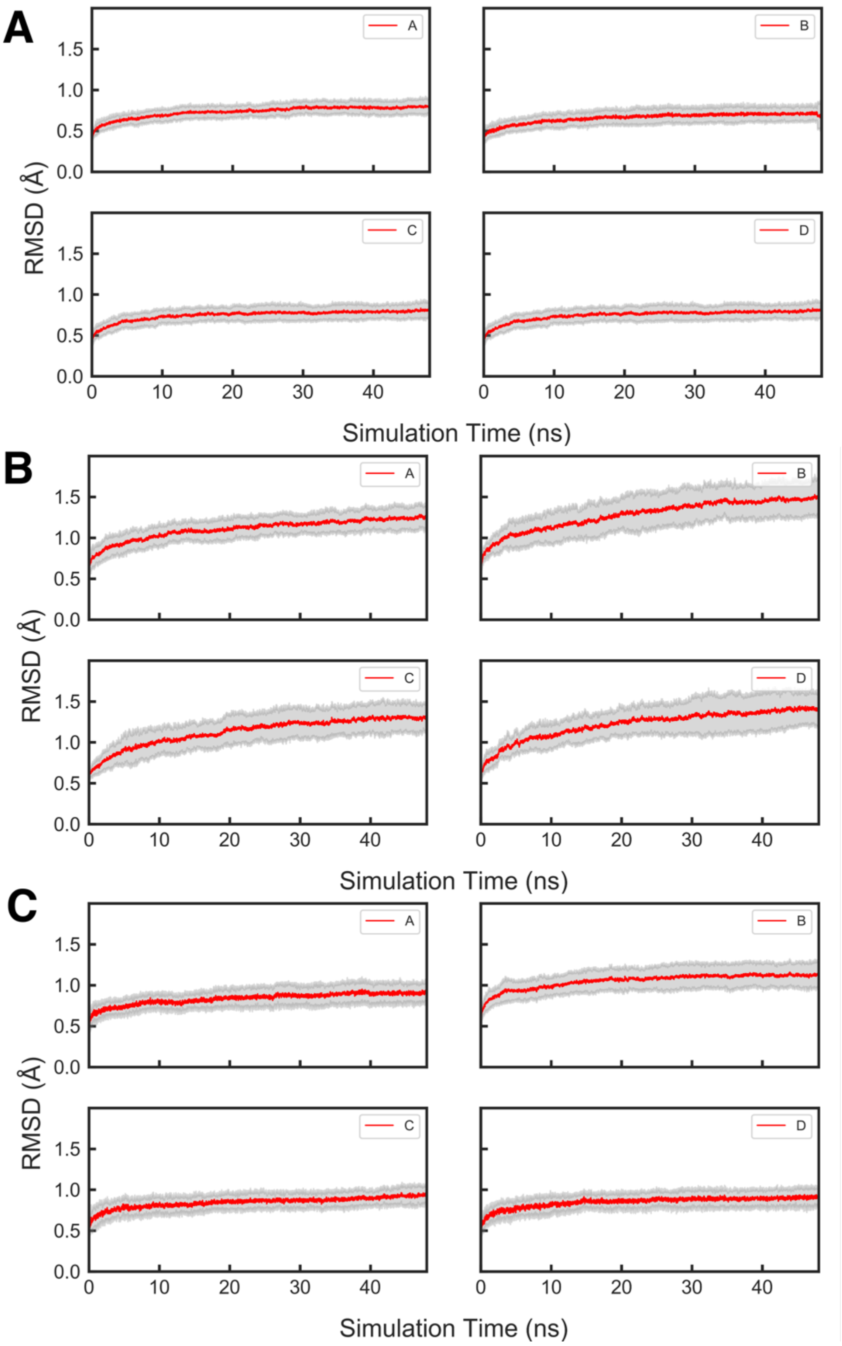


**Figure S20.** Root mean square deviations (RMSD, Å) of all backbone heavy atoms from EVB equilibration simulations of the hydrolysis of C6-HSL through the four studied mechanisms, “bridging hydroxide mechanism”, “terminal hydroxide mechanism”, “Asp mechanism” and “concerted mechanism”, as catalyzed by wild-type (**A**) GcL, (**B**) AiiA and (**C**) AaL. Data was collected every 10 ps from 30 replicas of 50 ns length each. The solid lines show rolling averages of the RMSD over all 30 replicas, and the shaded regions show the corresponding standard deviations in these values

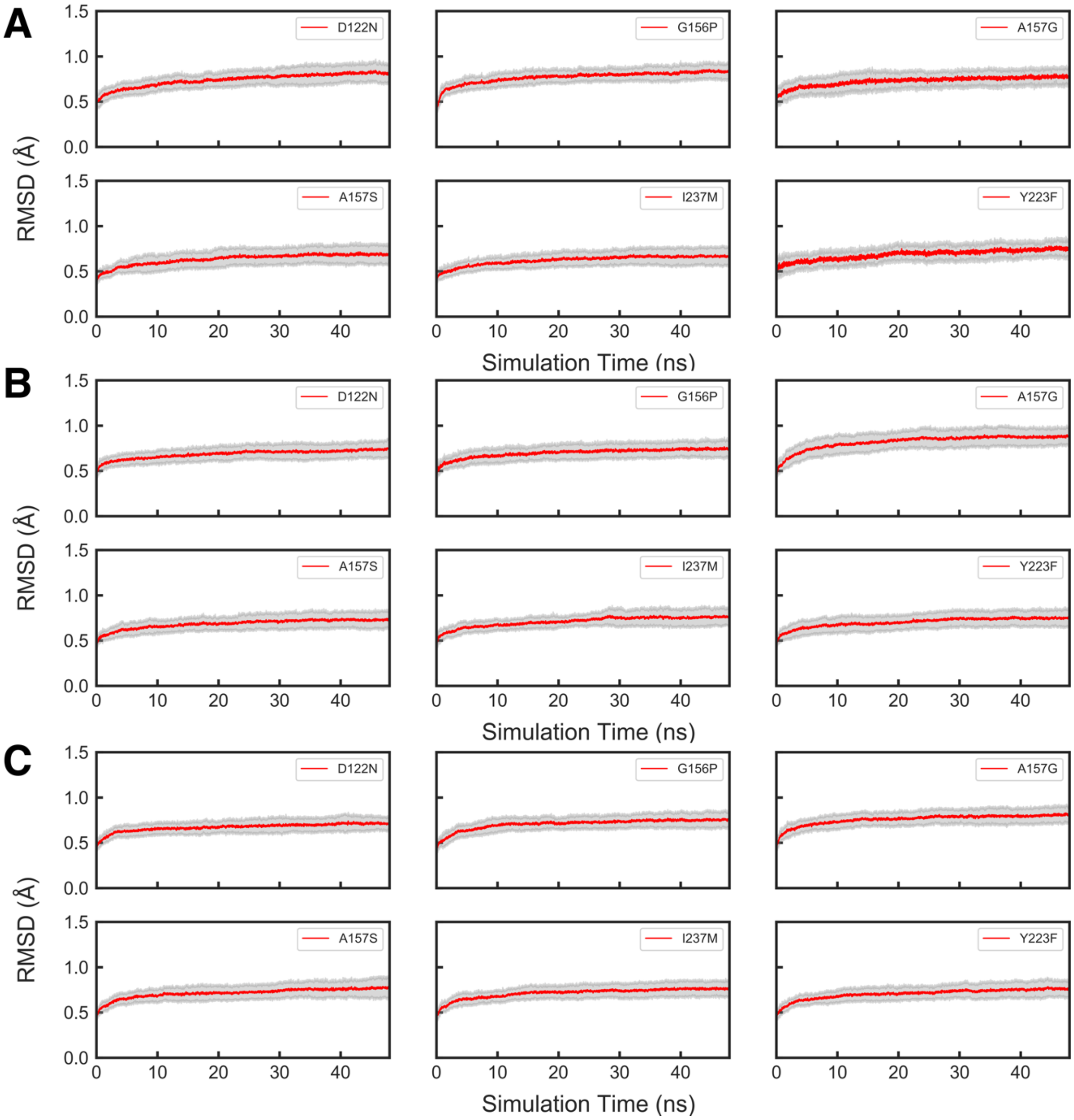


**Figure S21.** Root mean square deviations (RMSD, Å) of all backbone heavy atoms from EVB equilibration simulations of the hydrolysis of (**A**) C4-HSL, (**B**) C6-HSL and (**C**) C10-HSL as catalyzed by Asp122Asn, Gly156Pro, Ala157Gly, Ala157Ser, Ile237Met and Tyr223Phe GcL variants through the terminal hydroxide mechanism. Data was collected every 10 ps from 30 replicas of 50 ns length each. The solid lines show rolling averages of the RMSD over all 30 replicas, and the shaded regions show the corresponding standard deviations in these values.

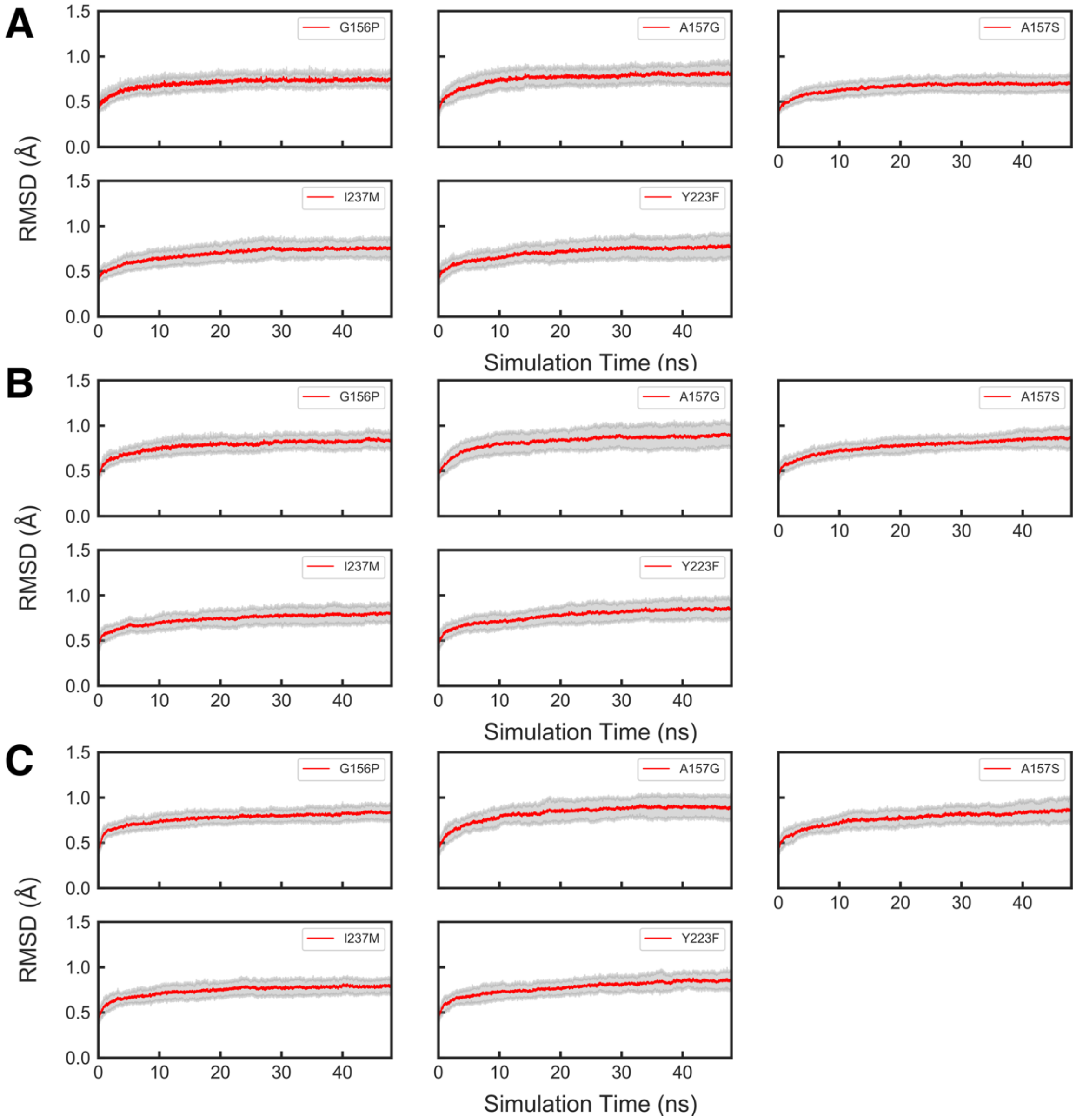


**Figure S22.** Root mean square deviations (RMSD, Å) of all backbone heavy atoms from EVB equilibration simulations of the hydrolysis of (**A**) C4-HSL, (**B**) C6-HSL and (**C**) C10-HSL as catalyzed by Gly156Pro, Ala157Gly, Ala157Ser, Ile237Met and Tyr223Phe GcL variants through the Asp mechanism. Data was collected every 10 ps from 30 replicas of 50 ns length each. The solid lines show rolling averages of the RMSD over all 30 replicas, and the shaded regions show the corresponding standard deviations in these values.

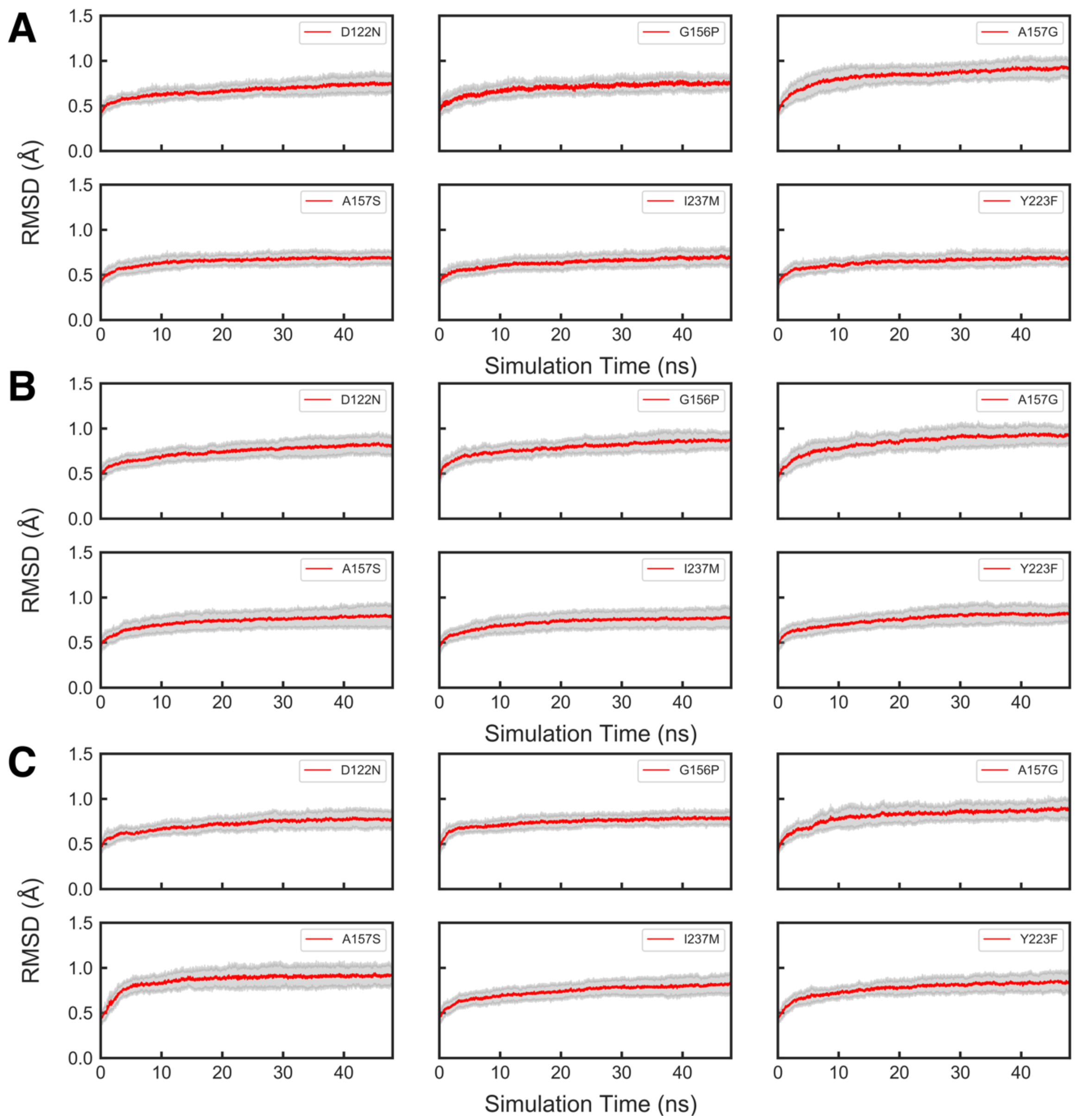


**Figure S23**. Root mean square deviations (RMSD, Å) of all backbone heavy atoms from EVB equilibration simulations of the hydrolysis of (**A**) C4-HSL, (**B**) C6-HSL and (**C**) C10-HSL as catalyzed by Asp122Asn, Gly156Pro, Ala157Gly, Ala157Ser, Ile237Met and Tyr223Phe GcL variants through the concerted mechanism. Data was collected every 10 ps from 30 replicas of 50 ns length each. The solid lines show rolling averages of the RMSD over all 30 replicas, and the shaded regions show the corresponding standard deviations in these values.

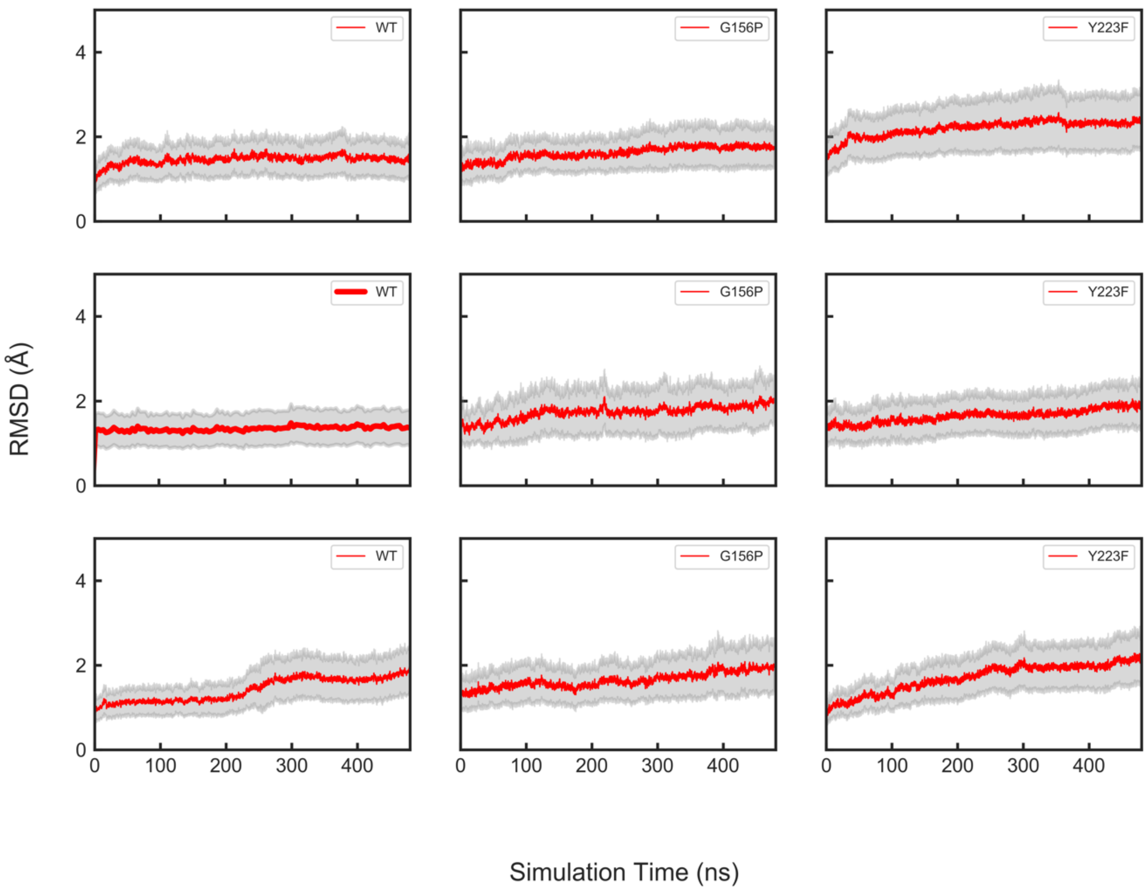


**Figure S24**. Root mean square deviations (RMSD, Å) of all backbone heavy atoms from MD simulations of wild-type, Gly156Pro and Tyr223Phe GcL variants bound to (top row) C4-HSL, (middle row) C6-HSL and (bottom row) C8-HSL. Data was collected every 50 ps from 3 replicas of 500 ns length each. The solid lines show rolling averages of the RMSD over all 3 replicas, and the shaded regions show the corresponding standard deviations in these values.

### S3. Supplementary Tables

**Table S2.** Data collection and refinement statistics.[a]

| | G156P | I237M | D122N (bi-metal) | D122N (bi-metal) | D122N (mono metal) | Bound to C6 HSL | Bound to C8 HSL reaction product |
|---|---|---|---|---|---|---|---|
| **PDB ID** | 9B2I | 9B2J | 9B2L | 9B2P | 9B2N | 9AYT | 9B2O |
| **Resolution (Å)** | 2.35 | 2.3 | 2.2 | 2.25 | 1.9 | 2.1 | 1.85 |
| **diffraction source** | APS 23ID-B | APS 23ID-B | APS 23IDD | APS 23IDD | APS 23ID-B | APS 23ID-B | APS 23IDD |
| **wavelength [Å]** | 0.99184 | 0.99184 | 1.03333 | 1.03333 | 1.03317 | 1.033199 | 1.03321 |
| **detector** | EIGER | EIGER | PILATUS | PILATUS | MARCCD300 | EIGER | PILATUS |
| **rotation range per image [º]** | 0.5 | 0.5 | 0.2 | 0.2 | 1 | 0.2 | 0.3 |
| **total rotation range [º]** | 220 | 220 | 220 | 220 | 250 | 220 | 240 |
| **Space Group** | C2 | C2 | H3 | C2 | C2 | H3 | C2 |
| **Unit cell parameters [Å]** | a=145.8, b=108.2, c=78.8 | a=147.4, b=110.3, c=79.4 | a=b=109.2, c=221.6 | a=160.8, b=109.1, c=97.2 | a=146.4, b=109.1, c=78.7 | a=b=108.5, c=228.5 | a=161.36, b= 109.81, c=97.37 |
| **[º]** | α= γ=90, β=116.1 | α=γ=90.0, β=115.9, γ=90.0 | α=β=90, γ=120 | α=γ=90, β=116.4 | α= γ=90, β=115.9 | α=β=90, γ=120 | α=γ=90, β=116.346 |
| **resolution range [Å]** | 2.35 (2.35 – 2.45) | 2.3 (2.3 – 2.4) | 2.2 (2.2 – 2.3) | 2.25 (2.25-2.35) | 1.9 (1.9 – 2.0) | 2.1 (2.1 - 2.2) | 1.85 (1.85-1.95) |
| **# of reflections (last bin)** | 179203 (20447) | 180550 (48263) | 310320 (39599) | 286795 (35543) | 359785 (51013) | 365720 (48653) | 573602 (80814) |
| **# of unique reflections (last bin)** | 44722 (5250) | 19707 (5628) | 49981 (6245) | 70302 (8539) | 87022 (12356) | 58454 (7610) | 124206 (17767) |
| **completeness % (last bin)** | 97.6 (97.8) | 94.9 (93.1) | 99.8 (99.8) | 98.4 (98.6) | 99.3 (99.5) | 99.8 (99.9) | 95.8 (94.1) |
| **redundancy (last bin)** | 4.00 (3.89) | 3.55 (3.50) | 6.21 (6.34) | 4.08 (4.16) | 4.13 (4.13) | 6.26 (6.39) | 4.62 (4.55) |
| **⟨I/σ(I)⟩ (last bin)** | 13.51 (2.61) | 21.20 (4.59) | 10.58 (3.19) | 12.49 (2.44) | 17.41 (2.30) | 9.75 (2.33) | 14.19 (2.86) |
| **$R_{meas}$ % (last bin)** | 6.3 (61.8) | 4.8 (38.3) | 10.2 (61.2) | 7.8 (70.6) | 6.4 (79.7) | 11.1 (78.2) | 6.3 (64.0) |
| **$CC_{1/2}$ (last bin)** | 99.8 (87.9) | 99.8 (98.0) | 99.8 (91.8) | 99.8 (92.5) | 99.9 (87.9) | 99.6 (85.3) | 99.9 (92.7) |

| | *Refinement statistics* | | | | | | |
|---|---|---|---|---|---|---|---|
| **$R_{free}$ / $R_{work}$** | 22.06/19.06 | 19.92/18.44 | 24.15/21.41 | 23.37/20.93 | 22.3/21.3 | 21.90/19.27 | 21.48/18.67 |
| **# total model atoms** | 7559 | 7876 | 4669 | 6945 | 7490 | 5068 | 7155 |
| **% Ramachandran Favored** | 86.0 | 87.5 | 87.9 | 88.4 | 87.8 | 86.4 | 88.8 |
| **% Ramachandran Allowed** | 13.7 | 12.1 | 11.6 | 11.2 | 11.8 | 13.2 | 10.8 |
| **% Ramachandran Outliers** | 0.3 | 0.4 | 0.4 | 0.4 | 0.4 | 0.4 | 0.4 |
| ***RMSD from ideal*** | | | | | | | |
| **Bond lengths [Å]** | 0.001 | 0.004 | 0.002 | 0.001 | 0.002 | 0.002 | 0.012 |
| **Bond angles [º]** | 0.733 | 0.923 | 0.764 | 0.686 | 0.775 | 0.607 | 1.702 |

[a] Ramachandran plots were calculated with PROCHECK.[29]

**Table S2**. Reacting distances (Å) at key stationary points obtained from EVB simulations of the hydrolysis of C6-HSL by wild-type GcL.[a]

| Distance | Crystal MC/PS | RS | TS1 | IS | TS2 | PS |
|---|---|---|---|---|---|---|
| **Bridging Hydroxide Mechanism** | | | | | | |
| $O_{nuc}\ldots C$ | - | 2.44 ± 0.01 | 1.43 ± 0.00 | 1.34 ± 0.00 | 1.36 ± 0.00 | 1.36 ± 0.00 |
| $C\ldots O_{R}$ | **1.44/2.58** | 1.54 ± 0.00 | 1.45 ± 0.00 | 1.40 ± 0.00 | 1.85 ± 0.00 | 2.37 ± 0.01 |
| $O_{Tyr}\ldots O_{R}$ | **3.51/3.60** | 4.29 ± 0.03 | 4.24 ± 0.03 | 4.28 ± 0.03 | 4.08 ± 0.03 | 3.69 ± 0.04 |
| $O_{Tyr}\ldots O$ | **2.87/3.50** | 3.96 ± 0.02 | 3.71 ± 0.02 | 3.68 ± 0.03 | 3.64 ± 0.03 | 3.37 ± 0.03 |
| **Terminal Hydroxide Mechanism** | | | | | | |
| $O_{nuc}\ldots C$ | - | 2.16 ± 0.01 | 1.55 ± 0.00 | 1.34 ± 0.00 | 1.30 ± 0.00 | 1.26 ± 0.00 |
| $C\ldots O_{R}$ | **1.44/2.58** | 1.54 ± 0.00 | 1.43 ± 0.00 | 1.39 ± 0.00 | 1.72 ± 0.00 | 2.45 + 0.01 |
| $O_{nuc}\ldots H_{nuc}$ | - | 0.97 ± 0.00 | 0.96 ± 0.00 | 0.97 ± 0.00 | 1.18 ± 0.00 | 2.09 ± 0.00 |
| $H_{nuc}\ldots O_{R}$ | - | 2.69 ± 0.01 | 2.62 ± 0.01 | 2.60 ± 0.01 | 1.14 ± 0.00 | 0.96 ± 0.00 |
| $O_{Tyr}\ldots O_{R}$ | **3.51/3.60** | 3.32 ± 0.04 | 3.22 ± 0.04 | 3.19 ± 0.05 | 3.35 ± 0.05 | 3.22 ± 0.05 |
| $O_{Tyr}\ldots O$ | **2.87/3.50** | 4.52 ± 0.04 | 4.45 ± 0.05 | 4.30 ± 0.05 | 4.42 ± 0.05 | 4.55 ± 0.05 |
| **Asp Mechanism** | | | | | | |
| $O_{nuc}\ldots C$ | - | 2.58 ± 0.06 | 1.49 ± 0.01 | 1.35 ± 0.01 | 1.38 ± 0.01 | 1.35 ± 0.01 |
| $C\ldots O_{R}$ | **1.44/2.58** | 1.54 ± 0.01 | 1.49 ± 0.01 | 1.42 ± 0.01 | 1.58 ± 0.02 | 2.63 ± 0.03 |
| $O_{nuc}\ldots H_{nuc}$ | - | 0.99 ± 0.01 | 1.21 ± 0.01 | 1.48 ± 0.02 | 2.50 ± 0.03 | 3.32 ± 0.06 |
| $H_{nuc}\ldots O_{Asp}$ | - | 1.60 ± 0.02 | 1.12 ± 0.01 | 0.99 ± 0.01 | 1.02 ± 0.01 | 1.58 ± 0.02 |
| $H_{Asp}\ldots O_{R}$ | - | 3.21 ± 0.06 | 2.82 ± 0.03 | 2.78 ± 0.04 | 1.33 ± 0.02 | 1.00 ± 0.01 |
| $O_{Tyr}\ldots O_{R}$ | **3.51/3.60** | 4.27 ± 0.11 | 4.40 ± 0.16 | 4.36 ± 0.13 | 4.29 ± 0.12 | 4.79 ± 0.14 |
| $O_{Tyr}\ldots O$ | **2.87/3.50** | 4.74 ± 0.11 | 5.01 ± 0.16 | 5.13 ± 0.13 | 5.59 ± 0.12 | 5.96 ± 0.14 |
| **Concerted Mechanism** | | | | | | |
| $O_{nuc}\ldots C$ | - | 3.03 ± 0.02 | 1.72 ± 0.01 | - | - | 1.27 ± 0.00 |
| $C\ldots O_{R}$ | **1.44/2.58** | 1.28 ± 0.00 | 1.63 ± 0.01 | - | - | 2.52 ± 0.01 |
| $O_{nuc}\ldots H_{nuc}$ | - | 0.97 ± 0.00 | 1.08 ± 0.00 | - | - | 2.05 ± 0.04 |
| $H_{nuc}\ldots O_{R}$ | - | 2.52 ± 0.03 | 1.26 ± 0.01 | - | - | 0.97 ± 0.00 |

| | | | | | | |
|---|---|---|---|---|---|---|
| $O_{Tyr}...O_R$ | **3.51/3.60** | 6.31 ± 0.07 | 6.40 ± 0.07 | - | - | 6.49 ± 0.07 |
| $O_{Tyr}...O_{nuc}$ | **2.87/3.50** | 4.07 ± 0.08 | 4.05 ± 0.08 | - | - | 3.87 ± 0.08 |

[a] All distances are averages and standard error of the mean over 30 independent EVB trajectories, as described in the **Materials and Methods**. The different mechanisms presented in this table are summarized in **Figure 1** of the main text. RS, TS1, IS, TS2 and PS denote the Michaelis complex, transition state for nucleophilic attack on the lactone, tetrahedral intermediate, transition state for ring opening and breakdown of the tetrahedral intermediate (in the case of the first three mechanisms, all of which involve an intermediate) and the final product complex. $O_{nuc}...C$ and $C...O_R$ denote the carbon-nucleophile and carbon-ring oxygen distances, respectively. $O_{nuc}$-$H_{nuc}$, $H_{nuc}...O_{Asp}$, $H_{Asp}...O_R$, and $H_{nuc}...O_R$ denote proton transfer distances (in the Asp and concerted mechanisms) between the nucleophile oxygen and proton being transferred, the proton being transferred and the side-chain oxygen of the aspartic acid, the proton being transferred (step 2, Asp mechanism) from the aspartic acid side-chain to the ring oxygen, and the proton being transferred from the nucleophilic water molecule to the ring oxygen (concerted mechanism), respectively. $O_{Tyr}...O_R$, $O_{Tyr}...O$ and $O_{Tyr}...O_{nuc}$ denote distances between Tyr223 oxygen and the carbon-ring oxygen, the lactone carboxyl oxygen and the oxygen of the nucleophilic water (concerted mechanism), respectively. The data for the corresponding non-enzymatic reaction is shown in **Table S3**.

**Table S3**. Reacting distances (Å) at key stationary points obtained from EVB simulations of the non-enzymatic hydrolysis of C6-HSL.[a]

| Distance | RS | TS1 | IS | TS2 | PS |
|---|---|---|---|---|---|
| **Bridging Hydroxide Mechanism** | | | | | |
| $O_{nuc}$…C | 4.04 ± 0.03 | 1.51 ± 0.00 | 1.34 ± 0.00 | 1.37 ± 0.00 | 1.36 ± 0.00 |
| C…$O_R$ | 1.53 ± 0.00 | 1.49 ± 0.00 | 1.40 ± 0.00 | 1.56 ± 0.00 | 2.61 ± 0.01 |
| **Terminal Hydroxide Mechanism** | | | | | |
| $O_{nuc}$…C | 4.04 ± 0.03 | 1.51 ± 0.00 | 1.34 ± 0.00 | 1.32 ± 0.00 | 1.25 ± 0.00 |
| C…$O_R$ | 1.53 ± 0.00 | 1.49 ± 0.00 | 1.40 ± 0.00 | 1.53 ± 0.00 | 2.55 ± 0.01 |
| **Asp Mechanism** | | | | | |
| $O_{nuc}$…C | 2.75 ± 0.05 | 1.51 ± 0.01 | 1.35 ± 0.01 | 1.40 ± 0.01 | 1.35 ± 0.01 |
| C…$O_R$ | 1.53 ± 0.01 | 1.48 ± 0.01 | 1.40 ± 0.01 | 1.59 ± 0.01 | 2.47 ± 0.03 |
| $O_{nuc}$…$H_{nuc}$ | 0.99 ± 0.01 | 1.23 ± 0.01 | 1.57 ± 0.02 | 2.66 ± 0.03 | 3.13 ± 0.05 |
| $H_{nuc}$…$O_{Asp}$ | 1.62 ± 0.03 | 1.12 ± 0.01 | 0.97 ± 0.01 | 1.06 ± 0.01 | 1.60 ± 0.02 |
| $H_{Asp}$…$O_R$ | 3.57 ± 0.06 | 2.89 ± 0.04 | 2.64 ± 0.04 | 1.26 ± 0.01 | 0.98 ± 0.01 |
| **Concerted Mechanism** | | | | | |
| $O_{nuc}$…C | 3.05±0.01 | 1.69±0.01 | - | - | 1.28±0.00 |
| C…$O_R$ | 1.29±0.00 | 1.67±0.00 | - | - | 2.62±0.01 |
| $O_{nuc}$…$H_{nuc}$ | 0.98±0.00 | 1.12±0.00 | - | - | 1.63±0.01 |
| $H_{nuc}$…$O_R$ | 1.68±0.01 | 1.21±0.00 | - | - | 0.97±0.00 |

[a] All distances are averages and standard error of the mean over 30 independent EVB trajectories, as described in the **Materials and Methods**. The different mechanisms presented in this table are summarized in **Figure 1** of the main text. RS, TS1, IS, TS2 and PS denote the reactant state, transition state for nucleophilic attack on the lactone, tetrahedral intermediate, transition state for ring opening and breakdown of the tetrahedral intermediate (in the case of the first three mechanisms, all of which involve an intermediate) and the final product state. $O_{nuc}$…C and C…$O_R$ denote the carbon-nucleophile and carbon-ring oxygen distances, respectively. $O_{nuc}$-$H_{nuc}$, $H_{nuc}$…$O_{Asp}$, $H_{Asp}$…$O_R$, and $H_{nuc}$…$O_R$ denote proton transfer distances (in the Asp and concerted mechanisms) between the nucleophile oxygen and proton being transferred, the proton being transferred and the side-chain oxygen of the propionic acid used as a model for the enzymatic aspartic acid side-chain, the proton being transferred (step 2,

Asp mechanism) from the propionic acid to the ring oxygen, and the proton being transferred from the nucleophilic water molecule to the ring oxygen (concerted mechanism), respectively. The data for the corresponding GcL-catalyzed reaction is shown in **Table S2**.

**Table S4**. Metal-metal and metal-ligand distances (Å) at key stationary points obtained from EVB simulations of the hydrolysis of C6-HSL by wild-type GcL.[a]

| Distance | RS | TS1 | IS | TS2 | PS |
|---|---|---|---|---|---|
| **Bridging Hydroxide Mechanism** | | | | | |
| Fe…Co | 3.28 ± 0.01 | 3.47 ± 0.01 | 3.61 ± 0.01 | 3.73 ± 0.01 | 3.38 ± 0.01 |
| Co…$O_C$ | 2.10 ± 0.00 | 2.06 ± 0.00 | 2.05 ± 0.00 | 2.06 ± 0.00 | 2.07 ± 0.00 |
| Fe…$O_R$ | 2.30 ± 0.01 | 2.18 ± 0.01 | 2.18 ± 0.01 | 2.10 ± 0.00 | 2.11 ± 0.00 |
| Co…$O_{nuc}$ | 2.04 ± 0.00 | 2.07 ± 0.00 | 2.06 ± 0.00 | 2.06 ± 0.00 | 2.11 ± 0.01 |
| Fe…$O_{nuc}$ | 2.10 ± 0.00 | 2.50 ± 0.01 | 2.82 ± 0.01 | 3.05 ± 0.01 | 3.11 ± 0.01 |
| **Terminal Hydroxide Mechanism** | | | | | |
| Fe…Co | 3.19 ± 0.01 | 3.16 ± 0.01 | 3.13 ± 0.01 | 3.16 ± 0.01 | 3.17 ± 0.01 |
| Co…$O_C$ | 2.09 ± 0.00 | 2.09 ± 0.00 | 2.09 ± 0.00 | 2.07 ± 0.00 | 2.05 ± 0.00 |
| Fe…$O_R$ | 4.63 ± 0.01 | 4.34 ± 0.01 | 4.08 ± 0.01 | 4.24 ± 0.01 | 4.65 ± 0.01 |
| Fe…$O_{nuc}$ | 2.07 ± 0.00 | 2.10 ± 0.00 | 2.14 ± 0.00 | 2.15 ± 0.00 | 2.16 ± 0.00 |
| **Asp Mechanism** | | | | | |
| Fe…Co | 3.29 ± 0.03 | 3.19 ± 0.03 | 3.17 ± 0.03 | 3.20 ± 0.03 | 3.24 ± 0.03 |
| Co…$O_C$ | 2.10 ± 0.02 | 2.10 ± 0.02 | 2.09 ± 0.02 | 2.10 ± 0.02 | 2.11 ± 0.02 |
| Fe…$O_R$ | 2.30 ± 0.03 | 2.20 ± 0.02 | 2.16 ± 0.02 | 2.44 ± 0.04 | 2.21 ± 0.02 |
| Fe…$O_{Asp}$ | 2.04 ± 0.02 | 2.10 ± 0.02 | 2.17 ± 0.02 | 2.12 ± 0.02 | 2.05 ± 0.02 |
| **Concerted Mechanism** | | | | | |
| Fe…Co | 3.30±0.01 | 3.49±0.02 | - | - | 3.51±0.02 |
| Co…$O_C$ | 2.65±0.05 | 2.53±0.04 | - | - | 2.44±0.03 |
| Fe…$O_R$ | 3.10±0.06 | 2.95±0.05 | - | - | 2.61±0.04 |

[a] All distances are averages and standard error of the mean over 30 independent EVB trajectories, as described in the **Materials and Methods**. The different mechanisms presented in this table are summarized in **Figure 1** of the main text. MC, TS1, IS, TS2 and PS denote the Michaelis complex, transition state for nucleophilic attack on the lactone, tetrahedral intermediate, transition state for ring opening and breakdown of the tetrahedral intermediate (in the case of the first three mechanisms, all of which involve an intermediate) and the final product complex.

$O_C$ denotes the carbonyl oxygen of the lactone, $O_R$ denotes the lactone ring oxygen, $O_{nuc}$ denotes the oxygen atom of the nucleophile, and $O_{Asp}$ denotes the metal coordinating oxygen of the catalytic aspartic acid side-chain.

**Table S5.** Individual activation and reaction free energies (kcal mol$^{-1}$) for the GcL catalyzed hydrolysis of a range of AHLs, proceeding *via* the terminal hydroxide mechanism.[a]

| Enzyme | Substrate | $\Delta G^{\ddagger}_{exp}$ | $\Delta G^{\ddagger}_{TS1}$ | $\Delta G^{0}_{IS}$ | $\Delta G^{\ddagger}_{TS2}$ | $\Delta G^{0}_{PS}$ |
|---|---|---|---|---|---|---|
| **WT** | C4 | 15.7 | **16.9 ± 0.5** | 11.9 ± 0.6 | 14.1 ± 0.9 | -0.2 ± 1.1 |
| | C6 | 16.2 | **16.3 ± 0.5** | 11.7 ± 0.6 | 13.8 ± 0.8 | 0.9 ± 1.1 |
| | C10 | 16.5 | **16.2 ± 0.6** | 10.5 ± 0.7 | 12.3 ± 1.0 | -4.8 ± 1.1 |
| **D122N** | C4 | 17.1 | 23.9 ± 0.8 | 21.4 ± 0.9 | **30.0 ± 1.5** | 12.6 ± 1.6 |
| | C6 | 16.9 | 29.4 ± 0.7 | 27.5 ± 0.6 | **30.8 ± 1.1** | 19.2 ± 1.1 |
| | C10 | 16.8 | 25.4 ± 0.5 | 24.5 ± 0.6 | **26.2 ± 0.9** | 8.8 ± 1.0 |
| **G156P** | C4 | 16.2 | **15.6 ± 0.5** | 11.9 ± 0.6 | 14.4 ± 1.0 | 0.8 ± 1.1 |
| | C6 | 16.9 | **15.6 ± 0.6** | 12.2 ± 0.8 | 14.7 ± 1.1 | 1.7 ± 1.2 |
| | C10 | 17.7 | **16.3 ± 0.6** | 10.9 ± 0.8 | 12.5 ± 1.1 | -4.9 ± 1.2 |
| **A157G** | C4 | 15.0 | **17.0 ± 0.6** | 13.2 ± 0.6 | 15.3 ± 0.9 | 0.9 ± 1.0 |
| | C6 | 16.1 | **17.0 ± 0.5** | 13.4 ± 0.6 | 15.6 ± 0.9 | 1.1 ± 1.1 |
| | C10 | 16.7 | **17.1 ± 0.5** | 11.8 ± 0.5 | 13.9 ± 0.8 | -3.9 ± 0.9 |
| **A157S** | C4 | 15.6 | **15.3 ± 0.6** | 10.3 ± 0.7 | 13.8 ± 0.9 | -1.6 ± 1.1 |
| | C6 | 16.9 | **16.4 ± 0.5** | 12.6 ± 0.6 | 15.4 ± 1.0 | 2.8 ± 1.0 |
| | C10 | 16.8 | **17.1 ± 0.7** | 11.0 ± 0.8 | 13.2 ± 1.1 | -4.6 ± 1.1 |
| **I237M** | C4 | 16.2 | **16.9 ± 0.5** | 11.9 ± 0.6 | 14.0 ± 0.8 | -0.6 ± 1.1 |
| | C6 | 17.5 | **16.6 ± 0.5** | 12.7 ± 0.6 | 16.2 ± 1.0 | 1.5 ± 1.1 |
| | C10 | 17.4 | **16.7 ± 0.5** | 10.8 ± 0.7 | 13.4 ± 1.1 | -4.9 ± 1.2 |
| **Y223F** | C4 | 17.6 | **15.9 ± 0.5** | 11.3 ± 0.5 | 13.4 ± 0.7 | -1.0 ± 0.8 |
| | C6 | 16.5 | **17.2 ± 0.5** | 13.5 ± 0.5 | 15.7 ± 0.8 | 2.1 ± 0.9 |
| | C10 | 17.4 | **17.5 ± 0.7** | 12.5 ± 0.9 | 14.8 ± 1.4 | -3.2 ± 1.3 |

[a] Experimental values are based on turnover numbers presented in **Table 1**, and calculated values are averages and standard error of the mean over 30 individual EVB trajectories, obtained as described in the **Materials and Methods**. For a description of the terminal hydroxide mechanism, see **Figure 1B** of the main text. The subscripts TS1, IS, TS2 and PS denote the transition state for nucleophilic attack on the lactone, the resulting intermediate, the transition state for ring opening / breakdown of the tetrahedral intermediate, and the final product state. Note that, in the case of this mechanism, a 2.6 kcal mol$^{-1}$ correction has been added to the energies of all steps to take into account the energetic cost of generating a metal bound hydroxide nucleophile, as described in the main text.

**Table S6.** Individual activation and reaction free energies (kcal mol$^{-1}$) for the GcL catalyzed hydrolysis of a range of AHLs, proceeding *via* the Asp mechanism.[a]

| Enzyme | Substrate | $\Delta G^{\ddagger}_{exp}$ | $\Delta G^{\ddagger}_{TS1}$ | $\Delta G^{0}_{IS}$ | $\Delta G^{\ddagger}_{TS2}$ | $\Delta G^{0}_{PS}$ |
|---|---|---|---|---|---|---|
| **WT** | C4 | 15.7 | 15.1 ± 0.2 | 8.5 ± 0.5 | **16.0 ± 0.8** | -4.4 ± 1.3 |
| | C6 | 16.2 | 15.8 ± 0.2 | 11.7 ± 0.5 | **16.2 ± 0.8** | -4.5 ± 0.7 |
| | C10 | 16.5 | 13.8 ± 0.2 | 4.3 ± 0.4 | **15.6 ± 0.9** | -12.1± 1.0 |
| **G156P** | C4 | 16.2 | 13.3 ± 0.4 | 6.9 ± 0.4 | **14.5 ±0.7** | -4.7 ± 1.1 |
| | C6 | 16.9 | 16.6 ± 0.3 | 13.0 ± 0.5 | **17.5 ± 0.8** | -3.1 ± 1.1 |
| | C10 | 17.7 | 14.7 ± 0.3 | 8.5 ± 0.5 | **16.9 ± 1.0** | -7.3 ± 1.5 |
| **A157G** | C4 | 15.0 | 14.5 ± 0.3 | 7.6 ± 0.5 | **16.0 ± 0.8** | -2.9 ± 1.5 |
| | C6 | 16.1 | 17.5 ± 0.3 | 13.9 ± 0.4 | **18.6 ± 0.6** | -2.5 ± 1.1 |
| | C10 | 16.7 | 14.9 ± 0.3 | 8.1 ± 0.6 | **18.2 ± 1.0** | -11.7 ± 1.2 |
| **A157S** | C4 | 15.6 | 14.2 ± 0.4 | 6.9 ± 0.7 | **15.0 ± 1.1** | -4.4 ± 1.3 |
| | C6 | 16.9 | 17.2 ± 0.4 | 13.6 ± 0.5 | **17.9 ± 0.8** | -3.1 ± 1.2 |
| | C10 | 16.8 | 14.9 ± 0.3 | 8.1 ± 0.6 | **17.7 ± 1.0** | -11.4 ± 1.4 |
| **I237M** | C4 | 16.2 | 13.8 ± 0.3 | 6.6 ± 0.5 | **14.6 ± 0.7** | -5.3 ± 0.9 |
| | C6 | 17.5 | 16.7 ± 0.3 | 12.3 ± 0.5 | **17.6 ± 0.8** | -4.0 ± 0.9 |
| | C10 | 17.4 | 14.6 ± 0.3 | 7.2 ± 0.5 | **17.3 ± 0.9** | -12.9 ± 1.0 |
| **Y223F** | C4 | 17.6 | 14.9 ± 0.4 | 7.5 ± 1.0 | **16.3 ± 1.5** | -3.9 ± 2.0 |
| | C6 | 16.5 | 15.6 ± 0.4 | 9.5 ± 0.7 | **15.9 ± 1.1** | -4.9 ± 1.4 |
| | C10 | 17.4 | 13.9 ± 0.5 | 5.3 ± 0.8 | **17.0 ± 1.3** | -13.5 ± 1.5 |

[a] Experimental values are based on turnover numbers presented in **Table 1**, and calculated values are averages and standard error of the mean over 30 individual EVB trajectories, obtained as described in the **Materials and Methods**. For a description of the Asp mechanism, see **Figure 1C** of the main text. The subscripts TS1, IS, TS2 and PS denote the transition state for nucleophilic attack on the lactone, the resulting intermediate, the transition state for ring opening / breakdown of the tetrahedral intermediate, and the final product state.

**Table S7.** Individual activation and reaction free energies (kcal mol$^{-1}$) for the GcL catalyzed hydrolysis of a range of AHLs, proceeding *via* the concerted mechanism.[a]

| Enzyme | Substrate | $\Delta G^{\ddagger}_{exp}$ | $\Delta G^{\ddagger}_{TS}$ | $\Delta G^{0}_{PS}$ |
|---|---|---|---|---|
| **WT** | C4 | 15.7 | 22.2 ± 0.3 | -1.3 ± 0.4 |
| | C6 | 16.2 | 21.1 ± 0.4 | -3.8 ± 0.5 |
| | C10 | 16.5 | 20.8 ± 0.3 | -6.9 ± 0.6 |
| **D122N** | C4 | 17.1 | 19.0 ± 0.3 | -10.8 ± 0.5 |
| | C6 | 16.9 | 19.4 ± 0.3 | -9.4 ± 0.4 |
| | C10 | 16.8 | 19.6 ± 0.6 | -11.0 ± 1.0 |
| **G156P** | C4 | 16.2 | 21.6 ± 0.4 | -2.8 ± 0.7 |
| | C6 | 16.9 | 21.5 ± 0.4 | -2.3 ± 0.7 |
| | C10 | 17.7 | 20.9 ± 0.4 | -3.6 ± 0.5 |
| **A157G** | C4 | 15.0 | 21.6 ± 0.3 | -2.7 ± 0.4 |
| | C6 | 16.1 | 23.0 ± 0.4 | -1.7 ± 0.6 |
| | C10 | 16.7 | 22.2 ± 0.5 | -4.2 ± 0.4 |
| **A157S** | C4 | 15.6 | 21.5 ± 0.3 | -2.4 ± 0.5 |
| | C6 | 16.9 | 21.6 ± 0.3 | -1.4 ± 0.5 |
| | C10 | 16.8 | 23.6 ± 0.4 | 0.6 ± 0.5 |
| **I237M** | C4 | 16.2 | 21.4 ± 0.4 | -2.9 ± 0.6 |
| | C6 | 17.5 | 21.4 ± 0.4 | -3.0 ± 0.5 |
| | C10 | 17.4 | 20.5 ± 0.4 | -4.6 ± 0.6 |
| **Y223F** | C4 | 17.6 | 21.6 ± 0.2 | -2.0 ± 0.4 |
| | C6 | 16.5 | 22.5 ± 0.3 | -1.6 ± 0.5 |
| | C10 | 17.4 | 21.1 ± 0.5 | -4.7 ± 0.8 |

[a] Experimental values are based on turnover numbers presented in **Table 1**, and calculated values are averages and standard error of the mean over 30 individual EVB trajectories, obtained as described in the **Materials and Methods**. For a description of the concerted mechanism, see **Figure 2** of the main text. The subscripts TS and PS denote the concerted transition state for nucleophilic attack on the lactone and opening of the lactone ring, and the final product state.

**Table S8.** A Comparison of experimental and calculated activation free energies (kcal mol$^{-1}$) for the GcL catalyzed hydrolysis of a range of AHLs.[a]

| Enzyme | Substrate | $\Delta G^{\ddagger}_{exp}$ | $\Delta G^{\ddagger}_{calc,Ter}$ | $\Delta G^{\ddagger}_{calc\rightarrow exp,Ter}$ | $\Delta G^{\ddagger}_{calc,Asp}$ | $\Delta G^{\ddagger}_{calc\rightarrow exp,Asp}$ | $\Delta G^{\ddagger}_{calc,Conc}$ | $\Delta G^{\ddagger}_{calc\rightarrow exp,Conc}$ |
|---|---|---|---|---|---|---|---|---|
| **WT** | C4 | 15.7 | 16.9 ± 0.5 | 1.2 | **16.0 ± 0.8** | 0.3 | 22.2 ± 0.3 | 6.5 |
| | C6 | 16.2 | 16.3 ± 0.5 | 0.1 | **16.2 ± 0.8** | 0.0 | 21.1 ± 0.4 | 4.9 |
| | C10 | 16.5 | 16.2 ± 0.6 | -0.3 | **15.6 ± 0.9** | -0.9 | 20.8 ± 0.3 | 4.3 |
| **D122N** | C4 | 17.1 | 30.0 ± 1.5 | 12.9 | - | - | **19.0 ± 0.3** | 1.9 |
| | C6 | 16.9 | 30.8 ± 1.1 | 13.9 | - | - | **19.4 ± 0.3** | 2.5 |
| | C10 | 16.8 | 26.2 ± 0.9 | 9.4 | - | - | **19.6 ± 0.6** | 2.8 |
| **G156P** | C4 | 16.2 | 15.6 ± 0.5 | -0.6 | **14.5 ± 0.7** | -1.7 | 21.6 ± 0.4 | 5.4 |
| | C6 | 16.9 | **15.6 ± 0.6** | -1.3 | 17.5 ± 0.8 | 0.6 | 21.5 ± 0.4 | 4.6 |
| | C10 | 17.7 | **16.3 ± 0.6** | -1.4 | 16.9 ± 1.0 | -0.8 | 20.9 ± 0.4 | 3.2 |
| **A157G** | C4 | 15.0 | 17.0 ± 0.6 | 2.0 | **16.0 ± 0.8** | 1.0 | 21.6 ± 0.3 | 6.6 |
| | C6 | 16.1 | **17.0 ± 0.5** | 0.9 | 18.6 ± 0.6 | 2.5 | 23.0 ± 0.4 | 6.9 |
| | C10 | 16.7 | **17.1 ± 0.5** | 0.4 | 18.2 ± 1.0 | 1.5 | 22.2 ± 0.5 | 5.5 |
| **A157S** | C4 | 15.6 | 15.3 ± 0.6 | -0.3 | **15.0 ± 1.1** | -0.6 | 21.5 ± 0.3 | 5.9 |
| | C6 | 16.9 | **16.4 ± 0.5** | -0.5 | 17.9 ± 0.8 | 1.0 | 21.6 ± 0.3 | 4.7 |
| | C10 | 16.8 | **17.1 ± 0.7** | 0.3 | 17.7 ± 1.0 | 0.9 | 23.6 ± 0.4 | 6.8 |
| **I237M** | C4 | 16.2 | 16.9 ± 0.5 | 0.7 | **14.6 ± 0.7** | -1.6 | 21.4 ± 0.4 | 5.2 |
| | C6 | 17.5 | **16.6 ± 0.5** | -0.9 | 17.6 ± 0.8 | 0.1 | 21.4 ± 0.4 | 3.9 |
| | C10 | 17.4 | **16.9 ± 0.5** | -0.5 | 17.3 ± 0.9 | -0.1 | 20.5 ± 0.4 | 3.1 |
| **Y223F** | C4 | 17.6 | **15.9 ± 0.5** | -1.7 | 16.3 ± 1.5 | -1.3 | 21.6 ± 0.2 | 4.0 |
| | C6 | 16.5 | **17.2 ± 0.5** | 0.7 | 15.9 ± 1.1 | -0.6 | 22.5 ± 0.3 | 6.0 |
| | C10 | 17.4 | 17.5 ± 0.7 | 0.1 | **17.0 ± 1.3** | -0.4 | 21.1 ± 0.5 | 3.7 |

[a] Experimental values are based on turnover numbers presented in **Table 1**, and calculated values are averages and standard error of the mean over 30 individual EVB trajectories, obtained as described in the **Materials and Methods**. In the case of the terminal hydroxide (Ter) and Asp mechanisms, only the activation free energy for the rate limiting step is shown, which in the case of the terminal hydroxide mechanism is initial nucleophilic attack step and in the case of the Asp mechanism is the second step of the reaction involving ring opening and the breakdown of the tetrahedral intermediate (**Figure 1**, note that in the case of the Asp mechanism the errors shown take propagation of errors into account from previous steps). The concerted mechanism (Conc) is a single-step reaction. The different plausible mechanisms are summarized in **Figure 1** of the main text, and the corresponding activation and reaction free energies for each of the individual reactions presented here are shown in **Tables S7 - S9** for the terminal hydroxide, Asp and concerted mechanisms, respectively.

**Table S9.** Hydrogen bonding interactions between the hydroxide group of Tyr223 and the AHL amide moiety of C4-, C6- and C8-HSL, bound to wild-type GcL, as well as the I237M and G156P GcL variants.[a]

| **WT** | **C4** | **C6** | **C8** |
|---|---|---|---|
| **Tyr-OH – AHL-NH** | 22% | 43% | 4% |
| **Tyr-OH – AHL-O** | - | - | 18% |
| **I237M** | **C4** | **C6** | **C8** |
| **Tyr-OH – AHL-NH** | 13% | 14% | 28% |
| **Tyr-OH – AHL-O** | 2% | 4% | 3% |
| **G156P** | **C4** | **C6** | **C8** |
| **Tyr-OH – AHL-NH** | 21% | 51% | 47% |
| **Tyr-OH – AHL-O** | - | - | - |

[a] Data correspond to the % of the simulated time where the interaction is present, across three independent replicas of 500 ns length each for each system.

**Table S10.** Experimental kinetic parameters for C6-HSL hydrolysis by wild-type AiiA and AaL.

| **Enzyme** | $k_{cat}$ ($s^{-1}$) | $K_M$ (mM) | $k_{cat}/K_M$ ($s^{-1}$ $M^{-1}$) | $\Delta G^{\ddagger}_{exp}$ (kcal $mol^{-1}$) |
|---|---|---|---|---|
| **AiiA**[a] | 91.00 ± 3.00 | 5.60 ± 0.60 | $1.6 \times 10^4$ | 14.8 |
| **AaL**[b] | 13.97 ± 0.43 | 0.08 ± 0.01 | $1.7 \times 10^5$ | 15.9 |

[a] Taken from experimental data presented in ref. [30]. [b] Taken from experimental data presented in ref. [31].

**Table S11**: Table of primers.

| **Mutation** | **Forward** | **Reverse** |
|---|---|---|
| **D122N** | 5'-CAC ATC TGC ACC TGA ACC ATG CGG GTT GC-3' | 5'-GCA ACC CGC ATG GTT CAG GTG CAG ATG TG-3' |
| **Y223F** | 5'AGT GAT GCC ATC TTT ACG G CC GAA-3' | 5'TTC GGC CGT AAA GAT GGC ATC ACT-3' |

**Table S12**. Protonation states of ionizable residues, as well as the protonation patterns of histidine residues, during the empirical valence bond (EVB) simulations.[a]

| Type | Residue Numbers | | |
|---|---|---|---|
| | GcL | AaL | AiiA |
| **Asp** | Chain A: 16, 23, 65, 122[b], 140, 162, 164, 220[b], 240, 267<br>Chain B: 240 | Chain A: 8, 15, 48, 57, 63, 114[b], 132, 154, 156, 173, 212[b], 232, 259<br>Chain B: 295, 512 | 17, 50, 81, 95, 108[b], 154, 191[b], 201, 209, 236 |
| **Glu** | Chain A: 44, 47, 81, 93, 94, 128, 139, 141, 155, 180, 208, 226, 249, 269<br>Chain B: 226 | Chain A: 36, 39, 86, 120, 131, 133, 147, 172, 218, 241, 253, 261<br>Chain B: 498 | 41, 55, 61, 70, 79, 80, 127, 129, 135, 136, 140, 152, 156, 179, 197, 200, 202, 211, 222, 227, 238, 240, 248 |
| **Lys** | Chain A: 24, 154, 168, 177, 190, 233, 272, 273, 276<br>Chain B: 24, 233 | Chain A: 16, 53, 160, 169, 182, 204, 225, 242, 244, 264, 268<br>Chain B: 296, 505 | 29, 76, 88, 139, 149, 196, 218, 221, 225, 226, 241 |
| **Arg** | Chain A: 19, 21, 78, 100, 151, 174, 178, 250, 252, 253 | Chain A: 11, 13, 70, 92, 143, 166, 170, 238, 245, 265, 267<br>Chain B: 518 | 13, 82, 89, 125, 134, 219, 244 |
| **His-δ** | Both Chains: 31, 98, 118, 123, 179, 198, 266 | Both Chains: 23, 90, 110, 115, 190, 198, 258 | 104, 109, 133, 145, 169, 173, 235 |
| **His-ε** | Both Chains: 37, 57, 120, 138, 206 | Both Chains: 49, 112, 130 | 18, 106 |

[a] The relevant protein chain is indicated. All residues not included in this table were kept in their unionized forms as they were located outside the simulation sphere (see the **Materials and Methods** for further details, this is why only a few residues from Chain B were kept in their ionized forms, as they fell within the simulation sphere). [b] The charges of these aspartic acid side-chains were modified according to cluster QM calculations in order to maintain the stability of the metal coordination spheres, and renamed as ASX (see **Table S13** for the parameters).

**Table S13.** Quantum chemically calculated charges for the ASX side-chain.

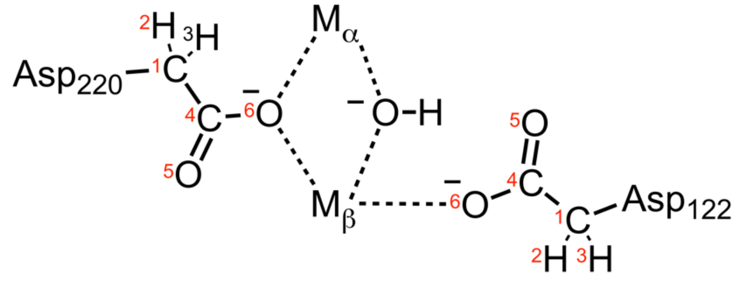


| Index | Atom name | Charge |
|---|---|---|
| 1 | CB | -0.2200 |
| 2 | HB2 | 0.0600 |
| 3 | HB3 | 0.0600 |
| 4 | CG | 0.7000 |
| 5 | OD1 | -0.6576 |
| 6 | OD2 | -0.9424 |

**Table S14.** EVB parameters used to describe the hydrolysis of a range of AHL substrates by the lactonases studied in this work. [a]

| Substrate | Reaction Step | Bridging $OH^-$ | Terminal $OH^-$ | Asp | Concerted |
|---|---|---|---|---|---|
| **C4-HSL** | 1 | - | $H_{ij}$: 21.0, $\alpha_{ij}$: 64.1 | $H_{ij}$: 47.0, $\alpha_{ij}$: 111.0 | $H_{ij}$: 72.5, $\alpha_{ij}$: 17.6 |
| | 2 | - | $H_{ij}$: 86.4, $\alpha_{ij}$: -110.1 | $H_{ij}$: 62.5, $\alpha_{ij}$: -99.8 | - |
| **C6-HSL** | 1 | $H_{ij}$: 21.2, $\alpha_{ij}$:78.3 | $H_{ij}$: 21.4, $\alpha_{ij}$: 71.1 | $H_{ij}$: 49.1 $a_{ij}$: 107.9 | $H_{ij}$: 72.6 $a_{ij}$: 19.0 |
| | 2 | $H_{ij}$:43.8, $\alpha_{ij}$: -93.3 | $H_{ij}$: 91.8, $\alpha_{ij}$: -144.3 | $H_{ij}$: 66.3, $\alpha_{ij}$: -93.7 | - |
| **C10-HSL** | 1 | - | $H_{ij}$: 21.9, $\alpha_{ij}$: 65.0 | $H_{ij}$: 47.8, $\alpha_{ij}$: 98.9 | $H_{ij}$: 71.8, $\alpha_{ij}$: 19.2 |
| | 2 | - | $H_{ij}$: 88.1, $\alpha_{ij}$: -99.0 | $H_{ij}$: 62.4, $\alpha_{ij}$: -85.9 | - |

[a] The EVB off-diagonal term, $H_{ij}$, and gas-phase shift, $\alpha_i$, were fit to reproduce Activation free energies for the hydrolysis of each AHL substrate *via* each mechanism considered here, as described in the **Supplementary Methodology**. The different mechanisms presented in this table are summarized in **Figure 1** of the main text. In the case of multi-step mechanisms, each reaction step was calibrated separately. The same parameters were then used unchanged across all enzymes and enzyme variants studied in this work. For a detailed description of the physical meaning of these parameters, see e.g. refs. [10, 11].
